# Feedback from the Nascent Chain Triggers Ribosomal Frameshifting and Transcript Decay

**DOI:** 10.64898/2025.12.23.696290

**Authors:** Patrick J. Carmody, Caden R. Sillman, Dyotima, Rohan Bhardwaj, Ali Farzam, Morvarid Golrokhmofrad, Braden Lewis, Wesley D. Penn, Bryon S. Drown, Charles P. Kuntz, Jonathan P. Schlebach

**Author notes:** Contributed Equally (Patrick J. Carmody, Caden R. Sillman, Dyotima).

## Abstract

Though ribosomes have several features that help them maintain their reading frame, these safeguards can be bypassed by RNA structures that promote −1 programmed ribosomal frameshifting (-1PRF). We recently found that conformational transitions in the nascent polypeptide can enhance -1PRF, though it’s unclear whether this feedback plays a general role in translational recoding. Here we demonstrate that the translocation of nascent transmembrane domains is sufficient to induce -1PRF during the decoding of slippery heptamers. We identify thousands of motifs that potentially trigger -1PRF along with proteomic identifications of 33 predicted human frameshift products. We also identify thousands of splicing-dependent motifs and demonstrate that the splicing-mediated reconfiguration of transmembrane domains alters -1PRF. Finally, we show that most transcripts bearing these motifs are sensitive to the nonsense-mediated decay regulator UPF1, suggesting they modulate mRNA turnover. Our findings show that the misassembly of growing polypeptides can trigger -1PRF, premature termination, and transcript decay.

## Introduction

Ribosomes cooperate with a variety of biosynthetic machinery to translate the genetic code and target nascent polypeptides to various cellular microenvironments. This translation process is kinetically controlled by a variety of cofactors and GTPases that collectively ensure protein synthesis proceeds from a contiguous reading frame in a manner that is synchronized with cotranslational protein assembly. However, there are certain coding RNA that contain structured regions that allow the ribosome to bypass this kinetic control and shift into an alternative reading frame.^1–3^ This translational recoding mechanism, which is known as −1 programmed ribosomal frameshifting (-1PRF), has been predominantly investigated in viruses, where it regulates translational stoichiometry and generates proteins encoded in the −1 reading frame.^4^ By comparison, there are relatively few known instances of functional -1PRF in eukaryotic systems.^5^ Nevertheless, emerging evidence suggests ribosomal frameshifting in higher organisms may play a role in transcriptome regulation,^6^ metabolic adaptation,^6–8^ and protein quality control.^9^ Notably, these regulatory events appear to respond to diverse molecular inputs, which suggests they may potentially involve distinct recoding mechanisms.

-1PRF sites typically feature specialized RNA structures that work in concert with various translation factors to modulate translational recoding.^10^ Ribosomal frameshifting most readily occurs during the decoding of certain “slippery” X_1_ XXY_4_ YYZ_7_ heptanucleotide sequences,^11,12^ where X, Y, and Z represent arbitrary nucleobases. This structure enables the anticodon loops of the P-site and A-site tRNAs to form isoenergetic base pairing interactions with alternative codons in the −1 reading frame.^13^ To override the kinetic barriers that typically prevent such frameshifts, -1PRF sites typically also contain an adjacent mRNA secondary structure that slows ribosomal elongation in the 0-frame relative to the −1 frame.^14–16^ Similar delays in translocation can also arise from the depletion of A-site tRNAs and/ or the association of various effector molecules with the transcript and/ or ribosome.^17,18^ Together, these factors provide the timing needed for the P-site and A-site tRNA(s) to shift into the −1 reading frame and then continue translation of a frameshifted polypeptide.

We recently found that the activity of the -1PRF motif within the Sindbis virus genome is mechanically coupled to conformational transitions in the nascent polypeptide.^4,19,20^ Our findings revealed that the efficiency of frameshifting is maximized when the decoding of the slippery sequence coincides with the translocon-mediated membrane integration of a nascent transmembrane domain (TMD). This translocation generates mechanical tension in the nascent chain, which we found to be statistically correlated with the net -1PRF efficiency across a panel of hundreds of sequence variants.^20^ Subsequent investigations of the structural basis of -1PRF in SARS CoV-2 also found that the structure of the nascent chain and its interaction with the ribosome impacts the net -1PRF efficiency.^21^ Together, these findings suggest that translational recoding is more generally coupled to conformational transitions in the nascent chain. Overall, our observations suggest that the mechanochemical fluctuations generated by cotranslational assembly may therefore provide the ribosome with a readout for the folding/ assembly of the nascent chain that can modulate decoding in real time. Nevertheless, the mechanistic basis for this mechanical coupling and the conditions under which this coupling is established remain unclear.

In this work, we explore the coupling between cotranslational folding and ribosomal frameshifting in the context of transcripts encoding integral membrane proteins. By introducing minimalistic changes to a transcript encoding a model membrane protein, we demonstrate that efficient ribosomal frameshifting occurs when the translocation of the nascent chain coincides with the decoding of a slippery heptamer. Based on these observations, we identify thousands of comparable “TMD-Slip” motifs and evaluate their statistical context within the human transcriptome. We find that these motifs are statistically depleted from the transcriptome overall but are over-represented among transcripts encoding receptors, transporters, and ion channels. To validate the activity of these recoding motifs, we identify 33 predicted frameshift products within a recently described deep proteomic dataset.^22^ Finally, we show that these motifs are statistically enriched among transcripts that are degraded by the NMD pathway and that these motifs can be “activated” by alternative splicing. Together, our findings reveal a new class of relatively common eukaryotic -1PRF site that appears to mediate negative translational regulation in response to the misassembly of a wide variety of receptors, transporters, and ion channels.

## Results

### Rational Design of an Artificial Nascent Chain-Mediated PRF Site

Viral -1PRF sites typically feature several regulatory elements that collectively modulate their net ribosomal frameshifting efficiency. Given the diverse mechanisms by which these effectors alter -1PRF, it remains challenging to identify the minimal requirements for coupling between cotranslational folding and ribosomal frameshifting. We therefore sought to identify sequence modifications that generate such coupling in the context of a transcript that lacks a natural ribosomal frameshift motif. For this purpose, we modified a transcript encoding a model membrane protein reporter (LepB) such that a −1 ribosomal frameshift at various positions produces a truncation product with a specific molecular weight (Fig. 1A, Doc. S1). To reduce the complexity of LepB translation products, we removed all but one N-linked glycosylation site. The OST-mediated addition of a ~2.8 kDa sugar at this position serves as a marker for the targeting of the nascent chain to the translocon by the signal recognition particle-a prerequisite for cotranslational folding within the membrane. As expected, *in vitro* translation of this construct in rabbit reticulocyte lysate containing ER membrane fragments (rough microsomes, RM) generates both a high-weight membrane-targeted glycoprotein and a lower weight untargeted and/ or unglycosylated protein (see Lane 3, Fig. 1B). Using this system, the impacts of sequence modifications that appreciably enhance ribosomal frameshifting can be tracked according to changes in the relative abundance of the full-length protein encoded in the 0-frame (FL, 407 residues, ~45 ± 2.8 kDa) and the −1 ribosomal frameshift product (RF, 327 AA, ~36 ± 2.8 kDa).

**Figure 1.**
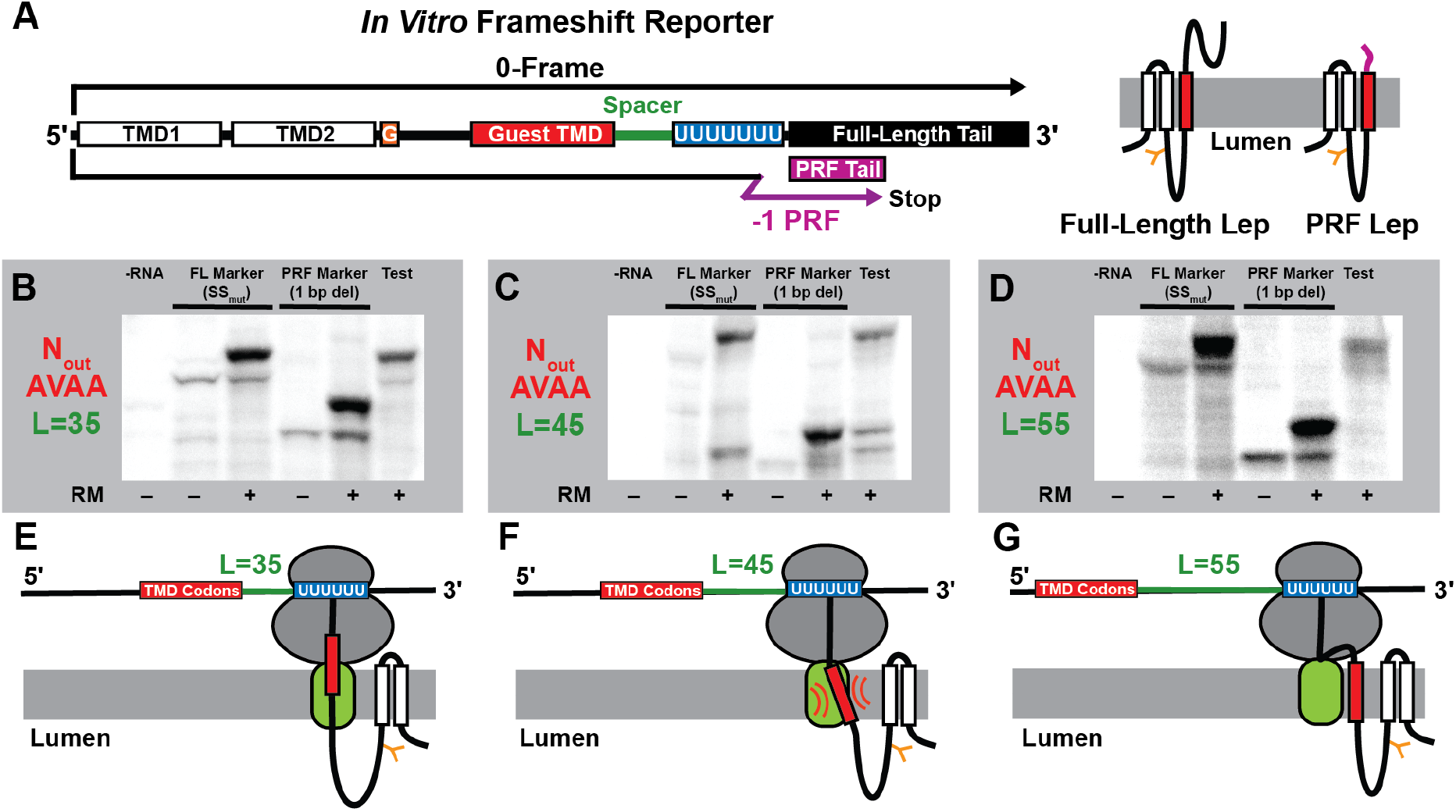
Programmed Ribosomal Frameshifting During *in vitro* Translation of LepB. A) A schematic depicts the relative orientation of the sequence elements in the LepB frameshift reporter constructs including the first two TMDs (white), the glycosylation site (orange), the guest TMD (red), the spacer region (green), the poly-U slip-site (blue), and the tail region encoding the full-length tail in the 0-frame (FL, black) as well as the ribosomal frameshift tail in the −1 frame (RF, purple). Cartoons to the right depict the orientation of the FL and RF products with respect to the membrane. B-D) Representative SDS-PAGE gels depict the translation products generated by LepB frameshift reporter constructs bearing B) 35 codon, C) 45 codon, and D) 55 codon spacer regions between the TMD region and the slippery sequence. Lane 1 in each gel is a negative control containing no RNA. FL molecular weight markers bearing mutations within the slip-site were generated in the absence (lane 2) and presence (lane 3) of rough microsomes. RF molecular weight markers in which the RF tail was moved to the 0-frame were also generated in the absence (lane 4) and presence (lane 5) of microsomes. The test constructs for each spacer length are shown in lane 6 for each gel. E-G) Cartoons depict the topological orientation of the nascent chain during the decoding of the slippery sequence for each linker length.

Our previous investigations revealed that the cotranslational membrane integration of a nascent TMD enhances frameshifting at a -1PRF site containing a heptanucleotide slip-site and an RNA stem-loop. To determine whether membrane translocation alone is sufficient to alter the decoding of slippery sequences, we introduced a slip-site (U_1_ UUU_4_ UUU_7_) at various positions relative to a region encoding a marginally hydrophobic guest TMD (AAAAVAAAAAAAAAVAAAA) (AVAA TMD, Fig. 1A). Translation of a LepB transcript bearing a slippery sequence 45 codons downstream of the AVAA TMD produced a series of products that correspond to the targeted and untargeted versions of the FL and RF products (Fig. 1C). Quantification of the relative abundance of these products suggests this modification triggers highly efficient ribosomal frameshifting (46 ± 8%, Table S1). In contrast, placement of the slip-site at either 35 or 55 codons upstream of the TMD produced no detectable frameshift products (Fig. 1 B & D). This observed distance dependence is consistent with our previous findings in the alphavirus structural polyprotein,^19^ which revealed that the coupling between folding and frameshifting is most pronounced when translocation coincides with the decoding of the slippery sequence (see cartoons in Fig. 1E-G). Notably, energetic predictions suggest any secondary structures within the portions of the transcripts 5-9 bases downstream of these slip-sites are unlikely to be highly stable (Fig. S1). Thus, these results suggest the translocation of the nascent chain is sufficient to enhance the recoding of slippery regions within a transcript.

### Sequence Constraints of Nascent Chain-Mediated Ribosomal Frameshifting

Our previous findings in the alphavirus structural polyprotein suggest the degree to which its nascent chain modulates ribosomal frameshifting is related to the probability that its nascent TMD is engaged by the translocon and partitions into the membrane.^19,20^ Similar to the TMD that enhances frameshifting in alphaviruses, the marginally hydrophobic AVAA TMD is known to be inefficiently recognized by the translocon.^23^ To determine whether -1PRF during LepB translation is sensitive to translocation efficiency, we replaced the AVAA TMD with a more hydrophobic segment that is known to be efficiently recognized by the translocon (ALAALALAALAALALAALA) (LALA TMD).^23^ Consistent with our initial observations, a prominent frameshift product was generated only when a slip-site was introduced 45 codons downstream from the LALA TMD, but not at distances of 35 or 55 codons (Fig. S2). Furthermore, enhancing the hydrophobicity of the TMD increases the observed frameshifting efficiency (52 ± 4%, Table S1), which is again consistent with our observations in alphaviruses. In addition to the distance dependence described above (Fig. 1 E-G), this observed dependence on TMD hydrophobicity provides additional evidence that the interaction of the nascent chain with the translocon stimulates ribosomal frameshifting in LepB.

The magnitude of the mechanical force generated by the translocation of the nascent chain is known to depend on its topological orientation.^24,25^ The N-terminal residues of the TMD that modulates alphavirus -1PRF are projected into the cytosol (N_in_), which is a topological orientation that generates a relatively weak force on the nascent chain.^24^ In contrast, the N-terminal residues of the TMD that modulates LepB frameshifting are projected into the lumen (N_out_), which should generate a greater force on the nascent chain. To determine whether the coupling between folding and frameshifting depends on topology, we introduced an additional upstream TMD into the LepB reporter in order to invert the orientation of the LALA and AVAA TMDs (Fig. 2A). LepB transcripts encoding these topologically modified proteins also efficiently generate frameshifted proteins in this orientation (Fig. 2 B & C). Interestingly, the observed frameshift efficiencies for LepB constructs bearing inverted (N_in_) LALA and AVAA TMDs 45 codons from the slip-site are similar to those observed for the N_out_ topology and appear to be insensitive to TMD hydrophobicity (Table S1, Fig. 2 B & C). Overall these data demonstrate that that the efficiency of the coupling between translocation and ribosomal frameshifting is much more dependent upon the position of the TMD than its hydrophobicity or topological orientation (Table S1).

**Figure 2.**
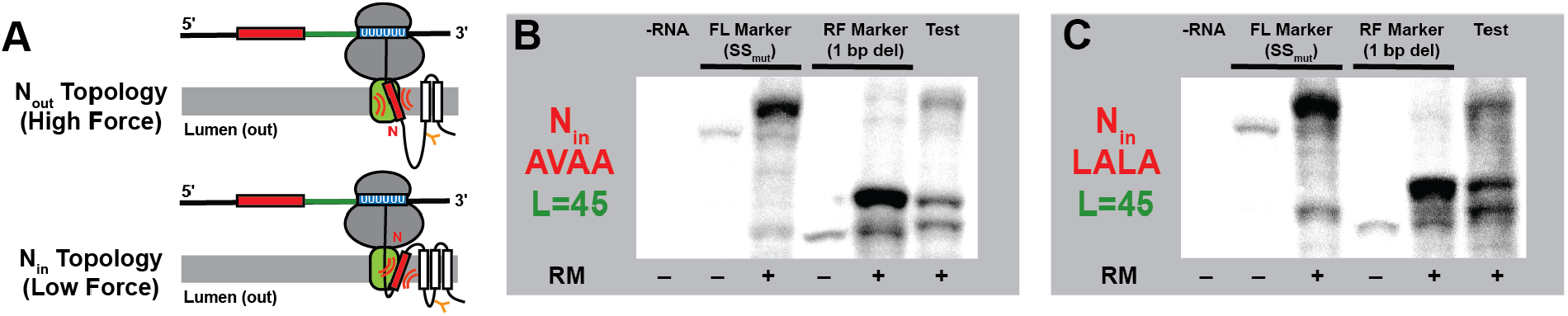
Impact of Topology on Programmed Ribosomal Frameshifting in LepB. A) Cartoons depict the topological orientation of the nascent chain during the decoding of the slippery sequence for LepB constructs bearing three total TMDs in which the guest TMD adopts an N_out_ orientation or four total TMDs in which the guest TMD adopts an N_in_ orientation. B-C) Representative SDS-PAGE gels depict the translation products generated by LepB frameshift reporter constructs bearing B) the AVAA or C) the LALA guest TMDs. Lane 1 in each gel is a negative control containing no RNA. Molecular weight markers for the full-length 0-frame product (FL), which bear mutations within the slip-site, were generated in the absence (lane 2) and presence (lane 3) of rough microsomes. Molecular weight markers for the ribosomal frameshift product (RF), in which the RF tail was moved to the 0-frame, were also generated in the absence (lane 4) and presence (lane 5) of microsomes. The test constructs for each spacer length are shown in lane 6 for each gel.

### LepB Frameshifting in the Cell

Given that ribosomal frameshifting is modulated by translation kinetics and polysome density,^14–16,26^ the observed LepB reporter frameshifting could potentially stem from the artificial time scales and/ or stoichiometries that occur in the context of *in vitro* translation systems. To determine whether these transcripts promote frameshifting in cells, we introduced these LepB constructs into a previously described genetic reporter^19,20^ in which -1PRF generates a GFP signal that can be normalized by the intensity of a second constitutively expressed fluorescent protein (IRES-mKate, Figs. 3A & Doc. S1). Unlike other -1PRF reporters, this design preserves SRP-mediated targeting-a key consideration to ensure cotranslational folding occurs at the ER membrane. We utilized RT-PCR to confirm these reporter constructs remain intact within HEK293T cells (Fig. S3), then measured cellular florescence levels by flow cytometry in order to compare frameshift product intensities within a fixed range of expression. A LepB reporter construct containing a slippery sequence that is optimally spaced (45 codons) relative to the LALA TMD in the N_in_ orientation generates GFP: mKate ratios in HEK293T cells that exceed the signals generated by three different negative controls (Fig. 3A, B, and E). Notably, these measurements were recorded at a consistent expression level and changes in the GFP: mKate ratio primarily reflect changes in GFP intensity, which confirms that these variations reflect differences in translational recoding (Fig. 3C-E). Consistent with biochemical measurements, the signals are less pronounced when slippery sequences instead fall 35 codons downstream of the TMD (Table S1), though elevated ratios were still observed at a 55-codon spacing (Fig. 3E). However, unlike our biochemical results, we find that frameshifting is most efficient when the LALA TMD adopts an N_in_ orientation (Fig. 3C-E, Table S1). This discrepancy could potentially reflect the differential recruitment of various translocon subunits under biochemical and cellular conditions (see *Discussion*).^27–30^ Based on a comparison of the observed GFP: mKate ratios to a control construct in which GFP is placed in the 0-frame, we estimate this construct achieves a PRF efficiency of 3.5 ± 0.5%-several orders of magnitude higher than baseline frameshifting efficiencies (~10^−4^-10^−5^).^31^ Though higher efficiencies were observed *in vitro*, these results confirm that the translocation of the nascent chain alone is sufficient to enhance translational recoding under cellular conditions. Together, our findings suggest the combination of a TMD and an appropriately spaced slippery sequence may constitute a generalized class of ribosomal frameshifting site, which we will refer to hereafter as a TMD-slip motif.

**Figure 3.**
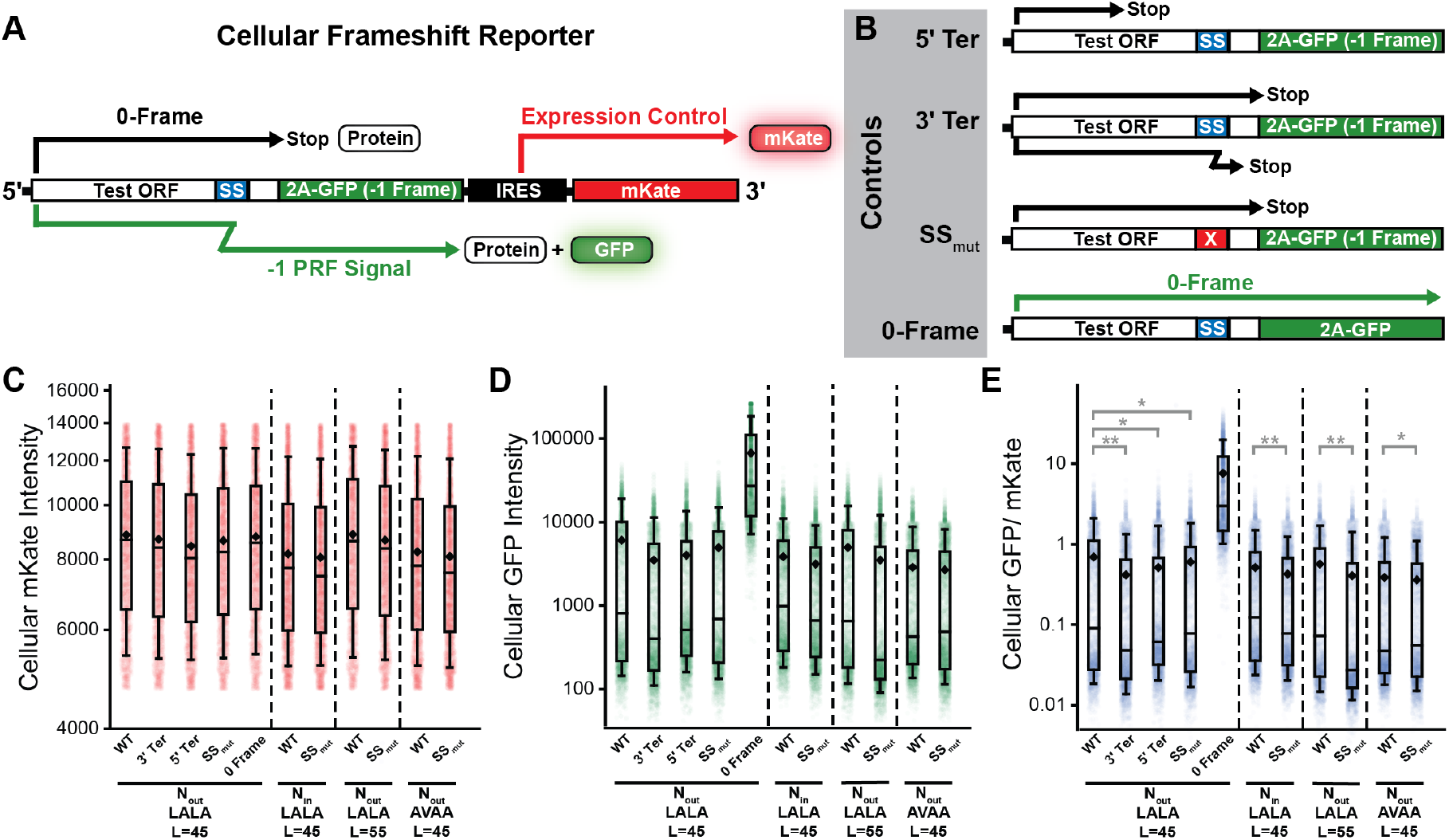
Cellular Measurements of LepB Programmed Ribosomal Frameshifting. A) A schematic outlines the general structure of the within the fluorogenic cellular -1PRF reporters. B) A series of schematics outline the nature of the control constructs utilized to validate the interpretation of the fluorescence signals generated by the test constructs. The 5’ Ter control should ablate the GFP signal by introducing a 0-frame stop codon upstream of the slip-site. The 3’ Ter control should ablate the GFP signal by introducing a stop codon in the -1 frame downstream of the slip-site. The SS_mut_ control introduces mutations in the slip-site that should reduce -1PRF and ablate the GFP signal. The 0-Frame control contains a single base deletion downstream of the slip-site that places the 2A-GFP in the 0-frame, which provides an estimate for the maximal GFP intensity. C) A box and whisker plot depicts the distribution of single-cell mKate intensity values among HEK293T cells transiently expressing various LepB sensor constructs from a representative biological replicate. D) A box and whisker plot depicts the distribution of single-cell GFP intensity values among HEK293T cells transiently expressing various LepB sensor constructs from a representative biological replicate. E) A box and whisker plot depicts the distribution of single-cell GFP: mKate ratios among HEK293T cells transiently expressing various LepB sensor constructs from a representative biological replicate. The mid-lines within boxes represent the median intensity values while the upper and lower edges of the boxes reflect the 75^th^ and 25^th^ percentile values, respectively. The upper and lower whiskers reflect the values of the 90^th^ and 10^th^ percentile values, respectively. Instances in which cells expressing negative controls exhibit statistically significant decreases in GFP: mKate values relative to the test construct in all three biological replicates according to a two-tailed Mann-Whitney U-test (*p* < 0.05) are indicated with a double asterix. Instances in which cells expressing negative controls exhibit statistically significant decreases in GFP: mKate values for two of three biological replicates are indicated with a single asterix.

### Bioinformatic Search for Ribosomal Frameshift Sites in Membrane Proteins

Previous efforts to search prokaryotic and eukaryotic transcriptomes for ribosomal frameshift sites have focused exclusively on structural features within mRNA.^32^ We therefore leveraged our biochemical findings to carry out a bioinformatic search for natural TMD-slip motifs within the human transcriptome. Briefly, we analyzed the ENSEMBL database using TOPCONS2,^33^ which identified 70,323 regions encoding TMDs within 24,678 total membrane proteins transcripts including 9,284 natively-spliced (canonical) transcripts and 15,394 alternatively-spliced isoforms. Based on Turner nearest neighbor parameters,^34^ we then identified a comprehensive set of 465 slippery heptamers (out of 16,384 possible heptamers total) for which the free energy difference in the codon-anticodon base pairing energetics in the 0- and −1 reading frames is negligible (see *Supplemental Theory*, Doc. S2). For each slippery heptamer within the transcriptome, we then identified the TMD that is closest to the ideal spacing (i.e. 45 codons). In cases where multiple slippery sequences were assigned to the same TMD, we selected only the heptamer that lies closest to the ideal spacing in order to avoid double-counting (see *Methods*).

The top candidate motif within these transcripts vary considerably with respect to distance. Overall, we identify 762 canonical transcripts that contain at least one TMD-slip motif with ideal spacing between the TMD and slippery sequence (L= 45, Doc. S3). We carried out two statistical analyses that each suggest the positions of slippery heptamers relative to TMDs appears is non-random (see *Supplemental Theory*). For instance, the observed distribution of distances between slippery heptamers and TMDs is significantly different from the corresponding distance distributions across an ensemble of reference transcriptomes with randomized TMD positions (16,384 iterations, Mann-Whitney U-test, *p* = 1.2 × 10^−10^). Similarly, a comparison of the observed distance distribution for slippery heptamers to those observed for 16,384 distinct randomized sets of 465 heptamers also suggests the observed TMD-slip motifs are unlikely to occur by random chance (Mann-Whitney U-test, *p* = 4.9 × 10^−48^). Notably, there are fewer natural TMD-Slip motifs that have an ideal spacing (L = 45 codons, 864 motifs) than are observed, on average, across the reference transcriptomes in which we score distances among randomized sets of heptamers (1,782 motifs, *χ*^2^ odds ratio = 0.60, *p* = 2.0 × 10^−48^, Fig. 4A), which suggests the ideal motifs are generally subject to negative selection.

**Figure 4.**
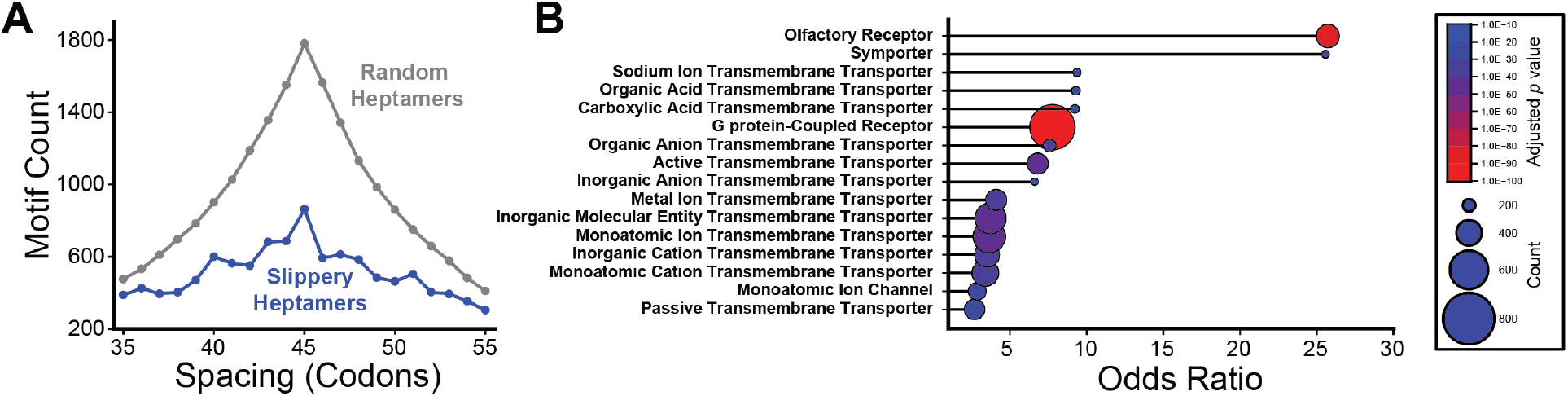
Bioinformatic Search for TMD-Slip Motifs within the Human Transcriptome. A) A histogram depicts the relative abundances of TMD-slip motifs in canonical transcripts for which the spacing between the heptamer and the region encoding the TMD range from 35-55 codons. The distribution for motifs containing the 465 slippery heptamers (blue) are plotted against the average number of motifs for 16,384 randomized sets of heptamers (gray). The peak at 45 codons occurs in both distributions because heptameric components of each motif are assigned to TMD regions that are positioned closest to the ideal spacing (L = 45 codons). B) A lollipop plot depicts the odds ratios associated with statistically enriched (>1.4-fold) gene ontology molecular function terms (depth 3-8) that are associated with at least 100 canonical transcripts bearing at least one TMD-slip motif (L = 35-55). The size of each point scales with the number of transcripts containing TMD-slip motifs that are associated with each term and are colored by the *p*-value associated with the statistical enrichment.

Based on our biochemical and cellular results (Figs. 1-3), there remains some uncertainty regarding how the activity of a motif may vary as a function of the distance between the slippery sequence and TMD. Given this caveat, we expanded the scope of our bioinformatic search to include all possible TMD-slip motifs with distances ranging from 35-55 codons. Using these criteria, we identified 25,097 potential motifs with distances ranging from L = 35 to L = 55 within the human transcriptome, 10,726 of which fall within natively-spliced transcripts. A gene ontology analysis of transcripts containing these TMD-Slip motifs suggest the translation of several protein families ranging from G protein-coupled receptors (GPCRs) to various classes of transporters and ion channels may be regulated by these motifs (Fig. 4B, Doc. S4). Notably, this list of candidates includes 2,395 potential TMD-slip motifs within canonical transcripts encoding 956 unique disease-linked integral membrane proteins. Together, these results identify a variety of new genes that are potentially translationally regulated by ribosomal frameshifting.

### Proteomic Validation of TMD-Slip Activity

To validate the activity of these motifs, we adapted a recent chemoproteomic analysis^35^ to search for predicted -1PRF products within a publicly-available deep proteomic dataset containing peptides derived from six distinct proteolytic digests of proteomic extracts from six cell lines that were analyzed with three ion fragmentation methods.^22^ We first reanalyzed the raw mass spectrometry data using MSFragger and removed any spectra that correspond to 0-fame peptides. We then carried out a second-pass analysis to identify frameshifts by querying the remaining spectra against a custom in-house database of predicted frameshifted peptides derived from *in silico* digests of the potential frameshift products. The results of this two-pass search identified 33 unique peptides corresponding to predicted frameshift products (1% FDR, Doc. S5). The identified peptides include 14 “transitional” peptides that consist of a string of 0-frame amino acids upstream of the slippery sequence followed by one amino acid encoded in an alternative reading frame. We also found six “premature termination” peptides that are predicted to arise from a direct ribosomal transition from the 0-frame to a −1 stop codon. Finally, we identified 13 peptides that are fully encoded in an alternative reading frame downstream of a TMD-slip motif. Notably, the alternative amino C-terminal amino acids of the transitional peptides suggest TMD-slip motifs may promote diverse frameshift transitions including canonical (two t-RNA), hungry (one-tRNA) −1 frameshifting, and even hungry −2 frameshifting (Doc. S5, see *Discussion*). The unexpected identification of hungry −2 frameshift products is also supported by the identification of several other fully-frameshifted peptides that are fully encoded in the +1 reading frame (Doc. S5). Together, these identifications provide clear experimental confirmation of the activity of these motifs in the context of native transcripts expressed in live cells.

Overall, 16 of the identified peptides are derived from the translation of integral membrane proteins, four of which correspond to peptides that are fully-encoded within the −1 frame of the lysophosphatidylcholine acyltransferase 1 transcript (LPCAT1, Fig. 5 A-B). Interestingly, the sequences of these frameshifted peptides are nested within an uninterrupted string of 121 codons within −1 frame downstream of its TMD-slip motif. Interestingly, AlphaFold predictions suggest this ribosomal frameshift should completely remodel the structure of its lumenal subdomain (Fig. 5A). In addition to integral membrane proteins, we identify 13 peptides derived from transcripts encoding secreted proteins that contain an N-terminal signal peptide, including HSP90B1 and four other ER chaperones (Doc. S5). Unlike the LPCAT1 motif, frameshifting in HSP90B1 results in prompt termination following the decoding of only eight codons in the −1 reading frame (Fig. 5C). Thus, while the recoding site in LPCAT1 potentially results in the translation of a functionally distinct protein, the motif in HSP90B1 and most others should simply prevent the translation of their functional domains. Finally, we identify three premature termination peptides that are encoded in alternatively spliced transcripts that normally encode proteins with N-terminal mitochondrial targeting sequences (Doc. S5). Overall, the distinct properties of these transcripts and motifs highlight potential differences in their regulatory context (see *Discussion*).

**Figure 5.**
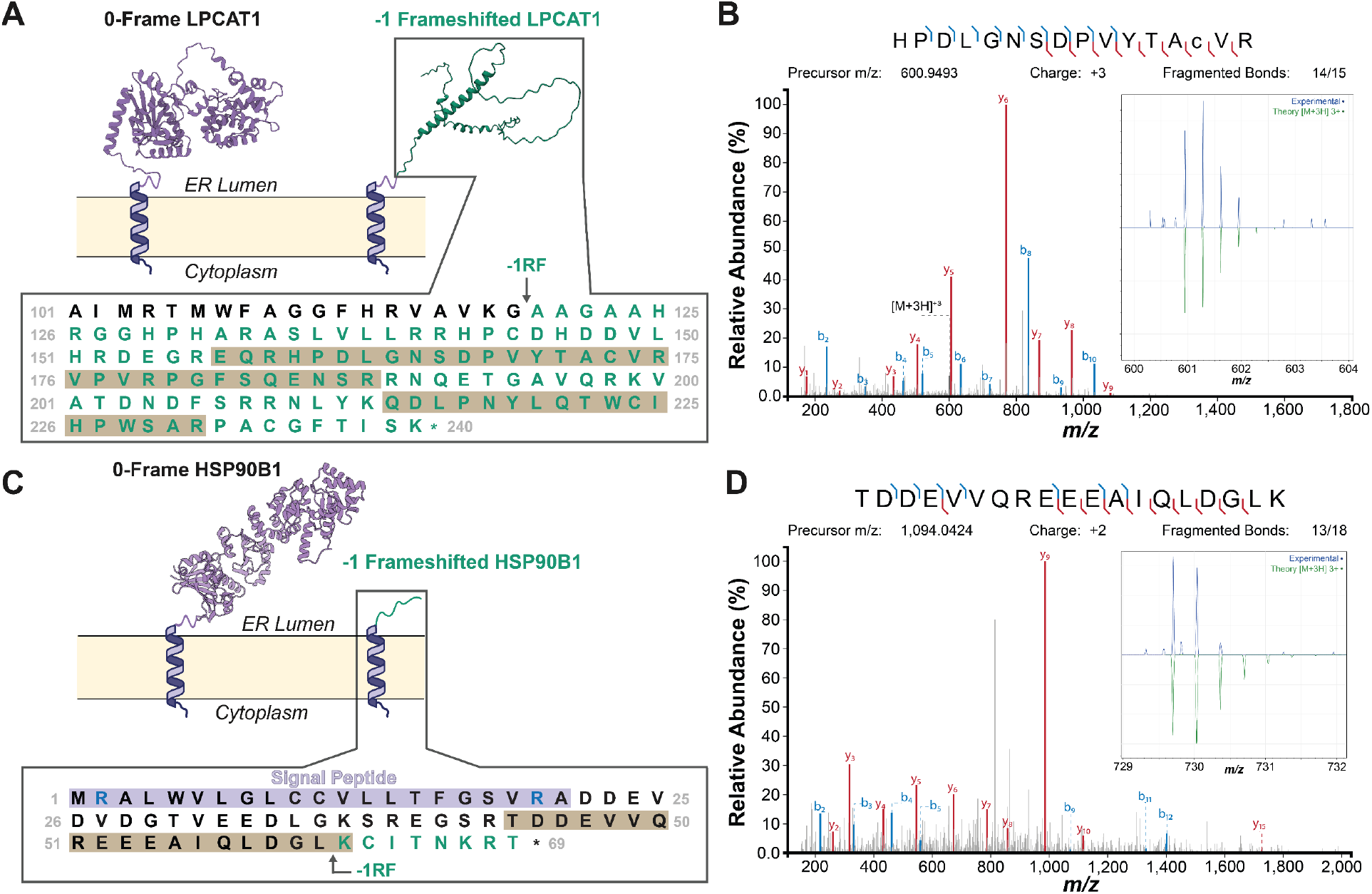
Proteomic Identification of Frameshifted Peptides. Two representative proteomic identifications of ribosomal frameshift products are shown. A) A cartoon depicts the structural effects of a ribosomal frameshift caused by a TMD-slip motif within the LPCAT1 transcript. AlphaFold3 models of its canonical (purple) and frameshifted (green) soluble domains are shown, the latter of which is predicted to contain 121 amino acids encoded in the −1 frame. The sequence of the transition from 0-frame amino acids (black) to frameshifted amino acids (green) is shown in relation to the positions of the identified peptides (gray). B) The HCD product ion spectrum for one of the frameshifted LPCAT1peptides is shown. The spectrum of the precursor along with the corresponding theoretical spectrum are shown in the inset for reference. C) A cartoon depicts the structural effects of a ribosomal frameshift caused by a TMD-slip motif within the HSP90B1 transcript. A crystallographic model of the luminal domain encoded in the 0-frame is shown (PDB 2O1U, purple). The frameshift product is predicted to generate only eight amino acids encoded in the −1 frame (green). The sequence of the transition from 0-frame amino acids (black) to frameshifted amino acids (green) is shown in relation to the position of the identified peptide (gray). D) The HCD product ion spectrum of the transitional frameshift peptide for HSP90B1 is shown. The spectrum of the precursor along with the corresponding theoretical spectrum are shown in the inset for reference.

### Impact of Splicing on TMD-Slip Activity

Notably, seven of the identified frameshift peptides can only be generated by transcripts that have undergone alternative splicing (Doc. S5). This class of peptides are representative of a wider set of 2,752 putative TMD-slip motifs (L = 35-55) that are specifically created by alternative splicing (Doc. S3). In these cases, differences in the relative positions of exons alter the coding sequence such that a TMD is more likely to occupy the translocon while the ribosome is decoding a slippery heptamer. To explore how splicing may alter TMD-slip activity, we characterized a splicing-dependent motif we identified within transcripts encoding the tetrameric voltage-gated potassium channel KCNQ1 (Fig. 6A). Our informatic search flagged an X_1_ XXY_4_ YYZ_7_ motif (A_1260_ AAA_1263_ AAG_1266_) nested within an adenosine-rich region of several KCNQ1 transcripts. The closest TMD to this heptamer in the canonical transcript (ENST00000155840.12) is too distant for translocon-mediated coupling between the nascent chain and peptidyl transferase center (L= 67 codons, Fig. 6B), though RNA secondary structures could potentially prolong its decoding (Fig. 6C). Alternatively, we identify a putative splice variant in which this segment instead falls only 39 codons downstream of a chimeric, non-native TMD (ENST00000646564.2, Fig. 6B). Interestingly, cells expressing a -1PRF reporter containing the initial 1365 bp of the natively-spliced KCNQ1 transcript generate GFP: mKate ratios that are higher than all three negative controls (see Figs. 3 A-B, Fig. 6 D-F, and Doc. S1). Our measurements suggest -1PRF is 1.8 ± 0.1% efficient at this site despite non-ideal spacing between the TMD and the slippery sequence, which potentially suggest the RNA structure alone may promote frameshifting. Indeed, this recoding event is insensitive to disease mutants that induce KCNQ1 misfolding (Table S2). However, introducing these splicing variations into the KCNQ1 reporter increases cellular GFP: mKate ratios 8-fold (-1PRF efficiency = 14.5 ± 0.7%, Fig. 6F). These results demonstrate how the splicing-mediated reorganization of the nascent chain can dramatically enhance ribosomal frameshifting.

**Figure 6.**
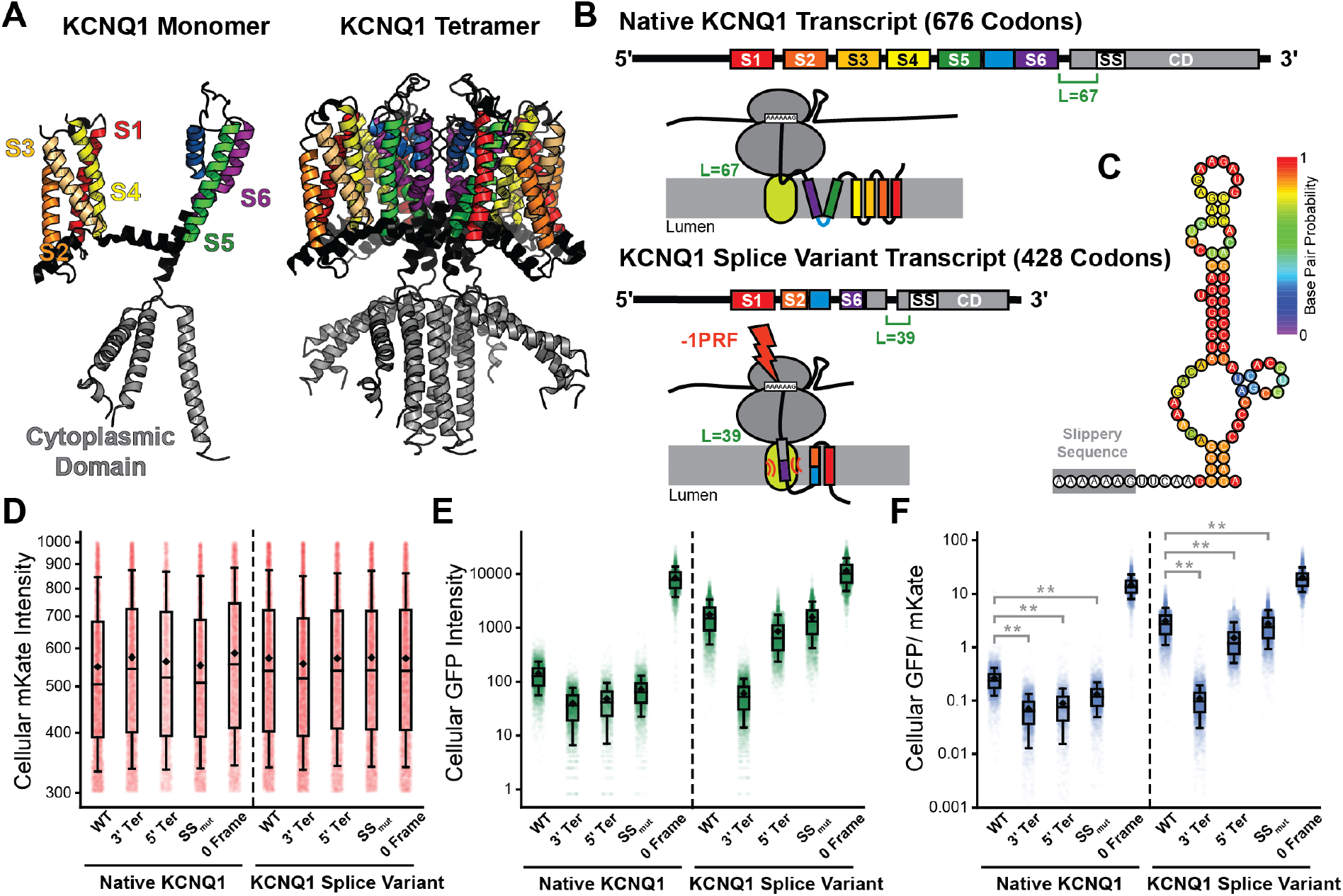
Programmed Ribosomal Frameshifting in KCNQ1. A) A structural model (PDB 6V00) of the native KCNQ1 monomer (left) and tetramer (right) are shown. The TMDs (S1-S6) and cytoplasmic domain are labelled for reference. B) Cartoons depict the relative positions the TMDs in the transcripts encoding the native and alternatively spliced KCNQ1 as well as the relative orientations of these domains with respect to the ribosome and translocon at the point of frameshifting. C) A cartoon depicts the predicted RNA secondary structure directly adjacent to the KCNQ1 slip-site. Bases are colored according to their base pairing probabilities as predicted by the ViennaRNA web server. D) A box and whisker plot depicts the distribution of single-cell mKate intensity values among HEK293T cells transiently expressing -1PRF sensors for WT and alternatively-spliced KCNQ1 from a representative biological replicate. E) A box and whisker plot depicts the distribution of single-cell GFP intensity values among HEK293T cells transiently expressing -1PRF sensors for WT and alternatively spliced KCNQ1 from a representative biological replicate. F) A box and whisker plot depicts the distribution of single-cell GFP: mKate ratios among HEK293T cells transiently expressing -1PRF sensors for WT and alternatively spliced KCNQ1 from a representative biological replicate. The mid-lines within boxes represent the median intensity values while the upper and lower edges of the boxes reflect the 75^th^ and 25^th^ percentile values, respectively. The upper and lower whiskers reflect the values of the 90^th^ and 10^th^ percentile values, respectively. Mann-Whitney U-tests suggest the cellular GFP: mKate intensity ratios of cells expressing the test construct are significantly higher than those of the corresponding negative controls (** *p* < 0.001).

Overall, transcript isoforms bearing these “splicing-dependent” motifs can be found in isoforms of transcripts that are derived from 1,439 different genes, which corresponds to ~25% of the TMD-slip containing genes overall. Interestingly The gene ontology terms associated with genes containing these splicing-dependent TMD-slip motifs are again enriched with certain molecular functions associated with transporters and ion channels (Fig. 7A). While several of these same terms are associated canonical transcripts containing TMD-slip motifs (Figs. 4B and 7A), these splicing-dependent motifs also exhibit distinct statistical enrichments for other terms such as “ATP Binding” (Doc. S4, Fig. 6B). Moreover, unlike those within canonical transcripts, splicing-dependent TMD-Slip motifs *do not* appear to be common among GPCR transcripts (Doc. S4). These findings suggest the splicing-dependent TMD-slip motifs may serve distinct biological purposes relative to those within canonical transcripts. Nevertheless, we note that these frameshift events effectively promote premature termination, which is likely to decrease the processivity of translation in the context of these improperly spliced transcripts.

**Figure 7.**
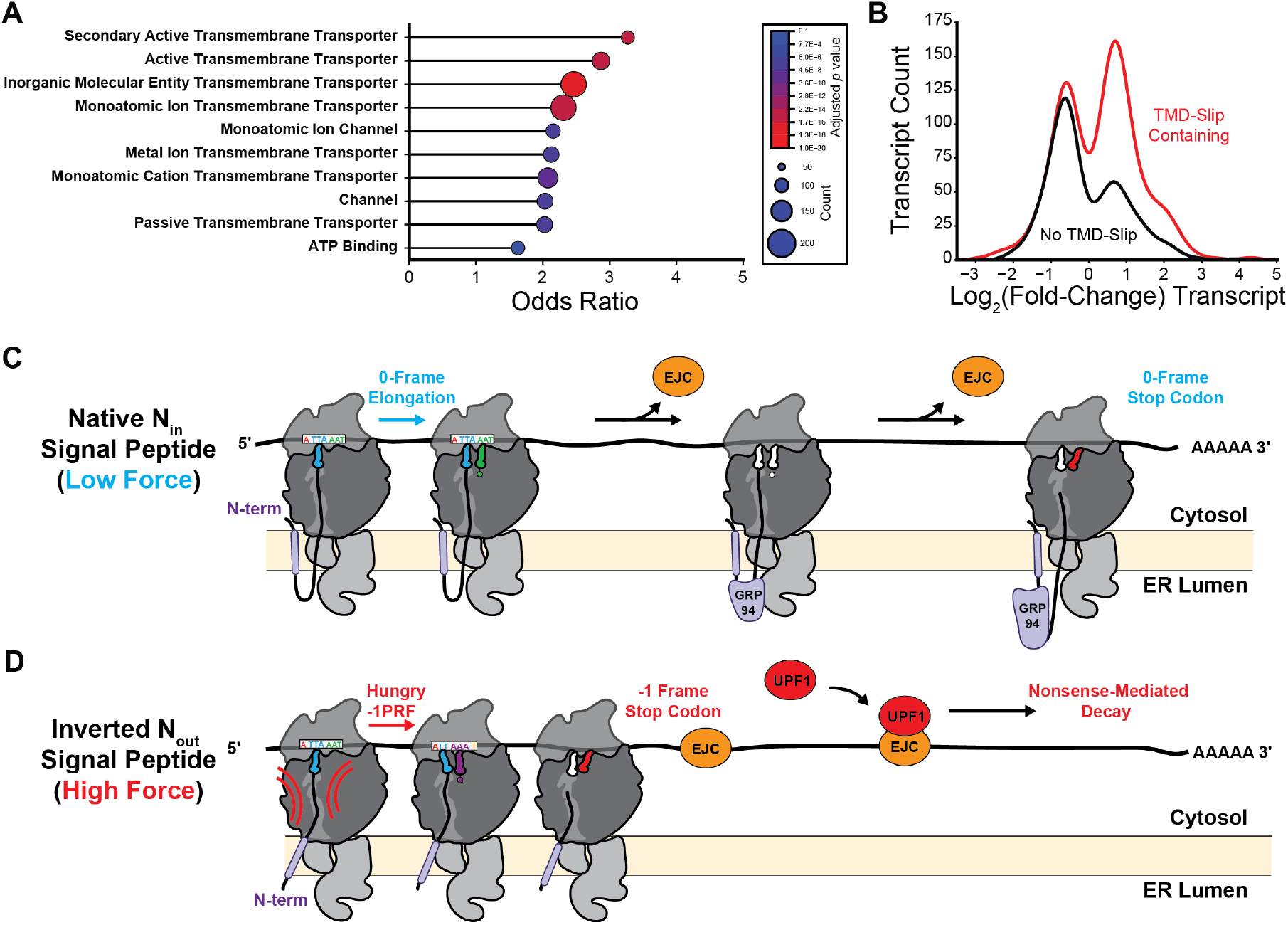
TMD-Slip Motifs in Alternative Splicing and RNA-Decay. A) A lollipop plot depicts the odds ratios associated with statistically enriched (>1.4-fold) gene ontology molecular function terms (depth 3-8) that are associated with at least 100 genes for which alternatively spliced isoform transcripts contain motifs that are not present in the corresponding canonical transcript (i.e. canonical L < 35 or L > 55). The size of each point scales with the number of genes containing splicing-dependent TMD-slip motifs that are associated with each term and are colored by the *p*-value associated with the statistical enrichment. B) A histogram compares log_2_(fold-change) abundance upon UPF1 degradation for membrane protein transcripts lacking (black, n=425) or containing (red, n=730) TMD-slip motifs. Log_2_(fold-change) values represent the average value for each transcript across all conditions with UPF1 depletion reported in reference 35. A kernel smoothing function was used to render each histogram. C) A cartoon depicts the progressive translation on the HSP90B1/ GRP94 transcript. When the signal peptide adopts its native N_in_ orientation, elongation proceeds in the 0-frame and ribosomes displace exon-joining complexes (EJC, orange) before they can recruit UPF1 (red). D) A cartoon depicts a putative scenario in which a topological defect triggers -1PRF. Topological inversion of the signal peptide increases tension on the nascent chain, which triggers -1PRF and premature termination. Stalled ribosomes then fail the clear EJCs from the transcript, which then recruit UPF1 to trigger nonsense-mediated decay of the transcript.

### Connections to Nonsense-Mediated mRNA Decay

Frameshifting at TMD-slip motifs results in the extension of the nascent chain by only 24 amino acids encoded in the −1 frame, on average, prior to termination (Doc. S3). This raises doubt as to whether most of these frameshifted proteins could potentially encode novel functions. Nevertheless, the premature termination caused by frameshifting events can destabilize their transcripts by triggering nonsense mediated decay (NMD); an ancient regulatory pathway that degrades transcripts bearing premature termination codons.^36^ To determine whether TMD-slip motifs predispose transcripts to NMD, we leveraged a recent characterization of NMD substrates^37^ to assess whether the NMD regulator UPF1 generally modulates the levels of the transcripts identified by our informatic search. Overall, TMD-slip motifs are present in 72% of membrane protein transcripts that exhibit increased abundance upon UPF1 degradation in HEK293 and/ or HCT116 cells (Doc. S3).^37^ Indeed, TMD-slip containing transcripts are statistically enriched among upregulated transcripts (Fisher’s Exact Test Odds Ratio = 1.6, *p* = 1.6 ×10^−6^) and depleted among downregulated transcripts (Fisher’s Exact Test Odds Ratio = 0.6, *p* = 1.9 ×10^−6^). The abundance of the 730 observed membrane protein transcripts bearing at least one TMD-slip motif increase by an average of 20% in response to UPF1 degradation (Mann Whitney U-test *p* = 1.2 × 10^−9^, Fig. 7B), which implies that many transcripts containing TMD-slip motifs are NMD substrates. Together, these findings suggest many TMD-slip motifs serve to destabilize their transcript, potentially in response to the misassembly of the nascent polypeptide.^9^

## Discussion

Most known -1PRF motifs are comprised of an RNA structure (or an ensemble of RNA structures)^38^ that prolongs the decoding of an adjacent slippery heptanucleotide sequence.^1^ Together, these transcript features increase the propensity of the ribosome to transition into the −1 reading frame and produce a frameshifted polypeptide. While these features play a central role in -1PRF activity, the net recoding efficiency can be tuned by a variety of additional effectors.^4,10^ Recent findings have revealed that cotranslational protein folding transitions that coincide with -1PRF can have a pronounced effect on recoding efficiency.^19–21^ Our investigations of the SINV structural polyprotein suggested this coupling is mediated by mechanical pulling forces on the nascent chain,^19,20^ which are generated at various points in cotranslational protein folding and translocation.^24,25,39,40^ Together, these considerations suggest fluctuations in mechanical tension on the nascent polypeptide may dynamically modulate translational recoding throughout elongation. However, both current examples of direct coupling between the nascent chain and -1PRF also involve viral RNA features that impact the net -1PRF efficiency.^4,19–21^ It is therefore difficult to discern from these examples whether the nascent chain impacts translational recoding more generally.

In this work, we leverage recent observations from viral systems to design and characterize novel -1PRF sites within a transcript encoding a chimeric *E. coli* LepB protein.^23,41,42^ Surprisingly, we show that the translocation of a nascent TMD is sufficient to trigger efficient -1PRF so long as it coincides with the decoding of a slippery heptamer. Notably, LepB -1PRF appears to be significantly more efficient *in vitro* than in HEK293T cells (Table S1), which likely arises from non-native translation kinetics *in vitro*. Given the apparent lack of an ideal LepB stem-loop or pseudoknot to pause the ribosome, differences in tRNA concentrations could potentially represent a key kinetic determinant of the ribosomal dwell time. Delayed delivery of tRNAs to the A-site in the context of a lysate could potentially result in “hungry” (one-tRNA) frameshifting,^17,18^ which we indeed observe at in our proteomic identifications of various human membrane proteins (Doc. S5). Notably, the LepB frameshift products also appear to vary in their glycosylation state, which may suggest -1PRF does not require proper ribosomal targeting to a translocon (see Figs. 1C & 2B-C). However, we cannot rule out that these frameshift products were instead generated via translocon complexes with variable OST occupancy.^28,29,43^ Considering translocons are dynamically remodeled during membrane protein biogenesis,^27–30^ it is also possible that the recruitment of distinct translocons may modulate the activity of the LepB and/ or other TMD-slip motifs.

Our results provide firm evidence that PRF occurs during the translation of human proteins (Figs. 4-5, Doc. S5). We identify 303 TMD-slip motifs that fall upstream of at least 100 amino acids encoded in the −1 reading frame including the LPCAT1 (Doc. S3, Fig. 5 A-B). In these rare cases, translational recoding could potentially produce functionally distinct proteins. Nevertheless, most TMD-slip motifs are instead followed a host of out-of-frame stop codons that preclude run-on translation in alternative reading frames.^44,45^ These motifs should trigger premature termination, which is typically cast as a costly translational error.^46^ However, there are many circumstances in which termination may improve fitness such as at the onset of cellular starvation^7^ or proteostatic collapse.^9^ In this work, we identify several additional roles for eukaryotic -1PRF. For instance, we identify 1,439 genes that bearing motifs that are selectively generated in response to alternative splicing (Doc. S3). In the case of KCNQ1, we show that a loss of exons creates an untimely coupling between the translocation of a nascent TMD and the decoding of a slippery sequence (Fig. 6B). We also validate frameshifts downstream of various ER QC proteins, such as HSP90B1, that rely on signal peptides for secretion in the ER lumen (Fig. 7 C-D). Proper secretion requires signal peptides to adopt an N_in_ topology, which is typically specified by the positioning of basic residues within the inner membrane leaflet (the “positive-inside” rule).^47^ Strangely, the HSP90B1 signal is flanked by arginine residues at both its N- and C-termini (Fig. 5C); a mark of “frustration”^48^ that could promote topological heterogeneity. Notably, the native N_in_ topology should generate less tension on the ribosome relative to the inverted, non-productive topology.^24,25^ We suspect this differential force profile may enhance frameshifting in response to translocation errors (Fig. 7 C-D). In either case, the resulting premature termination is likely to trigger nonsense-mediated transcript decay.^36^ This interpretation is corroborated by the negative regulation of TMD-slip transcripts by the NMD regulator UPF1 (Fig. 6A, Doc. S3). Taken together, our findings suggest a mechanism by which aberrant splicing, improper targeting, and/ or the repeated misassembly of nascent polypeptides^9^ may ultimately prompt the destruction of the template and the disassembly of the polysome. This mechanism could be of crucial importance in the regulation of proteostasis for large proteins that rely on numerous chaperones for proper assembly such as CFTR,^9,49^ as the absence of a single assembly factor within the polysome could produce many defective channels. TMD-slip motifs are present in 956 transcripts encoding unique disease-linked integral membrane proteins, suggesting this translational regulation may be involved in a variety of proteostasis diseases.

Strikingly, our search identified TMD-Slip motifs in 83% of all GPCR-encoding transcripts and 72% of all ion channel transcripts. Nevertheless, we suspect certain aspects of our findings are also likely generalizable to water-soluble proteins as well. In the context of our artificial TMD-slip motifs, we primarily observe activity when the region encoding the TMD resides ~45 codon upstream of the X_1_ XXY_4_ YYZ_7_ motif (Fig. 1 E-G). This corresponds to the specific distance at which translocation forces should spike while the PTC decodes the slippery heptamer.^24,25^ Coupling between assembly and frameshifting may therefore be more generally mediated by properly-timed mechanical tensions within the nascent chain.^19,20^ Though it remains unclear how this tension modulates frameshifting, conformational transitions in the nascent chain could potentially stall translation by decreasing the rate of the peptidyl transfer reaction^50^ and/ or by promoting allosteric transitions that occlude the peptidyl transferase center.^51^ Tension on the nascent chain could therefore potentially dampen elongation kinetics in a manner that mimics the effects of RNA structures typically associated with -1PRF. We note that our mass spectrometry identifications include several −2 frameshift transitions (Doc. S5), which suggests the nascent chain may alter the kinetic barriers that maintain the reading frame.^14^ These mechanistic interpretations remain speculative and additional investigations are needed to gain insights into the coupling between the nascent chain and PTC. But regardless of the mechanism, it is important to note that a variety of molecular transitions including chaperone interactions,^52^ ligand binding,^52,53^ and folding within/ near the exit tunnel^39,40,53,54^ are all capable of altering pulling forces on the nascent chain. Thus, variations in translational recoding and any resulting changes in the processivity of translation could potentially be coupled to a variety of cotranslational processes. Additional investigations are needed to ascertain the full scope of cotranslational recoding events in eukaryotes and how these are involved in the regulation of various biochemical processes.

## Materials and Methods

### Plasmid Design and Production

Biochemical -1PRF reporter constructs were made in the context of a previously described chimeric LepB topology reporter gene developed by the von Heijne group.^23,42^ Briefly, slippery sequences (U_1_ UUU_4_ UUU_7_) were introduced at various positions (35, 45, or 55 codons) downstream of the guested “H-segment” TMD (ALAALALAALAALALAALA or AAAAVAAAAAAAAAVAAAA). To simplify the banding pattern of the LepB protein products, site directed mutagenesis was used to remove a consensus glycosylation site that is conditionally modified in response to membrane integration of the H-segment. To improve the detection of the resulting translation products by ^35^S-labelled methionine, conservative substitutions were made to introduce seven additional methionine residues into the LepB protein. Finally, to improve the resolvability of the full-length and -1PRF products, we extended the C-terminal tail of the LepB protein in order to selectively increase the size of the full-length, non-frameshifted LepB protein. To invert the topology of the guested TMD, we used the NEBuilder kit (New England Biolabs, Ipswitch, MA) to introduce a gene block (Integrated DNA Technologies, Coralville, IA) encoding an additional TMD between the second LepB TMD and the H-segment. A comprehensive, annotated list of *in vitro* -1PRF reporter sequences can be found in Doc. S1.

The cellular PRF reporters described herein represent a modified version of a previously described bicistronic system in which mKate is produced as a result of -1PRF and GFP is constitutively translated from a downstream IRES.^19,20^ To increase the sensitivity of this reporter, we used the NEBuilder kit to switch the order of the fluorescent proteins so that the higher intensity GFP protein is generated by -1PRF and an IRES-mKate cassette serves as an expression control. We also inserted a self-cleaving P2A linker upstream of the -1GFP cassette to ensure the frameshift reporter protein (GFP) is released from the upstream portions of the nascent chain. NEBuilder was then used to introduce segments of transcripts containing putative -1PRF sites upstream of the - 1GFP cassette. Inserted segments contain regions from the genes of interest beginning at the initial methionine codon and ending 75 bases downstream from the putative slip-site. Site directed mutagenesis was used to introduce point mutations and control modifications that require single mutations (ie 5’ Ter, 3’ Ter, and 0-frame controls), while controls that require extensive mutagenesis (ie SS_mut_) were produced by GenScript (Piscataway, NJ). Sequence-verified plasmids, which were all made in a pCDNA5 backbone featuring a CMV promoter, were prepped using a ZymoPURE Express Plasmid Midiprep Kit (Zymo Research Corporation, Irvine, CA) prior to transfection. A comprehensive, annotated list of cellular -1PRF reporter sequences can be found in Doc. S1.

### In vitro Translation

Biochemical LepB -1PRF reporters were translated *in vitro* as previously described.^55^ Briefly, PCR amplification was first used to generate linear templates for each LepB reporter cDNA from circular plasmid DNA. Linear templates were then used to synthesize mRNA for each reporter using the RiboMAX *in vitro* transcription kit (Promega Corporation, Madison, WI). Transcripts were then translated in the context of a rabbit reticulocyte lysate (Promega Corporation, Madison, WI) supplemented with canine pancreas rough microsomes (tRNA Probes, College Station, TX) and EasyTag L-^35^S labeled Methionine (Perkin Elmer Inc., Downers Grove, IL). Translation products were then separated on an 12% SDS PAGE gel, which was then dried and exposed to a phosphorimaging screen overnight. The screen was then imaged using a Typhoon Imager (GE Health, Chicago, IL). LepB protein bands were integrated using ImageJ software.

### Frameshift Reporter Measurements in HEK293T Cells

Fluorogenic frameshift reporters were characterized in the context of HEK293T cells (American Type Culture Collection, Manassas, VA), which were grown in a humidified incubator at 37°C with 5% CO_2_ in a complete Dulbecco’s modified eagle medium (DMEM, Gibco, Grand Island, NY) containing 10% (v/v) fetal bovine serum (FBS, Corning Life Sciences, Corning, NY) as well as penicillin (100 U/ml)/ streptomycin (100 μg/ml) (Gibco, Grand Island, NY). To express -1PRF reporters, these cells were transfected in Optimem media (Gibco, Grand Island, NY) using Lipofectamine 3000 in accordance with the manufacturer’s recommendations (Invitrogen, Carlsbad, CA). The transfection media was replaced with complete DMEM media 24 hours after transfection then harvested 48 hours after transfection using 0.25% trypsin-EDTA (Gibco, Grand Island, NY). Cells were then washed twice and resuspended in phosphate buffered saline containing 2% FBS prior to the removal of cellular aggregates using a 20 µm cell strainer. Single cell fluorescence profiles were then analyzed using either an LSRII or LSRFortessa flow cytometer (BD biosciences, Franklin Lakes, NJ). The fluorescence profiles of positively-transfected cells were then selectively analyzed based on their characteristic mKate fluorescence intensities using FlowJo software (Treestar, Ashland, OR). To ensure GFP frameshift reporter intensities were compared at a consistent expression level, our analysis was restricted to cells with mKate intensities that fall within two standard deviations of the overall average (across samples) of the population median IRES-mKate fluorescence intensities from all of the samples in the analysis. To estimate -1PRF efficiencies, observed intensities were first corrected for background fluorescence by subtracting the mean GFP and mKate intensities of negatively-transfected cells within each sample from the corresponding mean intensities of the positively transfected cells within the defined range of expression. Efficiency values were then calculated by dividing the average mKate: GFP value of cells expressing the test construct by the corresponding ratio of the 0-frame control in which GFP was instead produced from the 0-reading frame. Statistical differences in the distribution of cellular GFP: mKate values were assessed using a two-tailed Mann-Whitney U-test, which we calculated using Origin Pro software (Origin Lab, North Hampton, MA).

### Reverse Transcriptase PCR

To ensure cellular -1PRF reporter constructs did not undergo additional splicing modifications,^5^ we first used the Aurum total RNA mini kit (Biorad Laboratories, Hercules, CA) to extract the mRNA from HEK293T cells transiently expressing each -1PRF sensor. Cellular mRNA was then purified using the Monarch RNA cleanup kit (New England Biolabs, Ipswitch, MA) prior to reverse transcription of cellular mRNAs into cDNAs using the iScript Advanced cDNA Synthesis kit (Biorad Laboratories, Hercules, CA). -1PRF reporter cDNAs were then PCR amplified and analyzed by gel electrophoresis and Sanger sequencing. To ensure each module within each reporter transcript were present at comparable levels, the relative abundances of the test ORF (i.e. Lep or KCNQ1), the GFP -1PRF reporter cassette, and the IRES-mKate expression control were also verified via qPCR in accordance with recently published guidelines for bicistronic reporter controls (see Fig. S3).^56^ The sizes of each product were also confirmed by gel electrophoresis and next generation sequencing (see Fig. S3).

### Bioinformatic Search for TMD-slip motifs

The portions of nascent membrane proteins that undergo translocon-mediated membrane integration are often distinct from the segments that span the membrane in the context of the native structure.^57,58^ Thus, to identify the points in synthesis in which engagement of the nascent chain by the translocon is likely to generate a mechanical force, we developed a computational python module which first identifies regions that are likely to encode native structural TMDs then next uses a knowledge-based energy scale^42^ to identify the specific residues within this region that are most likely to undergo translocon-mediated membrane integration. Briefly, this script first employs TOPCONS2^33^ to scan through the 0-frame protein sequence and identify structural TMDs. The script then uses a python-based implementation of the ΔG predictor^42^ to calculate the membrane partitioning energetics for each possible 16-25 amino acid segment within a sequence window beginning 10 amino acids upstream and ending 10 amino acids downstream of the TMD region identified by TOPCONS2. The edge residues of the segments that are predicted to partition most readily from the translocon to the membrane (lowest ΔG) were used to calculate the spacings between the relevant sequence elements.

Though viral -1PRF motifs typically occur at X_1_ XXY_4_ YYZ_7_ sites,^1^ there are a variety of heptanucleotide sequences that are capable of enhancing ribosomal frameshifting by facilitating comparable codon-anticodon base pairing energetics within the 0- and −1 reading frames.^11,13^ Therefore, to broaden our search for potential TMD-slip motifs, we developed a second python module that scans through every subsequent heptanucleotide sequence of a transcript then uses a knowledge-based energy scale to approximate the free energy difference between the tRNA anticodon base pairing interactions within the 0-frame A- and P-site codons in relation to that of the corresponding −1 reading frame codons (Δ*G*_FS_). We then defined a range of values for slippery heptanucleotide sequences based on the range of scores associated with 24 known X_1_ XXY_4_ YYZ_7_ motifs (ΔG_FS_ < +2.9 kcal/ mol, Doc. S2). Overall, our scoring system identified 465 of 16,384 total heptanucleotide sequences that are likely to enhance the propensity of the ribosome to undergo -1PRF (see *Supplemental Theory*).

Finally, we combined these modules to identify TMD-slip motifs within a comprehensive set of validated and predicted human transcripts (Ensembl CDS FASTA, Release 113, 08/15/2024). For simplicity, we used Ensembl annotations to differentiate canonical and alternative isoform transcript sequences (Doc. S3). We first used the TMD module to identify all transcripts that are predicted to contain at least one TMD. Transcripts without any predicted TMDs were then discarded from the subsequent analysis. We then used our base pairing energetics module to score every heptanucleotide sequence within each of these membrane protein transcripts. The overall distribution of TMD-slip distances was then calculated by identifying slippery heptamers (ΔG_FS_ < +2.9 kcal/ mol) that lie closest to the optimal upstream position relative to the TMD (45 codons). Gene ontology analyses were carried on a subset of top hits that contained a putative TMDs that lie 35-55 codons downstream of a slippery heptamer (Doc. S4). Our final code can is freely available online (GitHub). For more detailed information, see *Supplemental Theory*.

### Proteomic Identification of Frameshift Products

Protein-level evidence for ribosomal frameshift products was sought in public mass spectrometry datasets. Raw mass spectrometry data was obtained from MassIVE (MSV000086944), a large dataset containing analyses of peptides derived from six distinct proteolytic digests of proteomic extracts from six cell lines, which were then pre-fractionated and analyzed using three ion fragmentation modes.^22^ Mass spectra were analyzed using MSFragger (version 23.1).^59^ Post-search rescoring was carried out using MSBooster and Percolator.^60^ IonQuant (version 1.11.11)^61^ was used for label-free quantification. Database searching was performed using a previously described^35^ two-pass strategy in which a database of zero-frame products was first searched and matched spectra were removed prior to a secondary search against a database of frameshifted products. Percolator weights determined in the first pass were reused in the second pass to ensure identified peptides were rescored in the same context. Detailed search results are available as a reanalysis on the PRIDE database (PXD072350). Identified peptides were classified as zero frame, transitional, frameshift, or a combination using an in-house query tool and *in silico* digest of predicted frameshift products, which is available on the Schlebach Lab GitHub.

A protein-BLAST search and alignment was performed against non-redundant human sequences for peptides that were exclusively classified as transitional or frameshift to manually validate the predicted classifications.

## Supporting information

Doc S1 Reporter Sequences

Doc S2 Heptamer Scoring

Doc S3 TMD Slip Motifs

Doc S4 GO Enrichments

Doc S5 MS Table

## Data and Code Availability

Code for the bioinformatic identification, statistical analysis, gene ontology analysis, and proteomic analysis of TMD-slip motifs and can be found on the Schlebach Lab GitHub page (https://github.com/schlebachlab/PRF-Search). Original experimental data described herein have been freely shared in a Mendeley Data directory (https://data.mendeley.com/datasets/5rs7tfjfym/2). The mass spectrometry proteomics data have been deposited to the ProteomeXchange Consortium via the PRIDE partner repository^62^ with the dataset identifier PXD072350 and 10.6019/PXD072350. The sequences of all genetic constructs described herein can be found in Doc. S1. All materials described herein will be made freely available by J.P.S. upon request.

## Acknowledgements

We thank Bil Clemmons, Thomas Miller III, Huanghao Mai, Jonathan Dinman, David Giedroc, Marina Rodnina, Suchetana Mukhopadhyay, and Josh Arribere for their support and/ or for their helpful input. We thank Christiane Hassel and Jill Hutchcroft for technical input and assistance. We also acknowledge the support of the Indiana University Flow Cytometry Core Facility, the Purdue University Flow Cytometry and Cell Separation Facility, and the Indiana University Center for Genomics and Bioinformatics. This research was supported in part by grants from the National Institute of General Medical Sciences to J. P. S. (R01GM138845 and R35GM152086).

