## Supplemental Materials for "Feedback from the Nascent Chain Triggers Ribosomal Frameshifting and Transcript Decay"

†Contributed Equally
*Corresponding Authors: jschleba (at) purdue.edu, cpkuntz (at) purdue.edu, bsdrown (at) purdue.edu

**Contents**

-Figure S1
-Figure S2
-Figure S3
-Table S1
-Table S2
-Supplemental Theory

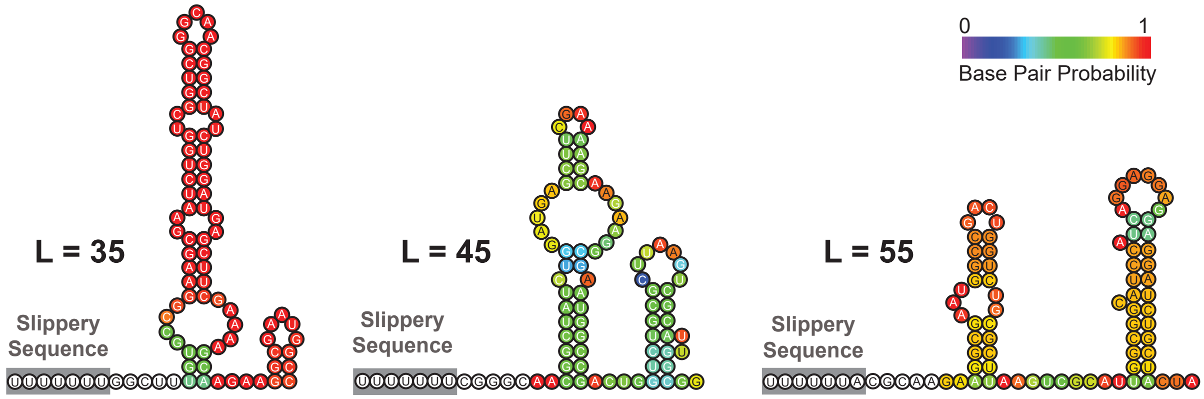


**Figure S1. Secondary Stucture Predictions for LepB Frameshift Reporter Constructs.** The cartoons above depict the basepairing interactions within the 75 basepair region beginning 5 bases downstream of the slippery sequences that were incorporated into various positions of the leader peptide frameshift reporter transcript, as is predicted by ViennaRNA web server. Bases are colored according to their predicted base pairing probabilies as is indicated by the color bar. The position of the engineered poly-uridine slippery sequence relative to the guest TM domain is indicated for each construct. We excluded the slippery sequences and the five adjacent 3' bases due to the fact that these bases should occupy the peptidyl transferase center of the ribosome and should therefore be unable to form secondary structure while the ribosome decodes the slippery sequence.

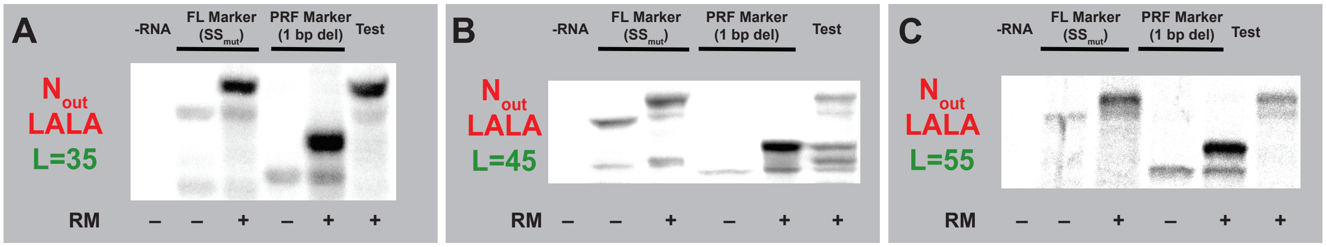


**Figure S2. Impact of Guest TMD Hydrophobicity on Programmed Ribosomal Frameshifting in LepB.** Representative SDS-PAGE gels depict the translation products generated by LepB frameshift reporter constructs bearing the LALA guest TMDs positioned either A) 35 codons, B) 45 codons, or C) 55 codons upstream of the slippery sequence. Lane 1 in each gel is a negative control containing no RNA. Full-Length (FL) molecular weight markers bear mutations within the slip-site and were generated in the absence (lane 2) and presence (lane 3) of rough microsomes. Ribosomal frameshift (RF) molecular weight markers in which the RF tail was moved to the 0-frame through the deletion of a single base pair were also generated in the absence (lane 4) and presence (lane 5) of microsomes. The test constructs for each spacer length are shown in lane 6 for each gel.

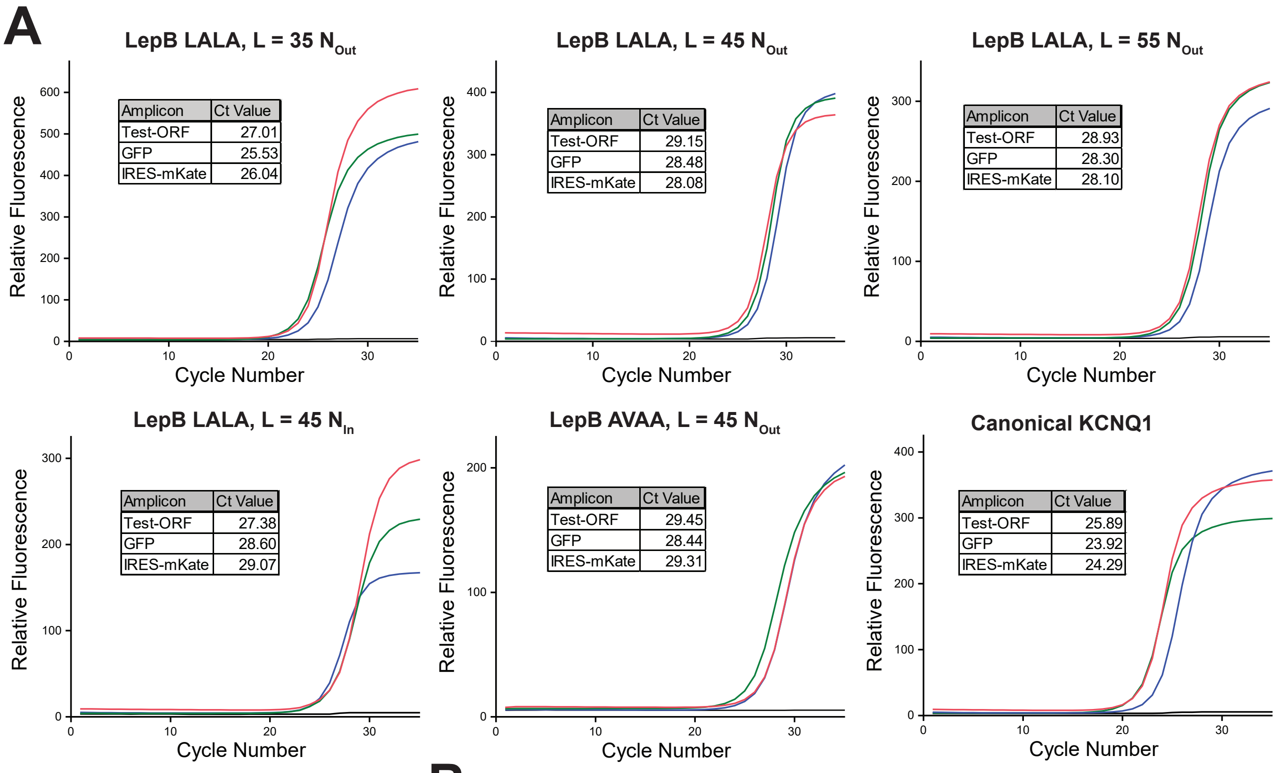


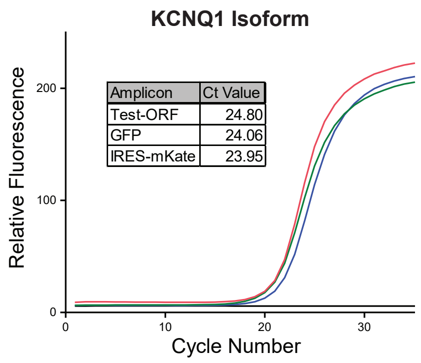


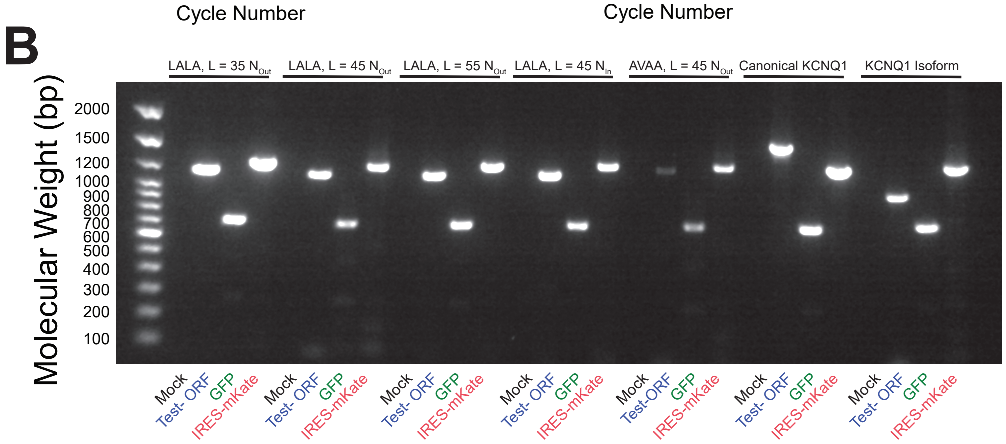


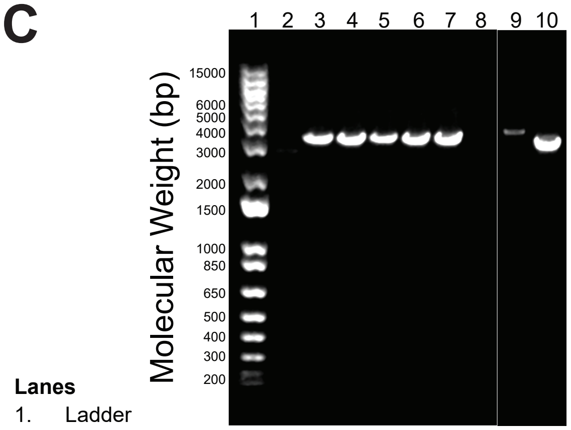


**Figure S3. Reporter Transcript Validation by qRT-PCR.** A series of fluorogenic -1PRF reporter constructs were transiently expressed in HEK293T cells prior to the extraction of cellular mRNA and the reverse transcription of the reporter transcript. A) Distinct regions within the reporter cDNA were quantified by qPCR in accordance with the MINDR guidelines. Scaled qPCR amplification curves along with the corresponding Ct values of each cassette are shown. B) An agarose gel shows the sizes of each qPCR product. Apmlicon sequences were validated by Next Generation Sequencing. C) A representative image of an agarose gel shows the relative size of the RT-PCR products of the full-length reporter constructs derived from cells expressing different reporters. The sequence of these products was validated by Sanger sequencing. Together, these results rule out spurious reporter signals that may arise from splicing artifacts.

**Lanes**

1. Ladder
2. Mock Transfection
3. LepB Fluorescent -1PRF Reporter (LALA, L = 35, $N_{out}$)
4. LepB Fluorescent -1PRF Reporter (LALA, L = 45, $N_{out}$)
5. LepB Fluorescent -1PRF Reporter (LALA, L = 45, $N_{in}$)
6. LepB Fluorescent -1PRF Reporter (LALA, L = 55, $N_{out}$)
7. LepB Fluorescent -1PRF Reporter (AVAA, L = 45, $N_{out}$)
8. Mock Transfection
9. Canonical KCNQ1 Fluorescent -1PRF Reporter
10. KCNQ1 Isoform Fluorescent -1PRF Reporter

### Table S1. Ribosomal Frameshifting Measurements for Modified LepB Constructs

| Construct | Guest TMD | Guest TMD Orientation | Spacer Length (codons) | *In Vitro* -1PRF Efficiency | Cellular -1PRF Efficiency |
|---|---|---|---|---|---|
| 1 | AVAA | $N_{out}$ | 35 | ND | - |
| 2 | AVAA | $N_{out}$ | 45 | 46 ± 8% | 2.1 ± 0.6 |
| 3 | AVAA | $N_{out}$ | 55 | ND* | - |
| 4 | AVAA | $N_{in}$ | 45 | 51 ± 3% | - |
| 5 | LALA | $N_{out}$ | 35 | ND* | 1.2 ± 0.1 |
| 6 | LALA | $N_{out}$ | 45 | 52 ± 4% | 2.3 ± 0.6 |
| 7 | LALA | $N_{out}$ | 55 | ND* | 2.1 ± 0.1 |
| 8 | LALA | $N_{in}$ | 45 | 50 ± 1% | 3.5 ± 0.5 |

PRF measurements represent the average of three replicates ± SD.
*Could not be reliably determined due to the low abundance of the frameshift products.

### Table S2. Impact of Pathogenic Mutations that Enhance KCNQ1 Misfolding on -1PRF

| KCNQ1 Variant | Location of Mutation | Median GFP / Median mKate Ratio* |
|---|---|---|
| WT | - | 0.368 ± 0.024 |
| $SS_{mut}$ | Slip Site | 0.240 ± 0.037 |
| Mut1 (G229D) | S4 | 0.382 ± 0.032 |
| Mut2 (L236P) | S4 | 0.361 ± 0.029 |
| Mut3 (I235N) | S4 | 0.408 ± 0.018 |
| Mut4 (W248R) | S4 | 0.378 ± 0.020 |
| Mut5 (L273R) | S5 | 0.383 ± 0.025 |
| Mut6 (V280E) | S5 | 0.372 ± 0.023 |
| Mut7 (L266R) | S5 | 0.390 ± 0.056 |
| Mut8 (L342H) | S6 | 0.388 ± 0.057 |
| Mut9 (A341E) | S6 | 0.378 ± 0.020 |
| Mut10 (G345R) | S6 | 0.377 ± 0.034 |

*Values represent the average of the median values across three biological replicates ± the standard deviation.

Measurements were carried out for each mutant in HEK293T cells using the natively spliced KCNQ1 -1PRF reporter. We chose mutations that were previously shown to induce misfolding that fall within the downstream TMDs of the KCNQ1 channel protein (see Ref. X).

### Supplemental Theory

*A Thermodynamic Approximation for the Resistance to -1 Ribosomal Frameshifting at Arbitrary Heptads*

Slippery heptanucleotide sequences represent one of the key features of efficient -1 programmed ribosomal frameshifting (-1PRF) motifs.[1–3] Generally, there are 24 well-characterized heptamers that correspond to well-characterized canonical $X_1$ $XXY_4$ $YYZ_7$ slippery sequences, in which X can be any of the four nucleotides (A, G, C, U/T), Y can be A or U only, and Z can be A, C, or U (not G).[4] However, there are other non-canonical heptamers that are also generally capable of contributing to an enhanced propensity for -1 ribosomal frameshifting.[3] Previous investigations have demonstrated that the contribution of individual "slippery" heptanucleotide sites to the net efficiency of a -1PRF site can be described in terms of the free energy difference associated with the A- and P-site t-RNA base pairing interactions in the -1 and 0- reading frames.[5] Based on this principle, we developed a python-based tool to scan through transcripts and "score" the resistance of successive heptads to -1 frameshifting according to the base pairing free energy differences in the two reading frames. In the following, we outline the logic of this computational approach.

We first developed a simple model to score the free energy difference associated with the binding of codons to their anticodons in the 0 and -1 reading frames in the PTC. To this end, we utilized Turner nearest neighbor energy parameters to score the energy of each individual base pairing interaction in the context of each frame.[6,7]. For each heptamer, we first reconstructed the corresponding codon sequences in both the 0 and -1 frames, focusing on the interactions between the tRNA anticodons and mRNA codons in the P-site and A-site of the ribosome. We then calculated the total base-pairing energy for each frame by considering Watson-Crick pairs, wobble pairs, and mismatches, as well as stacking interactions between adjacent nucleotide pairs, as defined by the Turner energy rules.[7] The energetic contributions include penalties for terminal mismatches and initiation, as well as adjustments for specific nearest-neighbor interactions and known thermodynamic properties of nucleotide pairings.

Using this energy scale, we calculated the free energy difference between the codon-anticodon base pairing energies in the 0 and -1 reading frame ($\Delta G_{FS}$) for all 16,384 possible heptameric sequences. Free energy differences calculated in this manner have units of kcal/ mol. A $\Delta G_{FS}$ value of 0 kcal/ mol corresponds to a heptad in which the base pairing interactions in the 0 and -1 reading frame are isoenergetic. Higher, positive scores indicate that a transition to the -1-reading frame incurs a net energetic penalty associated with unfavorable base pairing interactions. We used a python-based implementation of this algorithm to first evaluate $\Delta G_{FS}$ values for a set of 24 "canonical" slippery $X_1$ $XXY_4$ $YYZ_7$ heptads in which X can be any base, Y can be A or T, and Z can be A,C, or T. The scores for these motifs range from 0 to +2.9 kcal/ mol (see Doc. S2). Based on these benchmarks, we identified a comprehensive set of 465 heptamers that also fall within this energetic regime and used these as the basis of our search for TMD-slip motifs within the human transcriptome.

*Identification of Regions within Nascent Chains that are Likely to be Recognized by Translocons*

Though modern structure-prediction methodologies have provided comprehensive collections of molecular models for the three-dimensional structures of all human membrane proteins,[8,9] it is well established that native transmembrane domains (TMD's) do not always correspond to the regions of nascent polypeptides that undergo translocon-mediated membrane integration.[10,11] Because efficient nascent chain-mediated ribosomal frameshifting requires kinetic coupling between the decoding of slippery sequences and the engagement of the translocon,[12,13] which only occurs at certain spacings between the nascent TMD and slip-site,[14,15] we sought to accurately determine the positions of segments that are engaged by translocons within arbitrary transcripts. We first identified general regions within transcripts that are likely to encode TMDs using TOPCONS2.[16] We then developed a custom python script to adjust the boundaries of these TMD regions to reflect the position that is most likely to be engaged by the translocon complex using knowledge-based energy potentials derived by von Heijne and White (ΔG predictor).[17,18]

To identify these segments from sequences within the human transcriptome, we first translated all protein-coding transcripts from the *Homo sapiens* Ensembl coding sequence (CDS) database into amino acid sequences. We then input these coding sequences into the TOPCONS2 web server in order to identify the regions that are likely to encode TMDs. For each predicted TMD, we then used a sliding window of varying lengths (from 16 to 25 amino acids) to scan a region of the sequence beginning 10 amino acids upstream of the predicted TMD and ending 10 amino acids downstream of the predicted TMD. At each position and window length, we calculate the total transfer free energy (ΔG) associated within the translocon-mediated membrane integration based on depth-dependent transfer free energies using a python-based implementation of the von Heijne delta G predictor.[18]

By systematically evaluating all possible segments within the adjustment range (16-25 amino acids), we identify the specific segment that minimizes the membrane integration energy, which corresponds to the position that is most likely to be engaged by the translocon complex. We note that this approach ensures that regions encoding nascent TMDs do not overlap excessively and maintain minimal loop lengths between TMDs to account for the biochemical constraints of topogenesis.

*An Algorithmic Search for TMD-Slip Motifs within the Human Transcriptome*

Based on the biochemical and cellular measurements reported herein, we developed a custom python script to identify a comprehensive set of putative TMD-slip motifs at the transcriptomic scale based on the positions of slippery heptads relative to regions encoding segments that are likely to be engaged by translocons. This computational pipeline begins by parsing and filtering the Ensembl CDS database to extract high-quality protein-coding transcripts. The script first extracts essential metadata from the CDS FASTA files such as Ensembl gene IDs, transcript IDs, gene names, and nucleotide sequences associated with each transcript. To ensure the sequences are suitable for analysis, the scripts then implement quality control steps to reject any sequences that lack certain essential features such as a 5' methionine codon (ATG), a total number of bases that is divisible by three, and a terminal stop codon (TAG, TAA, or TGA). Transcripts are then translated into amino acid sequences prior to the identification of TMD coordinates using the module described above. For each transcript, the script scans the sequence for heptamers (i.e. the last 7 nucleotides of a 3-codon sliding window) and, if the heptamer is one of the 465 slippery heptamers ($\Delta G_{FS}$ of ≤ 2.9), the TMD that is closest to a position 45 codons upstream of the heptamer is assigned to the heptad. Heptamer's assigned to TMD regions beginning 35 to 55 codons are counted as TMD-slip motifs. For cases in which a single TMD is indexed to multiple heptamers, we selected the heptamer that is closest to the ideal distance (45 codons) as the most likely candidate and eliminated the other motifs to avoid double-counting. Nevertheless, it is possible that the translocation of a TMD may enhance frameshifting at multiple adjacent heptamers. For this reason, Doc S3 contains both the complete list of TMD-slip motifs as well as the pared down list of top candidate TMD-slip motifs. For each slippery heptamer, the script evaluates several key factors such as the $\Delta G_{FS}$ of the heptad and the number of codons between the slippery site and TMD. The script also calculates GC content of the intervening sequence, which may influence mRNA secondary structure, as well as the number of consecutive codons that fall downstream of the putative slippery sequence in the -1 frame prior to the encounter of a -1 stop codon. Finally, our script also evaluates the sequence that falls within 5-63 bases downstream of the slippery sequence for the presence of predicted the mRNA secondary structures using the ViennaRNA suite.[19]

*Statistical Assessments of TMD-Slip Motifs within the Human Transcriptome*

To determine whether the positions of slippery heptamers relative to TMD-encoding regions is nonrandom in canonical transcripts, we performed a permutation test in which the position of each TMD region is shuffled without altering their length or the number of TMDs within each transcript. We then measured the distance separating each slippery heptamer from the corresponding TMD region that is nearest to the ideal distance of 45 codons across each version of the shuffled transcriptome. There are 864 slippery heptamers that lie exactly at the 45-codons from an upstream TMD within canonical human transcripts. By comparison we find an average of 791 such motifs across 16,384 shuffled transcriptome iterations. Moreover, the full distance distribution in the native transcriptome is shifted further from the ideal distance (mean distance = +49.88 codons) relative to those within the shuffled transcriptomes (mean = +48.13 codons, $Z = +9.78$, $p_z = 1.4 \times 10^{-22}$; $p_{perm} = 6.1 \times 10^{-5}$). Additionally, a binomial test suggests the increased prevalence of motifs with ideal spacing within the native transcriptome relative to the shuffled transcriptomes is statistically significant ($p = 2.9 \times 10^{-3}$) and represents a statistically significant enrichment of ideal motifs ($\chi^2$ test $p = 2.6 \times 10^{-3}$, odds ratio = 1.11). Similar results were observed across motifs with a wider range of distance values (binomial test $p = 8.9 \times 10^{-8}$, $\chi^2$ test $p = 8.7 \times 10^{-8}$, odds ratio = 1.08). Together, these results demonstrate that slippery heptamers are more likely to be proceeded by a TMD-encoding region at the optimal spacing than would be expected by random chance.

As a complementary approach, we evaluated canonical transcripts to determine whether *slippery* heptamers, specifically, fall within non-random regions of the transcript relative to those encoding TMDs. To test this, we compared the distribution of spacings between heptamers and TMD-encoding regions among distinct, randomized sets of heptamers. We carried out this analysis for 16,384 iterations, each of which contains a unique set of 465 heptamers selected from the set of all 16,384 possible heptamers, which were selected with a frequency proportional to their relative abundance in the canonical transcriptome. For each iteration, we then computed the distances of the selected heptamers relative to the upstream TMD that falls closest to the ideal spacing of 45 codons in the manner described above. Across these iterations, there is an average of 1,782 random heptamers that fall exactly at 45 codons downstream of a TMD-encoding region; approximately twice the number of slippery heptamers within this region (864). This observation suggests there is a pronounced depletion of slippery heptamers at positions that may enhance ribosomal frameshifting. Because the search criteria are bookended on at the 5' by the TMD-encoding region, the average distance among random sets of heptamers skews towards shorter distances (mean distance = 43.9 codons); distances can be very long but cannot be negative. Notably, the distribution of distances for slippery heptamers instead skews towards longer distances (mean distance = +49.9 codons; $Z = +50.0$, $p_z < 10^{-30}$, $p_{perm} = 6.1 \times 10^{-5}$). A binomial test suggests the difference in the number of motifs formed by slippery heptamers versus random heptamer that have a spacing of 45 codons is statistically significant ($p = 4.1 \times 10^{-55}$). This depletion of slippery motifs at a distance of 45 codons is also statistically significant according to a $\chi^2$ test ($p = 2.0 \times 10^{-48}$, odds ratio = 0.605). Differences are also statistically significant across the wider window we employed for our TMD-slip motif search (35-55 codons) according to both the binomial test ($p < 10^{-30}$) and $\chi^2$ test ($p < 10^{-30}$, odds ratio = 0.360). Together, these results demonstrate strong negative selection against slippery heptamers at the optimal spacing than would be expected by random chance.

Importantly, any potential inconsistencies in the interpretations of these tests can be rectified with a more detailed consideration of the precise nature of their underlying hypotheses. For instance, the results of the TMD-shuffling test demonstrate that slippery heptamers are significantly more likely to be preceded by a transmembrane helix at the precise 45-codon spacing than would be expected if TMDs were randomly positioned. However, it is important to note that slippery heptamers are relatively under-represented in these transcripts overall. Thus, one interpretation of this test may be that, in the rare instances in which slippery heptamers appear within these transcripts, they are more likely to be preceded by an optimally spaced TMD-encoding region. This test addresses a fundamentally different hypothesis than the randomized heptamer test, which controls for deviations in the relative abundances of heptamers and tests whether those that fall within the optimal distance(s) tend to be more or less slippery than would be expected by chance. Together, the two tests show that slippery heptamers are generally depleted downstream of TMDs overall, but that when slippery heptamers do occur, they are more likely to fall within the optimal positions relative to TMDs.

*Gene Ontology analysis*

To assess whether our pool of putative TMD-slip motifs may be generally involved in certain biochemical mechanisms, we surveyed the “molecular function” Gene Ontology (GO) terms associated with the transcripts containing putative TMD-slip motifs. We first collected comprehensive UniProt metadata for the human proteins that are encoded by these transcripts as well as their associated GO annotations across three main categories including biological processes, cellular components, and molecular functions from the gene ontology consortium. This dataset provides a foundational understanding of the functional roles and cellular localization of the proteins associated with our putative TMD-Slip motifs. To determine whether certain GO terms are overrepresented across our set of transcripts containing TMD-Slip motifs (L = 35-55), we used the GOATOOLs python library to perform an enrichment analysis with a Fisher's Exact Test.[20] To perform this test, we compared the incidence of specific GO terms among these transcripts to their incidence across the larger set of human transcripts containing at least one TMD-encoding region as predicted by TOPCONS2.[16] Briefly, for each GO term, we constructed a contingency table that compares the number of transcripts containing TMD-Slip motifs that are annotated with a certain GO term to the corresponding number in the wider set of TMD-encoding transcripts. The Fisher's Exact Test then computes a *p*-value that reflects the probability that the observed incidence of this GO term among TMD-Slip containing transcripts should arise by chance (see Doc. S4). Low *p*-values suggest specific GO terms has been statistically enriched or depleted among TMD-Slip containing transcripts. Similarly, positive values of the corresponding odds ratios reflect an enrichment for a specific term whereas negative odds ratios reflect a depletion of transcripts associated with that GO term. To account for multiple hypothesis testing and control the false discovery rate, we adjust these *p*-values using the Benjamini-Hochberg method. This statistical correction helps to minimize false positives and to ensure the robust identification of statistically enriched GO terms.
