## Supplementary material for "Feedback from the Nascent Chain Triggers Ribosomal Frameshifting and Transcript Decay": Doc S1 Reporter Sequences

**Document S1. Annotated Sequences of Genetic Frameshift Reporters.**

The DNA sequences for each reporter construct characterized in this work is shown below.

For *in vitro* LepB -1PRF reporters (numbers 1-24), we have highlighted the sequence of the guest TMD in **blue**, the sequence of the slippery sequence in **green**, the -1 stop codon resulting in the truncation of the PRF product in **purple**, and the 0-frame stop codon in **red**. Bases that were inserted to switch the reading frames are highlighted in **yellow**. Estimated molecular weights of -1PRF products reflect the size of the proteins generated by a -1 frameshift with two tRNAs loaded into the peptidyl transferase center. The region encoding the additional TMD that we inserted to flip the topology of the guest TMD in the context of the N_in_ reporters is highlighted in **green**.

For cellular LepB -1PRF reporters (numbers 25-37), we have highlighted the sequence of the guest TMD in **blue**, the sequence of the slippery sequence in **green**, the stop codon at the end of the -1 GFP reporter in **purple**, and the 0-frame stop codon in **red**. Bases that were inserted to switch the reading frames are highlighted in **yellow**. The region encoding the additional TMD that we inserted to flip the topology of the guest TMD in the context of the N_in_ reporters is highlighted in **gray**. The region encoding the -1 P2A linker is highlighted in **blue** and the -1 GFP is highlighted in **green**.

For all other cellular -1PRF reporters (numbers 38-62), we have highlighted the sequence of the TMD that is closest to the optimal distance from the slippery sequence in **blue**, the sequence of the slippery sequence in **green**, the stop codon at the end of the -1 GFP reporter in **purple**, and the 0-frame stop codon in **red**. Bases that were inserted to switch the reading frames are highlighted in **yellow**. The region encoding the -1 P2A linker is highlighted in **blue** and the -1 GFP is highlighted in **green**.

**1. LepB *in vitro* Translation Reporter (N_out_, AVAA TMD, L=35) (0-Frame = 44.1 kDa, PRF ~ 35.4 kDa)**

ATGGCGAATATGTTTGCCCTGATTCTGGTGATTGCCACACTGGTGACGGGCATTTTATGGTGCGTGGATAAATTCTTTTTCGCACCTAAACGGCGGGAACGTCAGGCAGCGGCGCAGGCGGCTGCCGGGGACTCACTGGATAAAGCAACGTTGAAAAAGGTTGCGCCGAAGCCTGGCTGGCTGGAAACCGGTGCTTCTGTTatgCCGGTACTGatgATCGTATTGATTGTGCGTTCGTTTATTTATGAACCGTTCCAGATCCCGTCAGGTTCGATGATGCCGACTCTGaactctactGATTTTATTCTGGTAGAGAAGTTTGCTTATGGCATgAAAGATCCTATgTACCAGAAAACGaTGATCGAAACCGGTCATCCGAAACGCGGCGATATCaTGatgTTTAAATATCCGGAAGATCCAAAGCTTGATTACATCAAGCGCGCGGTGGGTTTACCGGGCGATAAAGTCACTTACGATCCGGTCTCAAAAGAGaTGACGATgCAACCGGGATGCAGTTCCGGCCAGGCGTGTGAAAACGCGCTGCCGGTCACCTACTCAAACGTGGAACCGAGCGATTTCGTTCAGACCTTCTCACGCCGTAATGGTGGGGAAGCGACCAGCGGATTCTTTGAAGTGCCGAAACAGGAAACCAAAGAAAATGGAATTCGTCTTTCCGAGactagtggTggTccagg**TGCAGCAGCTgcTGTTGCAGCTgcTGCTGCTGCAgcAGCAGCCGTTgcTGCTGCAGC**TggTcctggcggTacggtaccAGGTCAACAAaatgcaGTTTGGATTGTTCCTCCTGGACAATACTTCATGATGGGCGACAACCGCGACAACAGCGCGGACAGCCG**TTTTTTT**GGCTTTGTGCCGGAAGCGAATCTGGTCGGTCGGGCAACGGCTATCTGGATGAGCTTCGAaAAGCAAGAAGGCGAATGGCCGACTGGTCTGCGCTTAAGTCGCATTGGCGGCATCCA**TGa**tGATCCATCTTCGTTCACGTTGTCGCCGTTATGGCGACCCGGATCCCCGGGCGAGCTCGAATTCCGGTCTCCCTATAGTGAGTCGTATTccTTTCGATccGCCAGCTGCATTcAgGAATCGGCCAACGCGCGGGGAGAGGCGGTTTGCGTATTGGGCGCTCTTCCGCTTCCTCGCTCACTGACTCGCTGCGCTCGGTCGTTCGGCTGCGGCGAGCGGTATCAGCTCACTCAAAGGCGG**TAA**

**2. Full-Length Marker, LepB *in vitro* Translation Reporter (N_out_, AVAA TMD, L=35) (0-Frame = 44.1 kDa)**

ATGGCGAATATGTTTGCCCTGATTCTGGTGATTGCCACACTGGTGACGGGCATTTTATGGTGCGTGGATAAATTCTTTTTCGCACCTAAACGGCGGGAACGTCAGGCAGCGGCGCAGGCGGCTGCCGGGGACTCACTGGATAAAGCAACGTTGAAAAAGGTTGCGCCGAAGCCTGGCTGGCTGGAAACCGGTGCTTCTGTTatgCCGGTACTGatgATCGTATTGATTGTGCGTTCGTTTATTTATGAACCGTTCCAGATCCCGTCAGGTTCGATGATGCCGACTCTGaactctactGATTTTATTCTGGTAGAGAAGTTTGCTTATGGCATgAAAGATCCTATgTACCAGAAAACGaTGATCGAAACCGGTCATCCGAAACGCGGCGATATCaTGatgTTTAAATATCCGGAAGATCCAAAGCTTGATTACATCAAGCGCGCGGTGGGTTTACCGGGCGATAAAGTCACTTACGATCCGGTCTCAAAAGAGaTGACGATgCAACCGGGATGCAGTTCCGGCCAGGCGTGTGAAAACGCGCTGCCGGTCACCTACTCAAACGTGGAACCGAGCGATTTCGTTCAGACCTTCTCACGCCGTAATGGTGGGGAAGCGACCAGCGGATTCTTTGAAGTGCCGAAACAGGAAACCAAAGAAAATGGAATTCGTCTTTCCGAGactagtggTggTccaggT**GCAGCAGCTgcTGTTGCAGCTgcTGCTGCTGCAgcAGCAGCCGTTgcTGCTGCAGC**TggTcctggcggTacggtaccAGGTCAACAAaatgcaGTTTGGATTGTTCCTCCTGGACAATACTTCATGATGGGCGACAACCGCGACAACAGCGCGGACAGCCG**GTTCTTC**GGCTTTGTGCCGGAAGCGAATCTGGTCGGTCGGGCAACGGCTATCTGGATGAGCTTCGAaAAGCAAGAAGGCGAATGGCCGACTGGTCTGCGCTTAAGTCGCATTGGCGGCATCCATGatGATCCATCTTCGTTCACGTTGTCGCCGTTATGGCGACCCGGATCCCCGGGCGAGCTCGAATTCCGGTCTCCCTATAGTGAGTCGTATTccTTTCGATccGCCAGCTGCATTcAgGAATCGGCCAACGCGCGGGGAGAGGCGGTTTGCGTATTGGGCGCTCTTCCGCTTCCTCGCTCACTGACTCGCTGCGCTCGGTCGTTCGGCTGCGGCGAGCGGTATCAGCTCACTCAAAGGCGG**TAA**

***3.* PRF Marker, LepB *in vitro* Translation Reporter (N_out_, AVAA TMD, L=35) (0-Frame = 35.5 kDa)**

ATGGCGAATATGTTTGCCCTGATTCTGGTGATTGCCACACTGGTGACGGGCATTTTATGGTGCGTGGATAAATTCTTTTTCGCACCTAAACGGCGGGAACGTCAGGCAGCGGCGCAGGCGGCTGCCGGGGACTCACTGGATAAAGCAACGTTGAAAAAGGTTGCGCCGAAGCCTGGCTGGCTGGAAACCGGTGCTTCTGTTatgCCGGTACTGatgATCGTATTGATTGTGCGTTCGTTTATTTATGAACCGTTCCAGATCCCGTCAGGTTCGATGATGCCGACTCTGaactctactGATTTTATTCTGGTAGAGAAGTTTGCTTATGGCATgAAAGATCCTATgTACCAGAAAACGaTGATCGAAACCGGTCATCCGAAACGCGGCGATATCaTGatgTTTAAATATCCGGAAGATCCAAAGCTTGATTACATCAAGCGCGCGGTGGGTTTACCGGGCGATAAAGTCACTTACGATCCGGTCTCAAAAGAGaTGACGATgCAACCGGGATGCAGTTCCGGCCAGGCGTGTGAAAACGCGCTGCCGGTCACCTACTCAAACGTGGAACCGAGCGATTTCGTTCAGACCTTCTCACGCCGTAATGGTGGGGAAGCGACCAGCGGATTCTTTGAAGTGCCGAAACAGGAAACCAAAGAAAATGGAATTCGTCTTTCCGAGactagtggTggTccaggT**GCAGCAGCTgcTGTTGCAGCTgcTGCTGCTGCAgcAGCAGCCGTTgcTGCTGCAGC**TggTcctggcggTacggtaccAGGTCAACAAaatgcaGTTTGGATTGTTCCTCCTGGACAATACTTCATGATGGGCGACAACCGCGACAACAGCGCGGACAGCCG**GTTCTTCT**GGCTTTGTGCCGGAAGCGAATCTGGTCGGTCGGGCAACGGCTATCTGGATGAGCTTCGAaAAGCAAGAAGGCGAATGGCCGACTGGTCTGCGCTTAAGTCGCATTGGCGGCATCCA**TGa**tGATCCATCTTCGTTCACGTTGTCGCCGTTATGGCGACCCGGATCCCCGGGCGAGCTCGAATTCCGGTCTCCCTATAGTGAGTCGTATTccTTTCGATccGCCAGCTGCATTcAgGAATCGGCCAACGCGCGGGGAGAGGCGGTTTGCGTATTGGGCGCTCTTCCGCTTCCTCGCTCACTGACTCGCTGCGCTCGGTCGTTCGGCTGCGGCGAGCGGTATCAGCTCACTCAAAGGCGGTAA

**4. LepB *in vitro* Translation Reporter (N_out_, AVAA TMD, L=45) (0-Frame = 44.3 kDa, PRF ~ 35.6 kDa)**

ATGGCGAATATGTTTGCCCTGATTCTGGTGATTGCCACACTGGTGACGGGCATTTTATGGTGCGTGGATAAATTCTTTTTCGCACCTAAACGGCGGGAACGTCAGGCAGCGGCGCAGGCGGCTGCCGGGGACTCACTGGATAAAGCAACGTTGAAAAAGGTTGCGCCGAAGCCTGGCTGGCTGGAAACCGGTGCTTCTGTTatgCCGGTACTGatgATCGTATTGATTGTGCGTTCGTTTATTTATGAACCGTTCCAGATCCCGTCAGGTTCGATGATGCCGACTCTGaactctactGATTTTATTCTGGTAGAGAAGTTTGCTTATGGCATgAAAGATCCTATgTACCAGAAAACGaTGATCGAAACCGGTCATCCGAAACGCGGCGATATCaTGatgTTTAAATATCCGGAAGATCCAAAGCTTGATTACATCAAGCGCGCGGTGGGTTTACCGGGCGATAAAGTCACTTACGATCCGGTCTCAAAAGAGaTGACGATgCAACCGGGATGCAGTTCCGGCCAGGCGTGTGAAAACGCGCTGCCGGTCACCTACTCAAACGTGGAACCGAGCGATTTCGTTCAGACCTTCTCACGCCGTAATGGTGGGGAAGCGACCAGCGGATTCTTTGAAGTGCCGAAACAGGAAACCAAAGAAAATGGAATTCGTCTTTCCGAGactagtggTggTccaggT**GCAGCAGCTgcTGTTGCAGCTgcTGCTGCTGCAgcAGCAGCCGTTgcTGCTGCAGC**TggTcctggcggTacggtaccAGGTCAACAAaatgcaGTTTGGATTGTTCCTCCTGGACAATACTTCATGATGGGCGACAACCGCGACAACAGCGCGGACAGCCGTTACTGGGGCTTTGTGCCGGAAGCGAATC**TTTTTTT**TCGGGCAACGGCTATCTGGATGAGCTTCGAaAAGCAAGAAGGCGAATGGCCGACTGGTCTGCGCTTAAGTCGCATTGGCGGCATCCA**TGa**tGATCCATCTTCGTTCACGTTGTCGCCGTTATGGCGACCCGGATCCCCGGGCGAGCTCGAATTCCGGTCTCCCTATAGTGAGTCGTATTccTTTCGATccGCCAGCTGCATTcAgGAATCGGCCAACGCGCGGGGAGAGGCGGTTTGCGTATTGGGCGCTCTTCCGCTTCCTCGCTCACTGACTCGCTGCGCTCGGTCGTTCGGCTGCGGCGAGCGGTATCAGCTCACTCAAAGGCGG**TAA**

**5. Full-Length Marker, LepB *in vitro* Translation Reporter (N_out_, AVAA TMD, L=45) (0-Frame = 44.3 kDa)**

ATGGCGAATATGTTTGCCCTGATTCTGGTGATTGCCACACTGGTGACGGGCATTTTATGGTGCGTGGATAAATTCTTTTTCGCACCTAAACGGCGGGAACGTCAGGCAGCGGCGCAGGCGGCTGCCGGGGACTCACTGGATAAAGCAACGTTGAAAAAGGTTGCGCCGAAGCCTGGCTGGCTGGAAACCGGTGCTTCTGTTaTgCCGGTACTGatgATCGTATTGATTGTGCGTTCGTTTATTTATGAACCGTTCCAGATCCCGTCAGGTTCGATGATGCCGACTCTGaactctactGATTTTATTCTGGTAGAGAAGTTTGCTTATGGCATgAAAGATCCTATgTACCAGAAAACGaTGATCGAAACCGGTCATCCGAAACGCGGCGATATCaTGaTgTTTAAATATCCGGAAGATCCAAAGCTTGATTACATCAAGCGCGCGGTGGGTTTACCGGGCGATAAAGTCACTTACGATCCGGTCTCAAAAGAGaTGACGATgCAACCGGGATGCAGTTCCGGCCAGGCGTGTGAAAACGCGCTGCCGGTCACCTACTCAAACGTGGAACCGAGCGATTTCGTTCAGACCTTCTCACGCCGTAATGGTGGGGAAGCGACCAGCGGATTCTTTGAAGTGCCGAAACAGGAAACCAAAGAAAATGGAATTCGTCTTTCCGAGactagtggTggTccaggT**GCAGCAGCTgcTGTTGCAGCTgcTGCTGCTGCAgcAGCAGCCGTTgcTGCTGCAGC**TggTcctggcggTacggtaccAGGTCAACAAaatgcaGTTTGGATTGTTCCTCCTGGACAATACTTCATGATGGGCGACAACCGCGACAACAGCGCGGACAGCCGTTACTGGGGCTTTGTGCCGGAAGCGAATC**TGTTCTTC**CGGGCAACGGCTATCTGGATGAGCTTCGAaAAGCAAGAAGGCGAATGGCCGACTGGTCTGCGCTTAAGTCGCATTGGCGGCATCCATGatGATCCATCTTCGTTCACGTTGTCGCCGTTATGGCGACCCGGATCCCCGGGCGAGCTCGAATTCCGGTCTCCCTATAGTGAGTCGTATTccTTTCGATccGCCAGCTGCATTcAgGAATCGGCCAACGCGCGGGGAGAGGCGGTTTGCGTATTGGGCGCTCTTCCGCTTCCTCGCTCACTGACTCGCTGCGCTCGGTCGTTCGGCTGCGGCGAGCGGTATCAGCTCACTCAAAGGCGG**TAA**

**6. PRF Marker, LepB *in vitro* Translation Reporter (N_out_, AVAA TMD, L=45) (0-Frame = 35.6 kDa)**

ATGGCGAATATGTTTGCCCTGATTCTGGTGATTGCCACACTGGTGACGGGCATTTTATGGTGCGTGGATAAATTCTTTTTCGCACCTAAACGGCGGGAACGTCAGGCAGCGGCGCAGGCGGCTGCCGGGGACTCACTGGATAAAGCAACGTTGAAAAAGGTTGCGCCGAAGCCTGGCTGGCTGGAAACCGGTGCTTCTGTTatgCCGGTACTGatgATCGTATTGATTGTGCGTTCGTTTATTTATGAACCGTTCCAGATCCCGTCAGGTTCGATGATGCCGACTCTGaactctactGATTTTATTCTGGTAGAGAAGTTTGCTTATGGCATgAAAGATCCTATgTACCAGAAAACGaTGATCGAAACCGGTCATCCGAAACGCGGCGATATCaTGatgTTTAAATATCCGGAAGATCCAAAGCTTGATTACATCAAGCGCGCGGTGGGTTTACCGGGCGATAAAGTCACTTACGATCCGGTCTCAAAAGAGaTGACGATgCAACCGGGATGCAGTTCCGGCCAGGCGTGTGAAAACGCGCTGCCGGTCACCTACTCAAACGTGGAACCGAGCGATTTCGTTCAGACCTTCTCACGCCGTAATGGTGGGGAAGCGACCAGCGGATTCTTTGAAGTGCCGAAACAGGAAACCAAAGAAAATGGAATTCGTCTTTCCGAGactagtggTggTccaggT**GCAGCAGCTgcTGTTGCAGCTgcTGCTGCTGCAgcAGCAGCCGTTgcTGCTGCAGC**TggTcctggcggTacggtaccAGGTCAACAAaatgcaGTTTGGATTGTTCCTCCTGGACAATACTTCATGATGGGCGACAACCGCGACAACAGCGCGGACAGCCGTTACTGGGGCTTTGTGCCGGAAGCGAATC**TGTTCTTCT**CGGGCAACGGCTATCTGGATGAGCTTCGAaAAGCAAGAAGGCGAATGGCCGACTGGTCTGCGCTTAAGTCGCATTGGCGGCATCCA**TGa**tGATCCATCTTCGTTCACGTTGTCGCCGTTATGGCGACCCGGATCCCCGGGCGAGCTCGAATTCCGGTCTCCCTATAGTGAGTCGTATTccTTTCGATccGCCAGCTGCATTcAgGAATCGGCCAACGCGCGGGGAGAGGCGGTTTGCGTATTGGGCGCTCTTCCGCTTCCTCGCTCACTGACTCGCTGCGCTCGGTCGTTCGGCTGCGGCGAGCGGTATCAGCTCACTCAAAGGCGGTAA

**7. LepB *in vitro* Translation Reporter (N_out_, AVAA TMD, L=55) (0-Frame = 44.1 kDa, PRF ~ 35.6 kDa)**

ATGGCGAATATGTTTGCCCTGATTCTGGTGATTGCCACACTGGTGACGGGCATTTTATGGTGCGTGGATAAATTCTTTTTCGCACCTAAACGGCGGGAACGTCAGGCAGCGGCGCAGGCGGCTGCCGGGGACTCACTGGATAAAGCAACGTTGAAAAAGGTTGCGCCGAAGCCTGGCTGGCTGGAAACCGGTGCTTCTGTTatgCCGGTACTGatgATCGTATTGATTGTGCGTTCGTTTATTTATGAACCGTTCCAGATCCCGTCAGGTTCGATGATGCCGACTCTGaactctactGATTTTATTCTGGTAGAGAAGTTTGCTTATGGCATgAAAGATCCTATgTACCAGAAAACGaTGATCGAAACCGGTCATCCGAAACGCGGCGATATCaTGatgTTTAAATATCCGGAAGATCCAAAGCTTGATTACATCAAGCGCGCGGTGGGTTTACCGGGCGATAAAGTCACTTACGATCCGGTCTCAAAAGAGaTGACGATgCAACCGGGATGCAGTTCCGGCCAGGCGTGTGAAAACGCGCTGCCGGTCACCTACTCAAACGTGGAACCGAGCGATTTCGTTCAGACCTTCTCACGCCGTAATGGTGGGGAAGCGACCAGCGGATTCTTTGAAGTGCCGAAACAGGAAACCAAAGAAAATGGAATTCGTCTTTCCGAGactagtggTggTccaggT**GCAGCAGCTgcTGTTGCAGCTgcTGCTGCTGCAgcAGCAGCCGTTgcTGCTGCAGC**TggTcctggcggTacggtaccAGGTCAACAAaatgcaGTTTGGATTGTTCCTCCTGGACAATACTTCATGATGGGCGACAACCGCGACAACAGCGCGGACAGCCGTTACTGGGGCTTTGTGCCGGAAGCGAATCTGGTCGGTCGGGCAACGGCTATCTGGATGAG**TTTTTTT**AcGCAAGAAGGCGAATGGCCGACTGGTCTGCGCTTAAGTCGCATTGGCGGCATCCA**TGa**tGATCCATCTTCGTTCACGTTGTCGCCGTTATGGCGACCCGGATCCCCGGGCGAGCTCGAATTCCGGTCTCCCTATAGTGAGTCGTATTccTTTCGATccGCCAGCTGCATTcAgGAATCGGCCAACGCGCGGGGAGAGGCGGTTTGCGTATTGGGCGCTCTTCCGCTTCCTCGCTCACTGACTCGCTGCGCTCGGTCGTTCGGCTGCGGCGAGCGGTATCAGCTCACTCAAAGGCGG**TAA**

**8. Full-Length Marker, LepB *in vitro* Translation Reporter (N_out_, AVAA TMD, L=55) (0-Frame = 44.1 kDa)**

ATGGCGAATATGTTTGCCCTGATTCTGGTGATTGCCACACTGGTGACGGGCATTTTATGGTGCGTGGATAAATTCTTTTTCGCACCTAAACGGCGGGAACGTCAGGCAGCGGCGCAGGCGGCTGCCGGGGACTCACTGGATAAAGCAACGTTGAAAAAGGTTGCGCCGAAGCCTGGCTGGCTGGAAACCGGTGCTTCTGTTatgCCGGTACTGatgATCGTATTGATTGTGCGTTCGTTTATTTATGAACCGTTCCAGATCCCGTCAGGTTCGATGATGCCGACTCTGaactctactGATTTTATTCTGGTAGAGAAGTTTGCTTATGGCATgAAAGATCCTATgTACCAGAAAACGaTGATCGAAACCGGTCATCCGAAACGCGGCGATATCaTGatgTTTAAATATCCGGAAGATCCAAAGCTTGATTACATCAAGCGCGCGGTGGGTTTACCGGGCGATAAAGTCACTTACGATCCGGTCTCAAAAGAGaTGACGATgCAACCGGGATGCAGTTCCGGCCAGGCGTGTGAAAACGCGCTGCCGGTCACCTACTCAAACGTGGAACCGAGCGATTTCGTTCAGACCTTCTCACGCCGTAATGGTGGGGAAGCGACCAGCGGATTCTTTGAAGTGCCGAAACAGGAAACCAAAGAAAATGGAATTCGTCTTTCCGAGactagtggTggTccaggT**GCAGCAGCTgcTGTTGCAGCTgcTGCTGCTGCAgcAGCAGCCGTTgcTGCTGCAGC**TggTcctggcggTacggtaccAGGTCAACAAaatgcaGTTTGGATTGTTCCTCCTGGACAATACTTCATGATGGGCGACAACCGCGACAACAGCGCGGACAGCCGTTACTGGGGCTTTGTGCCGGAAGCGAATCTGGTCGGTCGGGCAACGGCTATCTGGATG**TCGTTCTTC**AcGCAAGAAGGCGAATGGCCGACTGGTCTGCGCTTAAGTCGCATTGGCGGCATCCATGatGATCCATCTTCGTTCACGTTGTCGCCGTTATGGCGACCCGGATCCCCGGGCGAGCTCGAATTCCGGTCTCCCTATAGTGAGTCGTATTccTTTCGATccGCCAGCTGCATTcAgGAATCGGCCAACGCGCGGGGAGAGGCGGTTTGCGTATTGGGCGCTCTTCCGCTTCCTCGCTCACTGACTCGCTGCGCTCGGTCGTTCGGCTGCGGCGAGCGGTATCAGCTCACTCAAAGGCGG**TAA**

**9. PRF Marker, LepB *in vitro* Translation Reporter (N_out_, AVAA TMD, L=55) (0-Frame = 35.6 kDa)**

ATGGCGAATATGTTTGCCCTGATTCTGGTGATTGCCACACTGGTGACGGGCATTTTATGGTGCGTGGATAAATTCTTTTTCGCACCTAAACGGCGGGAACGTCAGGCAGCGGCGCAGGCGGCTGCCGGGGACTCACTGGATAAAGCAACGTTGAAAAAGGTTGCGCCGAAGCCTGGCTGGCTGGAAACCGGTGCTTCTGTTatgCCGGTACTGatgATCGTATTGATTGTGCGTTCGTTTATTTATGAACCGTTCCAGATCCCGTCAGGTTCGATGATGCCGACTCTGaactctactGATTTTATTCTGGTAGAGAAGTTTGCTTATGGCATgAAAGATCCTATgTACCAGAAAACGaTGATCGAAACCGGTCATCCGAAACGCGGCGATATCaTGatgTTTAAATATCCGGAAGATCCAAAGCTTGATTACATCAAGCGCGCGGTGGGTTTACCGGGCGATAAAGTCACTTACGATCCGGTCTCAAAAGAGaTGACGATgCAACCGGGATGCAGTTCCGGCCAGGCGTGTGAAAACGCGCTGCCGGTCACCTACTCAAACGTGGAACCGAGCGATTTCGTTCAGACCTTCTCACGCCGTAATGGTGGGGAAGCGACCAGCGGATTCTTTGAAGTGCCGAAACAGGAAACCAAAGAAAATGGAATTCGTCTTTCCGAGactagtggTggTccaggT**GCAGCAGCTgcTGTTGCAGCTgcTGCTGCTGCAgcAGCAGCCGTTgcTGCTGCAGC**TggTcctggcggTacggtaccAGGTCAACAAaatgcaGTTTGGATTGTTCCTCCTGGACAATACTTCATGATGGGCGACAACCGCGACAACAGCGCGGACAGCCGTTACTGGGGCTTTGTGCCGGAAGCGAATCTGGTCGGTCGGGCAACGGCTATCTGGATG**TCGTTCTTCT**AcGCAAGAAGGCGAATGGCCGACTGGTCTGCGCTTAAGTCGCATTGGCGGCATCCA**TGa**tGATCCATCTTCGTTCACGTTGTCGCCGTTATGGCGACCCGGATCCCCGGGCGAGCTCGAATTCCGGTCTCCCTATAGTGAGTCGTATTccTTTCGATccGCCAGCTGCATTcAgGAATCGGCCAACGCGCGGGGAGAGGCGGTTTGCGTATTGGGCGCTCTTCCGCTTCCTCGCTCACTGACTCGCTGCGCTCGGTCGTTCGGCTGCGGCGAGCGGTATCAGCTCACTCAAAGGCGGTAA

**10. LepB *in vitro* Translation Reporter (N_out_, LALA TMD, L=35) (0-Frame = 44.3 kDa, PRF ~ 36.7 kDa)**

ATGGCGAATATGTTTGCCCTGATTCTGGTGATTGCCACACTGGTGACGGGCATTTTATGGTGCGTGGATAAATTCTTTTTCGCACCTAAACGGCGGGAACGTCAGGCAGCGGCGCAGGCGGCTGCCGGGGACTCACTGGATAAAGCAACGTTGAAAAAGGTTGCGCCGAAGCCTGGCTGGCTGGAAACCGGTGCTTCTGTTatgCCGGTACTGatgATCGTATTGATTGTGCGTTCGTTTATTTATGAACCGTTCCAGATCCCGTCAGGTTCGATGATGCCGACTCTGaactctactGATTTTATTCTGGTAGAGAAGTTTGCTTATGGCATgAAAGATCCTATgTACCAGAAAACGaTGATCGAAACCGGTCATCCGAAACGCGGCGATATCaTGatgTTTAAATATCCGGAAGATCCAAAGCTTGATTACATCAAGCGCGCGGTGGGTTTACCGGGCGATAAAGTCACTTACGATCCGGTCTCAAAAGAGaTGACGATgCAACCGGGATGCAGTTCCGGCCAGGCGTGTGAAAACGCGCTGCCGGTCACCTACTCAAACGTGGAACCGAGCGATTTCGTTCAGACCTTCTCACGCCGTAATGGTGGGGAAGCGACCAGCGGATTCTTTGAAGTGCCGAAACAGGAAACCAAAGAAAATGGAATTCGTCTTTCCGAGactagtGGTGGTCCAGGT**GCTCTGGCAGCTCTGGCACTGGCTGCATTAGCCGCTCTGGCCCTGGCCGCCTTAGCC**GGTCCTGGCGGTacggtaccAGGTCAACAAaatgcaGTTTGGATTGTTCCTCCTGGACAATACTTCATGATGGGCGACAACCGCGACAACAGCGCGGACAGCCG**TTTTTTT**GGCTTTGTGCCGGAAGCGAATCTGGTCGGTCGGGCAACGGCTATCTGGATGAGCTTCGAaAAGCAAGAAGGCGAATGGCCGACTGGTCTGCGCTTAAGTCGCATTGGCGGCATCCA**TGa**tGATCCATCTTCGTTCACGTTGTCGCCGTTATGGCGACCCGGATCCCCGGGCGAGCTCGAATTCCGGTCTCCCTATAGTGAGTCGTATTccTTTCGATccGCCAGCTGCATTcAgGAATCGGCCAACGCGCGGGGAGAGGCGGTTTGCGTATTGGGCGCTCTTCCGCTTCCTCGCTCACTGACTCGCTGCGCTCGGTCGTTCGGCTGCGGCGAGCGGTATCAGCTCACTCAAAGGCGG**TAA**

**11. Full-Length Marker, LepB *in vitro* Translation Reporter (N_out_, LALA TMD, L=35) (0-Frame = 44.3 kDa)**

ATGGCGAATATGTTTGCCCTGATTCTGGTGATTGCCACACTGGTGACGGGCATTTTATGGTGCGTGGATAAATTCTTTTTCGCACCTAAACGGCGGGAACGTCAGGCAGCGGCGCAGGCGGCTGCCGGGGACTCACTGGATAAAGCAACGTTGAAAAAGGTTGCGCCGAAGCCTGGCTGGCTGGAAACCGGTGCTTCTGTTatgCCGGTACTGatgATCGTATTGATTGTGCGTTCGTTTATTTATGAACCGTTCCAGATCCCGTCAGGTTCGATGATGCCGACTCTGaactctactGATTTTATTCTGGTAGAGAAGTTTGCTTATGGCATgAAAGATCCTATgTACCAGAAAACGaTGATCGAAACCGGTCATCCGAAACGCGGCGATATCaTGatgTTTAAATATCCGGAAGATCCAAAGCTTGATTACATCAAGCGCGCGGTGGGTTTACCGGGCGATAAAGTCACTTACGATCCGGTCTCAAAAGAGaTGACGATgCAACCGGGATGCAGTTCCGGCCAGGCGTGTGAAAACGCGCTGCCGGTCACCTACTCAAACGTGGAACCGAGCGATTTCGTTCAGACCTTCTCACGCCGTAATGGTGGGGAAGCGACCAGCGGATTCTTTGAAGTGCCGAAACAGGAAACCAAAGAAAATGGAATTCGTCTTTCCGAGactagtGGTGGTCCAGGT**GCTCTGGCAGCTCTGGCACTGGCTGCATTAGCCGCTCTGGCCCTGGCCGCCTTAGCC**GGTCCTGGCGGTacggtaccAGGTCAACAAaatgcaGTTTGGATTGTTCCTCCTGGACAATACTTCATGATGGGCGACAACCGCGACAACAGCGCGGACAGCCG**GTTCTTC**GGCTTTGTGCCGGAAGCGAATCTGGTCGGTCGGGCAACGGCTATCTGGATGAGCTTCGAaAAGCAAGAAGGCGAATGGCCGACTGGTCTGCGCTTAAGTCGCATTGGCGGCATCCATGatGATCCATCTTCGTTCACGTTGTCGCCGTTATGGCGACCCGGATCCCCGGGCGAGCTCGAATTCCGGTCTCCCTATAGTGAGTCGTATTccTTTCGATccGCCAGCTGCATTcAgGAATCGGCCAACGCGCGGGGAGAGGCGGTTTGCGTATTGGGCGCTCTTCCGCTTCCTCGCTCACTGACTCGCTGCGCTCGGTCGTTCGGCTGCGGCGAGCGGTATCAGCTCACTCAAAGGCGG**TAA**

**12. PRF Marker, LepB *in vitro* Translation Reporter (N_out_, LALA TMD, L=35) (0-Frame = 35.7 kDa)**

ATGGCGAATATGTTTGCCCTGATTCTGGTGATTGCCACACTGGTGACGGGCATTTTATGGTGCGTGGATAAATTCTTTTTCGCACCTAAACGGCGGGAACGTCAGGCAGCGGCGCAGGCGGCTGCCGGGGACTCACTGGATAAAGCAACGTTGAAAAAGGTTGCGCCGAAGCCTGGCTGGCTGGAAACCGGTGCTTCTGTTatgCCGGTACTGatgATCGTATTGATTGTGCGTTCGTTTATTTATGAACCGTTCCAGATCCCGTCAGGTTCGATGATGCCGACTCTGaactctactGATTTTATTCTGGTAGAGAAGTTTGCTTATGGCATgAAAGATCCTATgTACCAGAAAACGaTGATCGAAACCGGTCATCCGAAACGCGGCGATATCaTGatgTTTAAATATCCGGAAGATCCAAAGCTTGATTACATCAAGCGCGCGGTGGGTTTACCGGGCGATAAAGTCACTTACGATCCGGTCTCAAAAGAGaTGACGATgCAACCGGGATGCAGTTCCGGCCAGGCGTGTGAAAACGCGCTGCCGGTCACCTACTCAAACGTGGAACCGAGCGATTTCGTTCAGACCTTCTCACGCCGTAATGGTGGGGAAGCGACCAGCGGATTCTTTGAAGTGCCGAAACAGGAAACCAAAGAAAATGGAATTCGTCTTTCCGAGactagtGGTGGTCCAGGT**GCTCTGGCAGCTCTGGCACTGGCTGCATTAGCCGCTCTGGCCCTGGCCGCCTTAGCC**GGTCCTGGCGGTacggtaccAGGTCAACAAaatgcaGTTTGGATTGTTCCTCCTGGACAATACTTCATGATGGGCGACAACCGCGACAACAGCGCGGACAGCCG**GTTCTTCT**GGCTTTGTGCCGGAAGCGAATCTGGTCGGTCGGGCAACGGCTATCTGGATGAGCTTCGAaAAGCAAGAAGGCGAATGGCCGACTGGTCTGCGCTTAAGTCGCATTGGCGGCATCCA**TGa**tGATCCATCTTCGTTCACGTTGTCGCCGTTATGGCGACCCGGATCCCCGGGCGAGCTCGAATTCCGGTCTCCCTATAGTGAGTCGTATTccTTTCGATccGCCAGCTGCATTcAgGAATCGGCCAACGCGCGGGGAGAGGCGGTTTGCGTATTGGGCGCTCTTCCGCTTCCTCGCTCACTGACTCGCTGCGCTCGGTCGTTCGGCTGCGGCGAGCGGTATCAGCTCACTCAAAGGCGGTAA

**13. LepB *in vitro* Translation Reporter (N_out_, LALA TMD, L=45) (0-Frame = 44.5 kDa, PRF ~ 35.9 kDa)**

ATGGCGAATATGTTTGCCCTGATTCTGGTGATTGCCACACTGGTGACGGGCATTTTATGGTGCGTGGATAAATTCTTTTTCGCACCTAAACGGCGGGAACGTCAGGCAGCGGCGCAGGCGGCTGCCGGGGACTCACTGGATAAAGCAACGTTGAAAAAGGTTGCGCCGAAGCCTGGCTGGCTGGAAACCGGTGCTTCTGTTaTgCCGGTACTGatgATCGTATTGATTGTGCGTTCGTTTATTTATGAACCGTTCCAGATCCCGTCAGGTTCGATGATGCCGACTCTGaactctactGATTTTATTCTGGTAGAGAAGTTTGCTTATGGCATgAAAGATCCTATgTACCAGAAAACGaTGATCGAAACCGGTCATCCGAAACGCGGCGATATCaTGaTgTTTAAATATCCGGAAGATCCAAAGCTTGATTACATCAAGCGCGCGGTGGGTTTACCGGGCGATAAAGTCACTTACGATCCGGTCTCAAAAGAGaTGACGATgCAACCGGGATGCAGTTCCGGCCAGGCGTGTGAAAACGCGCTGCCGGTCACCTACTCAAACGTGGAACCGAGCGATTTCGTTCAGACCTTCTCACGCCGTAATGGTGGGGAAGCGACCAGCGGATTCTTTGAAGTGCCGAAACAGGAAACCAAAGAAAATGGAATTCGTCTTTCCGAGactagtGGTGGTCCAGGT**GCTCTGGCAGCTCTGGCACTGGCTGCATTAGCCGCTCTGGCCCTGGCCGCCTTAGCC**GGTCCTGGCGGTacggtaccAGGTCAACAAaatgcaGTTTGGATTGTTCCTCCTGGACAATACTTCATGATGGGCGACAACCGCGACAACAGCGCGGACAGCCGTTACTGGGGCTTTGTGCCGGAAGCGAATCT**TTTTTTT**CGGGCAACGGCTATCTGGATGAGCTTCGAaAAGCAAGAAGGCGAATGGCCGACTGGTCTGCGCTTAAGTCGCATTGGCGGCATCCA**TGa**tGATCCATCTTCGTTCACGTTGTCGCCGTTATGGCGACCCGGATCCCCGGGCGAGCTCGAATTCCGGTCTCCCTATAGTGAGTCGTATTccTTTCGATccGCCAGCTGCATTcAgGAATCGGCCAACGCGCGGGGAGAGGCGGTTTGCGTATTGGGCGCTCTTCCGCTTCCTCGCTCACTGACTCGCTGCGCTCGGTCGTTCGGCTGCGGCGAGCGGTATCAGCTCACTCAAAGGCGG**TAA**

**14. Full-Length Marker, LepB *in vitro* Translation Reporter (N_out_, LALA TMD, L=45) (0-Frame = 44.5 kDa)**

ATGGCGAATATGTTTGCCCTGATTCTGGTGATTGCCACACTGGTGACGGGCATTTTATGGTGCGTGGATAAATTCTTTTTCGCACCTAAACGGCGGGAACGTCAGGCAGCGGCGCAGGCGGCTGCCGGGGACTCACTGGATAAAGCAACGTTGAAAAAGGTTGCGCCGAAGCCTGGCTGGCTGGAAACCGGTGCTTCTGTTaTgCCGGTACTGatgATCGTATTGATTGTGCGTTCGTTTATTTATGAACCGTTCCAGATCCCGTCAGGTTCGATGATGCCGACTCTGaactctactGATTTTATTCTGGTAGAGAAGTTTGCTTATGGCATgAAAGATCCTATgTACCAGAAAACGaTGATCGAAACCGGTCATCCGAAACGCGGCGATATCaTGaTgTTTAAATATCCGGAAGATCCAAAGCTTGATTACATCAAGCGCGCGGTGGGTTTACCGGGCGATAAAGTCACTTACGATCCGGTCTCAAAAGAGaTGACGATgCAACCGGGATGCAGTTCCGGCCAGGCGTGTGAAAACGCGCTGCCGGTCACCTACTCAAACGTGGAACCGAGCGATTTCGTTCAGACCTTCTCACGCCGTAATGGTGGGGAAGCGACCAGCGGATTCTTTGAAGTGCCGAAACAGGAAACCAAAGAAAATGGAATTCGTCTTTCCGAGactagtGGTGGTCCAGGT**GCTCTGGCAGCTCTGGCACTGGCTGCATTAGCCGCTCTGGCCCTGGCCGCCTTAGCC**GGTCCTGGCGGTacggtaccAGGTCAACAAaatgcaGTTTGGATTGTTCCTCCTGGACAATACTTCATGATGGGCGACAACCGCGACAACAGCGCGGACAGCCGTTACTGGGGCTTTGTGCCGGAAGCGAATCT**GTTCTTC**CGGGCAACGGCTATCTGGATGAGCTTCGAaAAGCAAGAAGGCGAATGGCCGACTGGTCTGCGCTTAAGTCGCATTGGCGGCATCCATGatGATCCATCTTCGTTCACGTTGTCGCCGTTATGGCGACCCGGATCCCCGGGCGAGCTCGAATTCCGGTCTCCCTATAGTGAGTCGTATTccTTTCGATccGCCAGCTGCATTcAgGAATCGGCCAACGCGCGGGGAGAGGCGGTTTGCGTATTGGGCGCTCTTCCGCTTCCTCGCTCACTGACTCGCTGCGCTCGGTCGTTCGGCTGCGGCGAGCGGTATCAGCTCACTCAAAGGCGG**TAA**

**15. PRF Marker, LepB *in vitro* Translation Reporter (N_out_, LALA TMD, L=45) (0-Frame = 35.8 kDa)**

ATGGCGAATATGTTTGCCCTGATTCTGGTGATTGCCACACTGGTGACGGGCATTTTATGGTGCGTGGATAAATTCTTTTTCGCACCTAAACGGCGGGAACGTCAGGCAGCGGCGCAGGCGGCTGCCGGGGACTCACTGGATAAAGCAACGTTGAAAAAGGTTGCGCCGAAGCCTGGCTGGCTGGAAACCGGTGCTTCTGTTaTgCCGGTACTGatgATCGTATTGATTGTGCGTTCGTTTATTTATGAACCGTTCCAGATCCCGTCAGGTTCGATGATGCCGACTCTGaactctactGATTTTATTCTGGTAGAGAAGTTTGCTTATGGCATgAAAGATCCTATgTACCAGAAAACGaTGATCGAAACCGGTCATCCGAAACGCGGCGATATCaTGaTgTTTAAATATCCGGAAGATCCAAAGCTTGATTACATCAAGCGCGCGGTGGGTTTACCGGGCGATAAAGTCACTTACGATCCGGTCTCAAAAGAGaTGACGATgCAACCGGGATGCAGTTCCGGCCAGGCGTGTGAAAACGCGCTGCCGGTCACCTACTCAAACGTGGAACCGAGCGATTTCGTTCAGACCTTCTCACGCCGTAATGGTGGGGAAGCGACCAGCGGATTCTTTGAAGTGCCGAAACAGGAAACCAAAGAAAATGGAATTCGTCTTTCCGAGactagtGGTGGTCCAGGT**GCTCTGGCAGCTCTGGCACTGGCTGCATTAGCCGCTCTGGCCCTGGCCGCCTTAGCC**GGTCCTGGCGGTacggtaccAGGTCAACAAaatgcaGTTTGGATTGTTCCTCCTGGACAATACTTCATGATGGGCGACAACCGCGACAACAGCGCGGACAGCCGTTACTGGGGCTTTGTGCCGGAAGCGAATCT**GTTCTTCT**CGGGCAACGGCTATCTGGATGAGCTTCGAaAAGCAAGAAGGCGAATGGCCGACTGGTCTGCGCTTAAGTCGCATTGGCGGCATCCA**TGa**tGATCCATCTTCGTTCACGTTGTCGCCGTTATGGCGACCCGGATCCCCGGGCGAGCTCGAATTCCGGTCTCCCTATAGTGAGTCGTATTccTTTCGATccGCCAGCTGCATTcAgGAATCGGCCAACGCGCGGGGAGAGGCGGTTTGCGTATTGGGCGCTCTTCCGCTTCCTCGCTCACTGACTCGCTGCGCTCGGTCGTTCGGCTGCGGCGAGCGGTATCAGCTCACTCAAAGGCGGTAA

**16. LepB *in vitro* Translation Reporter (N_out_, LALA TMD, L=55) (0-Frame = 44.4 kDa, PRF ~ 35.8 kDa)**

ATGGCGAATATGTTTGCCCTGATTCTGGTGATTGCCACACTGGTGACGGGCATTTTATGGTGCGTGGATAAATTCTTTTTCGCACCTAAACGGCGGGAACGTCAGGCAGCGGCGCAGGCGGCTGCCGGGGACTCACTGGATAAAGCAACGTTGAAAAAGGTTGCGCCGAAGCCTGGCTGGCTGGAAACCGGTGCTTCTGTTatgCCGGTACTGatgATCGTATTGATTGTGCGTTCGTTTATTTATGAACCGTTCCAGATCCCGTCAGGTTCGATGATGCCGACTCTGaactctactGATTTTATTCTGGTAGAGAAGTTTGCTTATGGCATgAAAGATCCTATgTACCAGAAAACGaTGATCGAAACCGGTCATCCGAAACGCGGCGATATCaTGatgTTTAAATATCCGGAAGATCCAAAGCTTGATTACATCAAGCGCGCGGTGGGTTTACCGGGCGATAAAGTCACTTACGATCCGGTCTCAAAAGAGaTGACGATgCAACCGGGATGCAGTTCCGGCCAGGCGTGTGAAAACGCGCTGCCGGTCACCTACTCAAACGTGGAACCGAGCGATTTCGTTCAGACCTTCTCACGCCGTAATGGTGGGGAAGCGACCAGCGGATTCTTTGAAGTGCCGAAACAGGAAACCAAAGAAAATGGAATTCGTCTTTCCGAGactagtGGTGGTCCAGGT**GCTCTGGCAGCTCTGGCACTGGCTGCATTAGCCGCTCTGGCCCTGGCCGCCTTAGCC**GGTCCTGGCGGTacggtaccAGGTCAACAAaatgcaGTTTGGATTGTTCCTCCTGGACAATACTTCATGATGGGCGACAACCGCGACAACAGCGCGGACAGCCGTTACTGGGGCTTTGTGCCGGAAGCGAATCTGGTCGGTCGGGCAACGGCTATCTGGATGAG**TTTTTTT**AcGCAAGAAGGCGAATGGCCGACTGGTCTGCGCTTAAGTCGCATTGGCGGCATCCA**TGa**tGATCCATCTTCGTTCACGTTGTCGCCGTTATGGCGACCCGGATCCCCGGGCGAGCTCGAATTCCGGTCTCCCTATAGTGAGTCGTATTccTTTCGATccGCCAGCTGCATTcAgGAATCGGCCAACGCGCGGGGAGAGGCGGTTTGCGTATTGGGCGCTCTTCCGCTTCCTCGCTCACTGACTCGCTGCGCTCGGTCGTTCGGCTGCGGCGAGCGGTATCAGCTCACTCAAAGGCGG**TAA**

**17. Full-Length Marker, LepB *in vitro* Translation Reporter (N_out_, LALA TMD, L=55) (0-Frame = 44.4 kDa)**

ATGGCGAATATGTTTGCCCTGATTCTGGTGATTGCCACACTGGTGACGGGCATTTTATGGTGCGTGGATAAATTCTTTTTCGCACCTAAACGGCGGGAACGTCAGGCAGCGGCGCAGGCGGCTGCCGGGGACTCACTGGATAAAGCAACGTTGAAAAAGGTTGCGCCGAAGCCTGGCTGGCTGGAAACCGGTGCTTCTGTTatgCCGGTACTGatgATCGTATTGATTGTGCGTTCGTTTATTTATGAACCGTTCCAGATCCCGTCAGGTTCGATGATGCCGACTCTGaactctactGATTTTATTCTGGTAGAGAAGTTTGCTTATGGCATgAAAGATCCTATgTACCAGAAAACGaTGATCGAAACCGGTCATCCGAAACGCGGCGATATCaTGatgTTTAAATATCCGGAAGATCCAAAGCTTGATTACATCAAGCGCGCGGTGGGTTTACCGGGCGATAAAGTCACTTACGATCCGGTCTCAAAAGAGaTGACGATgCAACCGGGATGCAGTTCCGGCCAGGCGTGTGAAAACGCGCTGCCGGTCACCTACTCAAACGTGGAACCGAGCGATTTCGTTCAGACCTTCTCACGCCGTAATGGTGGGGAAGCGACCAGCGGATTCTTTGAAGTGCCGAAACAGGAAACCAAAGAAAATGGAATTCGTCTTTCCGAGactagtGGTGGTCCAGGT**GCTCTGGCAGCTCTGGCACTGGCTGCATTAGCCGCTCTGGCCCTGGCCGCCTTAGCC**GGTCCTGGCGGTacggtaccAGGTCAACAAaatgcaGTTTGGATTGTTCCTCCTGGACAATACTTCATGATGGGCGACAACCGCGACAACAGCGCGGACAGCCGTTACTGGGGCTTTGTGCCGGAAGCGAATCTGGTCGGTCGGGCAACGGCTATCTGGATG**TCGTTCTTC**AcGCAAGAAGGCGAATGGCCGACTGGTCTGCGCTTAAGTCGCATTGGCGGCATCCATGatGATCCATCTTCGTTCACGTTGTCGCCGTTATGGCGACCCGGATCCCCGGGCGAGCTCGAATTCCGGTCTCCCTATAGTGAGTCGTATTccTTTCGATccGCCAGCTGCATTcAgGAATCGGCCAACGCGCGGGGAGAGGCGGTTTGCGTATTGGGCGCTCTTCCGCTTCCTCGCTCACTGACTCGCTGCGCTCGGTCGTTCGGCTGCGGCGAGCGGTATCAGCTCACTCAAAGGCGG**TAA**

**18. PRF Marker, LepB *in vitro* Translation Reporter (N_out_, LALA TMD, L=55) (0-Frame = 35.8 kDa)**

ATGGCGAATATGTTTGCCCTGATTCTGGTGATTGCCACACTGGTGACGGGCATTTTATGGTGCGTGGATAAATTCTTTTTCGCACCTAAACGGCGGGAACGTCAGGCAGCGGCGCAGGCGGCTGCCGGGGACTCACTGGATAAAGCAACGTTGAAAAAGGTTGCGCCGAAGCCTGGCTGGCTGGAAACCGGTGCTTCTGTTatgCCGGTACTGatgATCGTATTGATTGTGCGTTCGTTTATTTATGAACCGTTCCAGATCCCGTCAGGTTCGATGATGCCGACTCTGaactctactGATTTTATTCTGGTAGAGAAGTTTGCTTATGGCATgAAAGATCCTATgTACCAGAAAACGaTGATCGAAACCGGTCATCCGAAACGCGGCGATATCaTGatgTTTAAATATCCGGAAGATCCAAAGCTTGATTACATCAAGCGCGCGGTGGGTTTACCGGGCGATAAAGTCACTTACGATCCGGTCTCAAAAGAGaTGACGATgCAACCGGGATGCAGTTCCGGCCAGGCGTGTGAAAACGCGCTGCCGGTCACCTACTCAAACGTGGAACCGAGCGATTTCGTTCAGACCTTCTCACGCCGTAATGGTGGGGAAGCGACCAGCGGATTCTTTGAAGTGCCGAAACAGGAAACCAAAGAAAATGGAATTCGTCTTTCCGAGactagtGGTGGTCCAGGT**GCTCTGGCAGCTCTGGCACTGGCTGCATTAGCCGCTCTGGCCCTGGCCGCCTTAGCC**GGTCCTGGCGGTacggtaccAGGTCAACAAaatgcaGTTTGGATTGTTCCTCCTGGACAATACTTCATGATGGGCGACAACCGCGACAACAGCGCGGACAGCCGTTACTGGGGCTTTGTGCCGGAAGCGAATCTGGTCGGTCGGGCAACGGCTATCTGGATG**TCGTTCTTCT**AcGCAAGAAGGCGAATGGCCGACTGGTCTGCGCTTAAGTCGCATTGGCGGCATCCA**TGa**tGATCCATCTTCGTTCACGTTGTCGCCGTTATGGCGACCCGGATCCCCGGGCGAGCTCGAATTCCGGTCTCCCTATAGTGAGTCGTATTccTTTCGATccGCCAGCTGCATTcAgGAATCGGCCAACGCGCGGGGAGAGGCGGTTTGCGTATTGGGCGCTCTTCCGCTTCCTCGCTCACTGACTCGCTGCGCTCGGTCGTTCGGCTGCGGCGAGCGGTATCAGCTCACTCAAAGGCGGTAA

**19. LepB *in vitro* Translation Reporter (N_in_, AVAA TMD, L=45) (0-Frame = 45.6 kDa, PRF ~ 36.9 kDa)**

ATGGCGAATATGTTTGCCCTGATTCTGGTGATTGCCACACTGGTGACGGGCATTTTATGGTGCGTGGATAAATTCTTTTTCGCACCTAAACGGCGGGAACGTCAGGCAGCGGCGCAGGCGGCTGCCGGGGACTCACTGGATAAAGCAACGTTGAAAAAGGTTGCGCCGAAGCCTGGCTGGCTGGAAACCGGTGCTTCTGTTatgCCGGTACTGatgATCGTATTGATTGTGCGTTCGTTTATTTATGAACCGTTCCAGATCCCGTCAGGTTCGATGATGCCGACTCTGaactctactGATTTTATTCTGGTAGAGAAGTTTGCTTATGGCATgAAAGATCCTATgTACCAGAAAACGaTGATCGAAACCGGTCATCCGAAACGCGGCGATATCaTGaTgTTTAAATATCCGGAAGATCCAAAGCTTGATTACATCAAGCGCGCGGTGGGTTTACCGGGCGATAAAGTCACTTACGATCCGGTCTCAAAAGAGaTGACGATgCAACCGGGATGCAGTTCCGGCCAGGCGTGTGAAAACGCGCTGCCG**ggtggtccaggcgcaTTAgcagcactggcactggcggcgctggctgcactggcaTTAgcggcactggcgggtccaggcggc**AGCGATTTCGTTCAGACCTTCTCACGCCGTAATGGTGGGGAAGCGACCAGCGGATTCTTTGAAGTGCCGAAACAGGAAACCAAAGAAAATGGAATTCGTCTTTCCGAGactagtggTggTccaggT**GCAGCAGCTgcTGTTGCAGCTgcTGCTGCTGCAgcAGCAGCCGTTgcTGCTGCAGC**TggTcctggcggTacggtaccAGGTCAACAAaatgcaGTTTGGATTGTTCCTCCTGGACAATACTTCATGATGGGCGACAACCGCGACAACAGCGCGGACAGCCGTTACTGGGGCTTTGTGCCGGAAGCGAATCT**TTTTTTT**CGGGCAACGGCTATCTGGATGAGCTTCGAaAAGCAAGAAGGCGAATGGCCGACTGGTCTGCGCTTAAGTCGCATTGGCGGCATCCA**TGa**tGATCCATCTTCGTTCACGTTGTCGCCGTTATGGCGACCCGGATCCCCGGGCGAGCTCGAATTCCGGTCTCCCTATAGTGAGTCGTATTccTTTCGATccGCCAGCTGCATTcAgGAATCGGCCAACGCGCGGGGAGAGGCGGTTTGCGTATTGGGCGCTCTTCCGCTTCCTCGCTCACTGACTCGCTGCGCTCGGTCGTTCGGCTGCGGCGAGCGGTATCAGCTCACTCAAAGGCGG**TAA**

**20. Full-Length Marker, LepB *in vitro* Translation Reporter (N_in_, AVAA TMD, L=45) (0-Frame = 45.6 kDa)**

ATGGCGAATATGTTTGCCCTGATTCTGGTGATTGCCACACTGGTGACGGGCATTTTATGGTGCGTGGATAAATTCTTTTTCGCACCTAAACGGCGGGAACGTCAGGCAGCGGCGCAGGCGGCTGCCGGGGACTCACTGGATAAAGCAACGTTGAAAAAGGTTGCGCCGAAGCCTGGCTGGCTGGAAACCGGTGCTTCTGTTaTgCCGGTACTGatgATCGTATTGATTGTGCGTTCGTTTATTTATGAACCGTTCCAGATCCCGTCAGGTTCGATGATGCCGACTCTGaactctactGATTTTATTCTGGTAGAGAAGTTTGCTTATGGCATgAAAGATCCTATgTACCAGAAAACGaTGATCGAAACCGGTCATCCGAAACGCGGCGATATCaTGaTgTTTAAATATCCGGAAGATCCAAAGCTTGATTACATCAAGCGCGCGGTGGGTTTACCGGGCGATAAAGTCACTTACGATCCGGTCTCAAAAGAGaTGACGATgCAACCGGGATGCAGTTCCGGCCAGGCGTGTGAAAACGCGCTGCCG**ggtggtccaggcgcaTTAgcagcactggcactggcggcgctggctgcactggcaTTAgcggcactggcgggtccaggcggc**AGCGATTTCGTTCAGACCTTCTCACGCCGTAATGGTGGGGAAGCGACCAGCGGATTCTTTGAAGTGCCGAAACAGGAAACCAAAGAAAATGGAATTCGTCTTTCCGAGactagtggTggTccaggT**GCAGCAGCTgcTGTTGCAGCTgcTGCTGCTGCAgcAGCAGCCGTTgcTGCTGCAGC**TggTcctggcggTacggtaccAGGTCAACAAaatgcaGTTTGGATTGTTCCTCCTGGACAATACTTCATGATGGGCGACAACCGCGACAACAGCGCGGACAGCCGTTACTGGGGCTTTGTGCCGGAAGCGAATCT**GTTCTTC**CGGGCAACGGCTATCTGGATGAGCTTCGAaAAGCAAGAAGGCGAATGGCCGACTGGTCTGCGCTTAAGTCGCATTGGCGGCATCCATGatGATCCATCTTCGTTCACGTTGTCGCCGTTATGGCGACCCGGATCCCCGGGCGAGCTCGAATTCCGGTCTCCCTATAGTGAGTCGTATTccTTTCGATccGCCAGCTGCATTcAgGAATCGGCCAACGCGCGGGGAGAGGCGGTTTGCGTATTGGGCGCTCTTCCGCTTCCTCGCTCACTGACTCGCTGCGCTCGGTCGTTCGGCTGCGGCGAGCGGTATCAGCTCACTCAAAGGCGG**TAA**

**21. PRF Marker, LepB *in vitro* Translation Reporter (N_in_, AVAA TMD, L=45) (0-Frame = 36.9 kDa)**

ATGGCGAATATGTTTGCCCTGATTCTGGTGATTGCCACACTGGTGACGGGCATTTTATGGTGCGTGGATAAATTCTTTTTCGCACCTAAACGGCGGGAACGTCAGGCAGCGGCGCAGGCGGCTGCCGGGGACTCACTGGATAAAGCAACGTTGAAAAAGGTTGCGCCGAAGCCTGGCTGGCTGGAAACCGGTGCTTCTGTTaTgCCGGTACTGatgATCGTATTGATTGTGCGTTCGTTTATTTATGAACCGTTCCAGATCCCGTCAGGTTCGATGATGCCGACTCTGaactctactGATTTTATTCTGGTAGAGAAGTTTGCTTATGGCATgAAAGATCCTATgTACCAGAAAACGaTGATCGAAACCGGTCATCCGAAACGCGGCGATATCaTGaTgTTTAAATATCCGGAAGATCCAAAGCTTGATTACATCAAGCGCGCGGTGGGTTTACCGGGCGATAAAGTCACTTACGATCCGGTCTCAAAAGAGaTGACGATgCAACCGGGATGCAGTTCCGGCCAGGCGTGTGAAAACGCGCTGCCG**ggtggtccaggcgcaTTAgcagcactggcactggcggcgctggctgcactggcaTTAgcggcactggcgggtccaggcggc**AGCGATTTCGTTCAGACCTTCTCACGCCGTAATGGTGGGGAAGCGACCAGCGGATTCTTTGAAGTGCCGAAACAGGAAACCAAAGAAAATGGAATTCGTCTTTCCGAGactagtggTggTccaggT**GCAGCAGCTgcTGTTGCAGCTgcTGCTGCTGCAgcAGCAGCCGTTgcTGCTGCAGC**TggTcctggcggTacggtaccAGGTCAACAAaatgcaGTTTGGATTGTTCCTCCTGGACAATACTTCATGATGGGCGACAACCGCGACAACAGCGCGGACAGCCGTTACTGGGGCTTTGTGCCGGAAGCGAATCT**GTTCTTCT**CGGGCAACGGCTATCTGGATGAGCTTCGAaAAGCAAGAAGGCGAATGGCCGACTGGTCTGCGCTTAAGTCGCATTGGCGGCATCCA**TGa**tGATCCATCTTCGTTCACGTTGTCGCCGTTATGGCGACCCGGATCCCCGGGCGAGCTCGAATTCCGGTCTCCCTATAGTGAGTCGTATTccTTTCGATccGCCAGCTGCATTcAgGAATCGGCCAACGCGCGGGGAGAGGCGGTTTGCGTATTGGGCGCTCTTCCGCTTCCTCGCTCACTGACTCGCTGCGCTCGGTCGTTCGGCTGCGGCGAGCGGTATCAGCTCACTCAAAGGCGGTAA

**22. LepB *in vitro* Translation Reporter (N_in_, LALA TMD, L=45) (0-Frame = kDa, PRF ~ kDa)**

ATGGCGAATATGTTTGCCCTGATTCTGGTGATTGCCACACTGGTGACGGGCATTTTATGGTGCGTGGATAAATTCTTTTTCGCACCTAAACGGCGGGAACGTCAGGCAGCGGCGCAGGCGGCTGCCGGGGACTCACTGGATAAAGCAACGTTGAAAAAGGTTGCGCCGAAGCCTGGCTGGCTGGAAACCGGTGCTTCTGTTaTgCCGGTACTGatgATCGTATTGATTGTGCGTTCGTTTATTTATGAACCGTTCCAGATCCCGTCAGGTTCGATGATGCCGACTCTGaactctactGATTTTATTCTGGTAGAGAAGTTTGCTTATGGCATgAAAGATCCTATgTACCAGAAAACGaTGATCGAAACCGGTCATCCGAAACGCGGCGATATCaTGaTgTTTAAATATCCGGAAGATCCAAAGCTTGATTACATCAAGCGCGCGGTGGGTTTACCGGGCGATAAAGTCACTTACGATCCGGTCTCAAAAGAGaTGACGATgCAACCGGGATGCAGTTCCGGCCAGGCGTGTGAAAACGCGCTGCCG**ggtggtccaggcgcaTTAgcagcactggcactggcggcgctggctgcactggcaTTAgcggcactggcgggtccaggcggc**AGCGATTTCGTTCAGACCTTCTCACGCCGTAATGGTGGGGAAGCGACCAGCGGATTCTTTGAAGTGCCGAAACAGGAAACCAAAGAAAATGGAATTCGTCTTTCCGAGactagtGGTGGTCCAGGT**GCTCTGGCAGCTCTGGCACTGGCTGCATTAGCCGCTCTGGCCCTGGCCGCCTTAGCC**GGTCCTGGCGGTacggtaccAGGTCAACAAaatgcaGTTTGGATTGTTCCTCCTGGACAATACTTCATGATGGGCGACAACCGCGACAACAGCGCGGACAGCCGTTACTGGGGCTTTGTGCCGGAAGCGAATCT**TTTTTTT**CGGGCAACGGCTATCTGGATGAGCTTCGAaAAGCAAGAAGGCGAATGGCCGACTGGTCTGCGCTTAAGTCGCATTGGCGGCATCCA**TGa**tGATCCATCTTCGTTCACGTTGTCGCCGTTATGGCGACCCGGATCCCCGGGCGAGCTCGAATTCCGGTCTCCCTATAGTGAGTCGTATTccTTTCGATccGCCAGCTGCATTcAgGAATCGGCCAACGCGCGGGGAGAGGCGGTTTGCGTATTGGGCGCTCTTCCGCTTCCTCGCTCACTGACTCGCTGCGCTCGGTCGTTCGGCTGCGGCGAGCGGTATCAGCTCACTCAAAGGCGG**TAA**

**23. Full-Length Marker, LepB in vitro Translation Reporter (N_in_, LALA TMD, L=45) (0-Frame = kDa)**

ATGGCGAATATGTTTGCCCTGATTCTGGTGATTGCCACACTGGTGACGGGCATTTTATGGTGCGTGGATAAATTCTTTTTCGCACCTAAACGGCGGGAACGTCAGGCAGCGGCGCAGGCGGCTGCCGGGGACTCACTGGATAAAGCAACGTTGAAAAAGGTTGCGCCGAAGCCTGGCTGGCTGGAAACCGGTGCTTCTGTTaTgCCGGTACTGatgATCGTATTGATTGTGCGTTCGTTTATTTATGAACCGTTCCAGATCCCGTCAGGTTCGATGATGCCGACTCTGaactctactGATTTTATTCTGGTAGAGAAGTTTGCTTATGGCATgAAAGATCCTATgTACCAGAAAACGaTGATCGAAACCGGTCATCCGAAACGCGGCGATATCaTGaTgTTTAAATATCCGGAAGATCCAAAGCTTGATTACATCAAGCGCGCGGTGGGTTTACCGGGCGATAAAGTCACTTACGATCCGGTCTCAAAAGAGaTGACGATgCAACCGGGATGCAGTTCCGGCCAGGCGTGTGAAAACGCGCTGCCG**ggtggtccaggcgcaTTAgcagcactggcactggcggcgctggctgcactggcaTTAgcggcactggcgggtccaggcggc**AGCGATTTCGTTCAGACCTTCTCACGCCGTAATGGTGGGGAAGCGACCAGCGGATTCTTTGAAGTGCCGAAACAGGAAACCAAAGAAAATGGAATTCGTCTTTCCGAGactagtGGTGGTCCAGGT**GCTCTGGCAGCTCTGGCACTGGCTGCATTAGCCGCTCTGGCCCTGGCCGCCTTAGCC**GGTCCTGGCGGTacggtaccAGGTCAACAAaatgcaGTTTGGATTGTTCCTCCTGGACAATACTTCATGATGGGCGACAACCGCGACAACAGCGCGGACAGCCGTTACTGGGGCTTTGTGCCGGAAGCGAATCT**GTTCTTC**CGGGCAACGGCTATCTGGATGAGCTTCGAaAAGCAAGAAGGCGAATGGCCGACTGGTCTGCGCTTAAGTCGCATTGGCGGCATCCATGatGATCCATCTTCGTTCACGTTGTCGCCGTTATGGCGACCCGGATCCCCGGGCGAGCTCGAATTCCGGTCTCCCTATAGTGAGTCGTATTccTTTCGATccGCCAGCTGCATTcAgGAATCGGCCAACGCGCGGGGAGAGGCGGTTTGCGTATTGGGCGCTCTTCCGCTTCCTCGCTCACTGACTCGCTGCGCTCGGTCGTTCGGCTGCGGCGAGCGGTATCAGCTCACTCAAAGGCGG**TAA**

**24. PRF Marker, LepB in vitro Translation Reporter (N_in_, LALA TMD, L=45) (0-Frame = kDa)**

ATGGCGAATATGTTTGCCCTGATTCTGGTGATTGCCACACTGGTGACGGGCATTTTATGGTGCGTGGATAAATTCTTTTTCGCACCTAAACGGCGGGAACGTCAGGCAGCGGCGCAGGCGGCTGCCGGGGACTCACTGGATAAAGCAACGTTGAAAAAGGTTGCGCCGAAGCCTGGCTGGCTGGAAACCGGTGCTTCTGTTaTgCCGGTACTGatgATCGTATTGATTGTGCGTTCGTTTATTTATGAACCGTTCCAGATCCCGTCAGGTTCGATGATGCCGACTCTGaactctactGATTTTATTCTGGTAGAGAAGTTTGCTTATGGCATgAAAGATCCTATgTACCAGAAAACGaTGATCGAAACCGGTCATCCGAAACGCGGCGATATCaTGaTgTTTAAATATCCGGAAGATCCAAAGCTTGATTACATCAAGCGCGCGGTGGGTTTACCGGGCGATAAAGTCACTTACGATCCGGTCTCAAAAGAGaTGACGATgCAACCGGGATGCAGTTCCGGCCAGGCGTGTGAAAACGCGCTGCCG**ggtggtccaggcgcaTTAgcagcactggcactggcggcgctggctgcactggcaTTAgcggcactggcgggtccaggcggc**AGCGATTTCGTTCAGACCTTCTCACGCCGTAATGGTGGGGAAGCGACCAGCGGATTCTTTGAAGTGCCGAAACAGGAAACCAAAGAAAATGGAATTCGTCTTTCCGAGactagtGGTGGTCCAGGT**GCTCTGGCAGCTCTGGCACTGGCTGCATTAGCCGCTCTGGCCCTGGCCGCCTTAGCC**GGTCCTGGCGGTacggtaccAGGTCAACAAaatgcaGTTTGGATTGTTCCTCCTGGACAATACTTCATGATGGGCGACAACCGCGACAACAGCGCGGACAGCCGTTACTGGGGCTTTGTGCCGGAAGCGAATCT**GTTCTTCT**CGGGCAACGGCTATCTGGATGAGCTTCGAaAAGCAAGAAGGCGAATGGCCGACTGGTCTGCGCTTAAGTCGCATTGGCGGCATCCA**TGa**tGATCCATCTTCGTTCACGTTGTCGCCGTTATGGCGACCCGGATCCCCGGGCGAGCTCGAATTCCGGTCTCCCTATAGTGAGTCGTATTccTTTCGATccGCCAGCTGCATTcAgGAATCGGCCAACGCGCGGGGAGAGGCGGTTTGCGTATTGGGCGCTCTTCCGCTTCCTCGCTCACTGACTCGCTGCGCTCGGTCGTTCGGCTGCGGCGAGCGGTATCAGCTCACTCAAAGGCGGTAA

**25. LepB Cellular Reporter (N_out_, AVAA TMD, L=45)**

ATGGCGAATATGTTTGCCCTGATTCTGGTGATTGCCACACTGGTGACGGGCATTTTATGGTGCGTGGATAAATTCTTTTTCGCACCTAAACGGCGGGAACGTCAGGCAGCGGCGCAGGCGGCTGCCGGGGACTCACTGGATAAAGCAACGTTGAAAAAGGTTGCGCCGAAGCCTGGCTGGCTGGAAACCGGTGCTTCTGTTATGCCGGTACTGATGATCGTATTGATTGTGCGTTCGTTTATTTATGAACCGTTCCAGATCCCGTCAGGTTCGATGATGCCGACTCTGAACTCTACTGATTTTATTCTGGTAGAGAAGTTTGCTTATGGCATGAAAGATCCTATGTACCAGAAAACGATGATCGAAACCGGTCATCCGAAACGCGGCGATATCATGATGTTTAAATATCCGGAAGATCCAAAGCTTGATTACATCAAGCGCGCGGTGGGTTTACCGGGCGATAAAGTCACTTACGATCCGGTCTCAAAAGAGATGACGATGCAACCGGGATGCAGTTCCGGCCAGGCGTGTGAAAACGCGCTGCCGGTCACCTACTCAAACGTGGAACCGAGCGATTTCGTTCAGACCTTCTCACGCCGTAATGGTGGGGAAGCGACCAGCGGATTCTTTGAAGTGCCGAAACAGGAAACCAAAGAAAATGGAATTCGTCTTTCCGAGACTAGTGGTGGTCCAGGT**GCAGCAGCTGCTGTTGCAGCTGCTGCTGCTGCAGCAGCAGCCGTTGCTGCTGCAGC**TGGTCCTGGCGGTACGGTACCAGGTCAACAAAATGCAGTTTGGATTGTTCCTCCTGGACAATACTTCATGATGGGCGACAACCGCGACAACAGCGCGGACAGCCGTTACTGGGGCTTTGTGCCGGAAGCGAATC**TTTTTTTT**CGGGCAACGGCTATCTGGATGAGCTTCGAAAAGCAAGAAGGCGAATGGCCGACTGGTCTGCGCTTAAGTCGCATTGGCGGCATCCATCAGGAGGAGGAGGATCT**GCTACTAATTTTTCACTTCTTAAGCAAGCCGGGGATGTCGAAGAAAATCCGGGACCAATGGCCACAACCATGACGGCCCTGACAGAAGGTGCGAAGCTGTTCGAGAAGGAGATTCCCTATATCACAGAATTGGAGGGGGATGTAGAGGGTATGAAGTTTATCATCAAAGGCGAAGGGACAGGGGATGCAACAACTGGAACAATTAAGGCTAAGTACATTTGCACGACCGGCGACGTCCCGGTGCCCTGGTCCACGCTCGTCACCACGCTCACGTACGGAGCCCAGTGCTTTGCCAAATATGGCCCTGAACTTAAAGACTTCTACAAGTCGTGTATGCCGGAGGGATACGTGCAAGAGAGGACGATCACCTTTGAAGGTGACGGAGTATTCAAAACAAGAGCGGAGGTGACGTTCGAGAATGGATCGGTCTATAACCGGGTCAAGCTCAACGGACAGGGCTTTAAGAAAGATGGACACGTCCTTGGGAAGAATTTGGAGTTCAATTTCACCCCGCATTGTCTTTACATCTGGGGTGATCAGGCGAATCACGGGTTGAAATCAGCGTTCAAGATCATGCACGAGATTACGGGGAGCAAAGAGGACTTTATCGTGGCAGACCACACTCAGATGAACACTCCAATCGGAGGGGGTCCCGTACACGTACCCGAGTATCATCACCTGACCGTCTGGACATCGTTTGGAAAAGACCCTGACGACGATGAAACTGATCATCTCAACATTGTGGAAGTGATCAAGGCGGTGGACTTGGAAACATACCGGTGA**

**26. SS_mut_ Control, LepB Cellular Reporter (N_out_, AVAA TMD, L=45)**

ATGGCGAATATGTTTGCCCTGATTCTGGTGATTGCCACACTGGTGACGGGCATTTTATGGTGCGTGGATAAATTCTTTTTCGCACCTAAACGGCGGGAACGTCAGGCAGCGGCGCAGGCGGCTGCCGGGGACTCACTGGATAAAGCAACGTTGAAAAAGGTTGCGCCGAAGCCTGGCTGGCTGGAAACCGGTGCTTCTGTTATGCCGGTACTGATGATCGTATTGATTGTGCGTTCGTTTATTTATGAACCGTTCCAGATCCCGTCAGGTTCGATGATGCCGACTCTGAACTCTACTGATTTTATTCTGGTAGAGAAGTTTGCTTATGGCATGAAAGATCCTATGTACCAGAAAACGATGATCGAAACCGGTCATCCGAAACGCGGCGATATCATGATGTTTAAATATCCGGAAGATCCAAAGCTTGATTACATCAAGCGCGCGGTGGGTTTACCGGGCGATAAAGTCACTTACGATCCGGTCTCAAAAGAGATGACGATGCAACCGGGATGCAGTTCCGGCCAGGCGTGTGAAAACGCGCTGCCGGTCACCTACTCAAACGTGGAACCGAGCGATTTCGTTCAGACCTTCTCACGCCGTAATGGTGGGGAAGCGACCAGCGGATTCTTTGAAGTGCCGAAACAGGAAACCAAAGAAAATGGAATTCGTCTTTCCGAGACTAGTGGTGGTCCAGGT**GCAGCAGCTGCTGTTGCAGCTGCTGCTGCTGCAGCAGCAGCCGTTGCTGCTGCAGC**TGGTCCTGGCGGTACGGTACCAGGTCAACAAAATGCAGTTTGGATTGTTCCTCCTGGACAATACTTCATGATGGGCGACAACCGCGACAACAGCGCGGACAGCCGTTACTGGGGCTTTGTGCCGGAAGCGAATCT**GTTCTTC**CGGGCAACGGCTATCTGGATGAGCTTCGAAAAGCAAGAAGGCGAATGGCCGACTGGTCTGCGCTTAAGTCGCATTGGCGGCATCCATCAGGAGGAGGAGGATCT**GCTACTAATTTTTCACTTCTTAAGCAAGCCGGGGATGTCGAAGAAAATCCGGGACCAATGGCCACAACCATGACGGCCCTGACAGAAGGTGCGAAGCTGTTCGAGAAGGAGATTCCCTATATCACAGAATTGGAGGGGGATGTAGAGGGTATGAAGTTTATCATCAAAGGCGAAGGGACAGGGGATGCAACAACTGGAACAATTAAGGCTAAGTACATTTGCACGACCGGCGACGTCCCGGTGCCCTGGTCCACGCTCGTCACCACGCTCACGTACGGAGCCCAGTGCTTTGCCAAATATGGCCCTGAACTTAAAGACTTCTACAAGTCGTGTATGCCGGAGGGATACGTGCAAGAGAGGACGATCACCTTTGAAGGTGACGGAGTATTCAAAACAAGAGCGGAGGTGACGTTCGAGAATGGATCGGTCTATAACCGGGTCAAGCTCAACGGACAGGGCTTTAAGAAAGATGGACACGTCCTTGGGAAGAATTTGGAGTTCAATTTCACCCCGCATTGTCTTTACATCTGGGGTGATCAGGCGAATCACGGGTTGAAATCAGCGTTCAAGATCATGCACGAGATTACGGGGAGCAAAGAGGACTTTATCGTGGCAGACCACACTCAGATGAACACTCCAATCGGAGGGGGTCCCGTACACGTACCCGAGTATCATCACCTGACCGTCTGGACATCGTTTGGAAAAGACCCTGACGACGATGAAACTGATCATCTCAACATTGTGGAAGTGATCAAGGCGGTGGACTTGGAAACATACCGGTGA**

**27. LepB Cellular Reporter (N_out_, LALA TMD, L=35)**

ATGGCGAATATGTTTGCCCTGATTCTGGTGATTGCCACACTGGTGACGGGCATTTTATGGTGCGTGGATAAATTCTTTTTCGCACCTAAACGGCGGGAACGTCAGGCAGCGGCGCAGGCGGCTGCCGGGGACTCACTGGATAAAGCAACGTTGAAAAAGGTTGCGCCGAAGCCTGGCTGGCTGGAAACCGGTGCTTCTGTTATGCCGGTACTGATGATCGTATTGATTGTGCGTTCGTTTATTTATGAACCGTTCCAGATCCCGTCAGGTTCGATGATGCCGACTCTGAACTCTACTGATTTTATTCTGGTAGAGAAGTTTGCTTATGGCATGAAAGATCCTATGTACCAGAAAACGATGATCGAAACCGGTCATCCGAAACGCGGCGATATCATGATGTTTAAATATCCGGAAGATCCAAAGCTTGATTACATCAAGCGCGCGGTGGGTTTACCGGGCGATAAAGTCACTTACGATCCGGTCTCAAAAGAGATGACGATGCAACCGGGATGCAGTTCCGGCCAGGCGTGTGAAAACGCGCTGCCGGTCACCTACTCAAACGTGGAACCGAGCGATTTCGTTCAGACCTTCTCACGCCGTAATGGTGGGGAAGCGACCAGCGGATTCTTTGAAGTGCCGAAACAGGAAACCAAAGAAAATGGAATTCGTCTTTCCGAGACTAGTGGTGGTCCAGGT**GCTCTGGCAGCTCTGGCACTGGCTGCATTAGCCGCTCTGGCCCTGGCCGCCTTAGCC**GGTCCTGGCGGTACGGTACCAGGTCAACAAAATGCAGTTTGGATTGTTCCTCCTGGACAATACTTCATGATGGGCGACAACCGCGACAACAGCGCGGACAGCCG**TTTTTTT**GGCTTTGTGCCGGAAGCGAATCTGGTCGGTCGGGCAACGGCTATCTGGATGAGCTTCGAAAAGCAAGAAGGCGAATGGCCGACTGGTCTGCGCTTAAGTCGCATTGGCGGCATCCATCAGGAGGAGGAGGATCT**GCTACTAATTTTTCACTTCTTAAGCAAGCCGGGGATGTCGAAGAAAATCCGGGACCAATGGCCACAACCATGACGGCCCTGACAGAAGGTGCGAAGCTGTTCGAGAAGGAGATTCCCTATATCACAGAATTGGAGGGGGATGTAGAGGGTATGAAGTTTATCATCAAAGGCGAAGGGACAGGGGATGCAACAACTGGAACAATTAAGGCTAAGTACATTTGCACGACCGGCGACGTCCCGGTGCCCTGGTCCACGCTCGTCACCACGCTCACGTACGGAGCCCAGTGCTTTGCCAAATATGGCCCTGAACTTAAAGACTTCTACAAGTCGTGTATGCCGGAGGGATACGTGCAAGAGAGGACGATCACCTTTGAAGGTGACGGAGTATTCAAAACAAGAGCGGAGGTGACGTTCGAGAATGGATCGGTCTATAACCGGGTCAAGCTCAACGGACAGGGCTTTAAGAAAGATGGACACGTCCTTGGGAAGAATTTGGAGTTCAATTTCACCCCGCATTGTCTTTACATCTGGGGTGATCAGGCGAATCACGGGTTGAAATCAGCGTTCAAGATCATGCACGAGATTACGGGGAGCAAAGAGGACTTTATCGTGGCAGACCACACTCAGATGAACACTCCAATCGGAGGGGGTCCCGTACACGTACCCGAGTATCATCACCTGACCGTCTGGACATCGTTTGGAAAAGACCCTGACGACGATGAAACTGATCATCTCAACATTGTGGAAGTGATCAAGGCGGTGGACTTGGAAACATACCGGTGA**

**28. SS_mut_ Control, LepB Cellular Reporter (N_out_, LALA TMD, L=35)**

ATGGCGAATATGTTTGCCCTGATTCTGGTGATTGCCACACTGGTGACGGGCATTTTATGGTGCGTGGATAAATTCTTTTTCGCACCTAAACGGCGGGAACGTCAGGCAGCGGCGCAGGCGGCTGCCGGGGACTCACTGGATAAAGCAACGTTGAAAAAGGTTGCGCCGAAGCCTGGCTGGCTGGAAACCGGTGCTTCTGTTATGCCGGTACTGATGATCGTATTGATTGTGCGTTCGTTTATTTATGAACCGTTCCAGATCCCGTCAGGTTCGATGATGCCGACTCTGAACTCTACTGATTTTATTCTGGTAGAGAAGTTTGCTTATGGCATGAAAGATCCTATGTACCAGAAAACGATGATCGAAACCGGTCATCCGAAACGCGGCGATATCATGATGTTTAAATATCCGGAAGATCCAAAGCTTGATTACATCAAGCGCGCGGTGGGTTTACCGGGCGATAAAGTCACTTACGATCCGGTCTCAAAAGAGATGACGATGCAACCGGGATGCAGTTCCGGCCAGGCGTGTGAAAACGCGCTGCCGGTCACCTACTCAAACGTGGAACCGAGCGATTTCGTTCAGACCTTCTCACGCCGTAATGGTGGGGAAGCGACCAGCGGATTCTTTGAAGTGCCGAAACAGGAAACCAAAGAAAATGGAATTCGTCTTTCCGAGACTAGTGGTGGTCCAGGT**GCTCTGGCAGCTCTGGCACTGGCTGCATTAGCCGCTCTGGCCCTGGCCGCCTTAGCC**GGTCCTGGCGGTACGGTACCAGGTCAACAAAATGCAGTTTGGATTGTTCCTCCTGGACAATACTTCATGATGGGCGACAACCGCGACAACAGCGCGGACAGCCG**GTTCTTC**GGCTTTGTGCCGGAAGCGAATCTGGTCGGTCGGGCAACGGCTATCTGGATGAGCTTCGAAAAGCAAGAAGGCGAATGGCCGACTGGTCTGCGCTTAAGTCGCATTGGCGGCATCCATCAGGAGGAGGAGGATCT**GCTACTAATTTTTCACTTCTTAAGCAAGCCGGGGATGTCGAAGAAAATCCGGGACCAATGGCCACAACCATGACGGCCCTGACAGAAGGTGCGAAGCTGTTCGAGAAGGAGATTCCCTATATCACAGAATTGGAGGGGGATGTAGAGGGTATGAAGTTTATCATCAAAGGCGAAGGGACAGGGGATGCAACAACTGGAACAATTAAGGCTAAGTACATTTGCACGACCGGCGACGTCCCGGTGCCCTGGTCCACGCTCGTCACCACGCTCACGTACGGAGCCCAGTGCTTTGCCAAATATGGCCCTGAACTTAAAGACTTCTACAAGTCGTGTATGCCGGAGGGATACGTGCAAGAGAGGACGATCACCTTTGAAGGTGACGGAGTATTCAAAACAAGAGCGGAGGTGACGTTCGAGAATGGATCGGTCTATAACCGGGTCAAGCTCAACGGACAGGGCTTTAAGAAAGATGGACACGTCCTTGGGAAGAATTTGGAGTTCAATTTCACCCCGCATTGTCTTTACATCTGGGGTGATCAGGCGAATCACGGGTTGAAATCAGCGTTCAAGATCATGCACGAGATTACGGGGAGCAAAGAGGACTTTATCGTGGCAGACCACACTCAGATGAACACTCCAATCGGAGGGGGTCCCGTACACGTACCCGAGTATCATCACCTGACCGTCTGGACATCGTTTGGAAAAGACCCTGACGACGATGAAACTGATCATCTCAACATTGTGGAAGTGATCAAGGCGGTGGACTTGGAAACATACCGGTGA**

**29. LepB Cellular Reporter (N_out_, LALA TMD, L=45)**

ATGGCGAATATGTTTGCCCTGATTCTGGTGATTGCCACACTGGTGACGGGCATTTTATGGTGCGTGGATAAATTCTTTTTCGCACCTAAACGGCGGGAACGTCAGGCAGCGGCGCAGGCGGCTGCCGGGGACTCACTGGATAAAGCAACGTTGAAAAAGGTTGCGCCGAAGCCTGGCTGGCTGGAAACCGGTGCTTCTGTTATGCCGGTACTGATGATCGTATTGATTGTGCGTTCGTTTATTTATGAACCGTTCCAGATCCCGTCAGGTTCGATGATGCCGACTCTGAACTCTACTGATTTTATTCTGGTAGAGAAGTTTGCTTATGGCATGAAAGATCCTATGTACCAGAAAACGATGATCGAAACCGGTCATCCGAAACGCGGCGATATCATGATGTTTAAATATCCGGAAGATCCAAAGCTTGATTACATCAAGCGCGCGGTGGGTTTACCGGGCGATAAAGTCACTTACGATCCGGTCTCAAAAGAGATGACGATGCAACCGGGATGCAGTTCCGGCCAGGCGTGTGAAAACGCGCTGCCGGTCACCTACTCAAACGTGGAACCGAGCGATTTCGTTCAGACCTTCTCACGCCGTAATGGTGGGGAAGCGACCAGCGGATTCTTTGAAGTGCCGAAACAGGAAACCAAAGAAAATGGAATTCGTCTTTCCGAGACTAGTGGTGGTCCAGGT**GCTCTGGCAGCTCTGGCACTGGCTGCATTAGCCGCTCTGGCCCTGGCCGCCTTAGCC**GGTCCTGGCGGTACGGTACCAGGTCAACAAAATGCAGTTTGGATTGTTCCTCCTGGACAATACTTCATGATGGGCGACAACCGCGACAACAGCGCGGACAGCCGTTACTGGGGCTTTGTGCCGGAAGCGAATCT**TTTTTTT**CGGGCAACGGCTATCTGGATGAGCTTCGAAAAGCAAGAAGGCGAATGGCCGACTGGTCTGCGCTTAAGTCGCATTGGCGGCATCCATCAGGAGGAGGAGGATCT**GCTACTAATTTTTCACTTCTTAAGCAAGCCGGGGATGTCGAAGAAAATCCGGGACCAATGGCCACAACCATGACGGCCCTGACAGAAGGTGCGAAGCTGTTCGAGAAGGAGATTCCCTATATCACAGAATTGGAGGGGGATGTAGAGGGTATGAAGTTTATCATCAAAGGCGAAGGGACAGGGGATGCAACAACTGGAACAATTAAGGCTAAGTACATTTGCACGACCGGCGACGTCCCGGTGCCCTGGTCCACGCTCGTCACCACGCTCACGTACGGAGCCCAGTGCTTTGCCAAATATGGCCCTGAACTTAAAGACTTCTACAAGTCGTGTATGCCGGAGGGATACGTGCAAGAGAGGACGATCACCTTTGAAGGTGACGGAGTATTCAAAACAAGAGCGGAGGTGACGTTCGAGAATGGATCGGTCTATAACCGGGTCAAGCTCAACGGACAGGGCTTTAAGAAAGATGGACACGTCCTTGGGAAGAATTTGGAGTTCAATTTCACCCCGCATTGTCTTTACATCTGGGGTGATCAGGCGAATCACGGGTTGAAATCAGCGTTCAAGATCATGCACGAGATTACGGGGAGCAAAGAGGACTTTATCGTGGCAGACCACACTCAGATGAACACTCCAATCGGAGGGGGTCCCGTACACGTACCCGAGTATCATCACCTGACCGTCTGGACATCGTTTGGAAAAGACCCTGACGACGATGAAACTGATCATCTCAACATTGTGGAAGTGATCAAGGCGGTGGACTTGGAAACATACCGGTGA**

**30. 3ʹ Ter Control, LepB Cellular Reporter (N_out_, LALA TMD, L=45)**

ATGGCGAATATGTTTGCCCTGATTCTGGTGATTGCCACACTGGTGACGGGCATTTTATGGTGCGTGGATAAATTCTTTTTCGCACCTAAACGGCGGGAACGTCAGGCAGCGGCGCAGGCGGCTGCCGGGGACTCACTGGATAAAGCAACGTTGAAAAAGGTTGCGCCGAAGCCTGGCTGGCTGGAAACCGGTGCTTCTGTTATGCCGGTACTGATGATCGTATTGATTGTGCGTTCGTTTATTTATGAACCGTTCCAGATCCCGTCAGGTTCGATGATGCCGACTCTGAACTCTACTGATTTTATTCTGGTAGAGAAGTTTGCTTATGGCATGAAAGATCCTATGTACCAGAAAACGATGATCGAAACCGGTCATCCGAAACGCGGCGATATCATGATGTTTAAATATCCGGAAGATCCAAAGCTTGATTACATCAAGCGCGCGGTGGGTTTACCGGGCGATAAAGTCACTTACGATCCGGTCTCAAAAGAGATGACGATGCAACCGGGATGCAGTTCCGGCCAGGCGTGTGAAAACGCGCTGCCGGTCACCTACTCAAACGTGGAACCGAGCGATTTCGTTCAGACCTTCTCACGCCGTAATGGTGGGGAAGCGACCAGCGGATTCTTTGAAGTGCCGAAACAGGAAACCAAAGAAAATGGAATTCGTCTTTCCGAGACTAGTGGTGGTCCAGGT**GCTCTGGCAGCTCTGGCACTGGCTGCATTAGCCGCTCTGGCCCTGGCCGCCTTAGCC**GGTCCTGGCGGTACGGTACCAGGTCAACAAAATGCAGTTTGGATTGTTCCTCCTGGACAATACTTCATGATGGGCGACAACCGCGACAACAGCGCGGACAGCCGTTACTGGGGCTTTGTGCCGGAAGCGAATCT**TTTTTTT**CGGGCAACGGCTATCTGGATGAGCTTCGAAAAGCAAGAAGG**TGA**ATGGCCGACTGGTCTGCGCTTAAGTCGCATTGGCGGCATCCATCAGGAGGAGGAGGATCT**GCTACTAATTTTTCACTTCTTAAGCAAGCCGGGGATGTCGAAGAAAATCCGGGACCAATGGCCACAACCATGACGGCCCTGACAGAAGGTGCGAAGCTGTTCGAGAAGGAGATTCCCTATATCACAGAATTGGAGGGGGATGTAGAGGGTATGAAGTTTATCATCAAAGGCGAAGGGACAGGGGATGCAACAACTGGAACAATTAAGGCTAAGTACATTTGCACGACCGGCGACGTCCCGGTGCCCTGGTCCACGCTCGTCACCACGCTCACGTACGGAGCCCAGTGCTTTGCCAAATATGGCCCTGAACTTAAAGACTTCTACAAGTCGTGTATGCCGGAGGGATACGTGCAAGAGAGGACGATCACCTTTGAAGGTGACGGAGTATTCAAAACAAGAGCGGAGGTGACGTTCGAGAATGGATCGGTCTATAACCGGGTCAAGCTCAACGGACAGGGCTTTAAGAAAGATGGACACGTCCTTGGGAAGAATTTGGAGTTCAATTTCACCCCGCATTGTCTTTACATCTGGGGTGATCAGGCGAATCACGGGTTGAAATCAGCGTTCAAGATCATGCACGAGATTACGGGGAGCAAAGAGGACTTTATCGTGGCAGACCACACTCAGATGAACACTCCAATCGGAGGGGGTCCCGTACACGTACCCGAGTATCATCACCTGACCGTCTGGACATCGTTTGGAAAAGACCCTGACGACGATGAAACTGATCATCTCAACATTGTGGAAGTGATCAAGGCGGTGGACTTGGAAACATACCGGTGA**

**31. 5ʹTer Control, LepB Cellular Reporter (N_out_, LALA TMD, L=45)**

ATGGCGAATATGTTTGCCCTGATTCTGGTGATTGCCACACTGGTGACGGGCATTTTATGGTGCGTGGATAAATTCTTTTTCGCACCTAAACGGCGGGAACGTCAGGCAGCGGCGCAGGCGGCTGCCGGGGACTCACTGGATAAAGCAACGTTGAAAAAGGTTGCGCCGAAGCCTGGCTGGCTGGAAACCGGTGCTTCTGTTATGCCGGTACTGATGATCGTATTGATTGTGCGTTCGTTTATTTATGAACCGTTCCAGATCCCGTCAGGTTCGATGATGCCGACTCTGAACTCTACTGATTTTATTCTGGTAGAGAAGTTTGCTTATGGCATGAAAGATCCTATGTACCAGAAAACGATGATCGAAACCGGTCATCCGAAACGCGGCGATATCATGATGTTTAAATATCCGGAAGATCCAAAGCTTGATTACATCAAGCGCGCGGTGGGTTTACCGGGCGATAAAGTCACTTACGATCCGGTCTCAAAAGAGATGACGATGCAACCGGGATGCAGTTCCGGCCAGGCGTGTGAAAACGCGCTGCCGGTCACCTACTCAAACGTGGAACCGAGCGATTTCGTTCAGACCTTCTCACGCCGTAATGGTGGGGAAGCGACCAGCGGATTCTTTGAAGTGCCGAAACAGGAAACCAAAGAAAATGGAATTCGTCTTTCCGAGACTAGTGGTGGTCCAGGT**GCTCTGGCAGCTCTGGCACTGGCTGCATTAGCCGCTCTGGCCCTGGCCGCCTTAGCC**GGTCCTGGCGGTACGGTACCAGGTCAACAAAATGCAGTTTGGATTGTTCCTCCTGGA**TAA**TACTTCATGATGGGCGACAACCGCGACAACAGCGCGGACAGCCGTTACTGGGGCTTTGTGCCGGAAGCGAATCT**TTTTTTT**CGGGCAACGGCTATCTGGATGAGCTTCGAAAAGCAAGAAGGCGAATGGCCGACTGGTCTGCGCTTAAGTCGCATTGGCGGCATCCATCAGGAGGAGGAGGATCT**GCTACTAATTTTTCACTTCTTAAGCAAGCCGGGGATGTCGAAGAAAATCCGGGACCAATGGCCACAACCATGACGGCCCTGACAGAAGGTGCGAAGCTGTTCGAGAAGGAGATTCCCTATATCACAGAATTGGAGGGGGATGTAGAGGGTATGAAGTTTATCATCAAAGGCGAAGGGACAGGGGATGCAACAACTGGAACAATTAAGGCTAAGTACATTTGCACGACCGGCGACGTCCCGGTGCCCTGGTCCACGCTCGTCACCACGCTCACGTACGGAGCCCAGTGCTTTGCCAAATATGGCCCTGAACTTAAAGACTTCTACAAGTCGTGTATGCCGGAGGGATACGTGCAAGAGAGGACGATCACCTTTGAAGGTGACGGAGTATTCAAAACAAGAGCGGAGGTGACGTTCGAGAATGGATCGGTCTATAACCGGGTCAAGCTCAACGGACAGGGCTTTAAGAAAGATGGACACGTCCTTGGGAAGAATTTGGAGTTCAATTTCACCCCGCATTGTCTTTACATCTGGGGTGATCAGGCGAATCACGGGTTGAAATCAGCGTTCAAGATCATGCACGAGATTACGGGGAGCAAAGAGGACTTTATCGTGGCAGACCACACTCAGATGAACACTCCAATCGGAGGGGGTCCCGTACACGTACCCGAGTATCATCACCTGACCGTCTGGACATCGTTTGGAAAAGACCCTGACGACGATGAAACTGATCATCTCAACATTGTGGAAGTGATCAAGGCGGTGGACTTGGAAACATACCGGTGA**

**32. SS_mut_ Control, LepB Cellular Reporter (N_out_, LALA TMD, L=45)**

ATGGCGAATATGTTTGCCCTGATTCTGGTGATTGCCACACTGGTGACGGGCATTTTATGGTGCGTGGATAAATTCTTTTTCGCACCTAAACGGCGGGAACGTCAGGCAGCGGCGCAGGCGGCTGCCGGGGACTCACTGGATAAAGCAACGTTGAAAAAGGTTGCGCCGAAGCCTGGCTGGCTGGAAACCGGTGCTTCTGTTATGCCGGTACTGATGATCGTATTGATTGTGCGTTCGTTTATTTATGAACCGTTCCAGATCCCGTCAGGTTCGATGATGCCGACTCTGAACTCTACTGATTTTATTCTGGTAGAGAAGTTTGCTTATGGCATGAAAGATCCTATGTACCAGAAAACGATGATCGAAACCGGTCATCCGAAACGCGGCGATATCATGATGTTTAAATATCCGGAAGATCCAAAGCTTGATTACATCAAGCGCGCGGTGGGTTTACCGGGCGATAAAGTCACTTACGATCCGGTCTCAAAAGAGATGACGATGCAACCGGGATGCAGTTCCGGCCAGGCGTGTGAAAACGCGCTGCCGGTCACCTACTCAAACGTGGAACCGAGCGATTTCGTTCAGACCTTCTCACGCCGTAATGGTGGGGAAGCGACCAGCGGATTCTTTGAAGTGCCGAAACAGGAAACCAAAGAAAATGGAATTCGTCTTTCCGAGACTAGTGGTGGTCCAGGT**GCTCTGGCAGCTCTGGCACTGGCTGCATTAGCCGCTCTGGCCCTGGCCGCCTTAGCC**GGTCCTGGCGGTACGGTACCAGGTCAACAAAATGCAGTTTGGATTGTTCCTCCTGGACAATACTTCATGATGGGCGACAACCGCGACAACAGCGCGGACAGCCGTTACTGGGGCTTTGTGCCGGAAGCGAATCT**GTTCTTC**CGGGCAACGGCTATCTGGATGAGCTTCGAAAAGCAAGAAGGCGAATGGCCGACTGGTCTGCGCTTAAGTCGCATTGGCGGCATCCATCAGGAGGAGGAGGATCT**GCTACTAATTTTTCACTTCTTAAGCAAGCCGGGGATGTCGAAGAAAATCCGGGACCAATGGCCACAACCATGACGGCCCTGACAGAAGGTGCGAAGCTGTTCGAGAAGGAGATTCCCTATATCACAGAATTGGAGGGGGATGTAGAGGGTATGAAGTTTATCATCAAAGGCGAAGGGACAGGGGATGCAACAACTGGAACAATTAAGGCTAAGTACATTTGCACGACCGGCGACGTCCCGGTGCCCTGGTCCACGCTCGTCACCACGCTCACGTACGGAGCCCAGTGCTTTGCCAAATATGGCCCTGAACTTAAAGACTTCTACAAGTCGTGTATGCCGGAGGGATACGTGCAAGAGAGGACGATCACCTTTGAAGGTGACGGAGTATTCAAAACAAGAGCGGAGGTGACGTTCGAGAATGGATCGGTCTATAACCGGGTCAAGCTCAACGGACAGGGCTTTAAGAAAGATGGACACGTCCTTGGGAAGAATTTGGAGTTCAATTTCACCCCGCATTGTCTTTACATCTGGGGTGATCAGGCGAATCACGGGTTGAAATCAGCGTTCAAGATCATGCACGAGATTACGGGGAGCAAAGAGGACTTTATCGTGGCAGACCACACTCAGATGAACACTCCAATCGGAGGGGGTCCCGTACACGTACCCGAGTATCATCACCTGACCGTCTGGACATCGTTTGGAAAAGACCCTGACGACGATGAAACTGATCATCTCAACATTGTGGAAGTGATCAAGGCGGTGGACTTGGAAACATACCGGTGA**

**33. 0-Frame Control, LepB Cellular Reporter (N_out_, LALA TMD, L=45)**

ATGGCGAATATGTTTGCCCTGATTCTGGTGATTGCCACACTGGTGACGGGCATTTTATGGTGCGTGGATAAATTCTTTTTCGCACCTAAACGGCGGGAACGTCAGGCAGCGGCGCAGGCGGCTGCCGGGGACTCACTGGATAAAGCAACGTTGAAAAAGGTTGCGCCGAAGCCTGGCTGGCTGGAAACCGGTGCTTCTGTTATGCCGGTACTGATGATCGTATTGATTGTGCGTTCGTTTATTTATGAACCGTTCCAGATCCCGTCAGGTTCGATGATGCCGACTCTGAACTCTACTGATTTTATTCTGGTAGAGAAGTTTGCTTATGGCATGAAAGATCCTATGTACCAGAAAACGATGATCGAAACCGGTCATCCGAAACGCGGCGATATCATGATGTTTAAATATCCGGAAGATCCAAAGCTTGATTACATCAAGCGCGCGGTGGGTTTACCGGGCGATAAAGTCACTTACGATCCGGTCTCAAAAGAGATGACGATGCAACCGGGATGCAGTTCCGGCCAGGCGTGTGAAAACGCGCTGCCGGTCACCTACTCAAACGTGGAACCGAGCGATTTCGTTCAGACCTTCTCACGCCGTAATGGTGGGGAAGCGACCAGCGGATTCTTTGAAGTGCCGAAACAGGAAACCAAAGAAAATGGAATTCGTCTTTCCGAGACTAGTGGTGGTCCAGGT**GCTCTGGCAGCTCTGGCACTGGCTGCATTAGCCGCTCTGGCCCTGGCCGCCTTAGCC**GGTCCTGGCGGTACGGTACCAGGTCAACAAAATGCAGTTTGGATTGTTCCTCCTGGACAATACTTCATGATGGGCGACAACCGCGACAACAGCGCGGACAGCCGTTACTGGGGCTTTGTGCCGGAAGCGAATCT**GTTCTTCT**CGGGCAACGGCTATCTGGA**TGA**GCTTCGAAAAGCAAGAAGGCGAATGGCCGACTGGTCTGCGCTTAAGTCGCATTGGCGGCATCCATCAGGAGGAGGAGGATCT**GCTACTAATTTTTCACTTCTTAAGCAAGCCGGGGATGTCGAAGAAAATCCGGGACCAATGGCCACAACCATGACGGCCCTGACAGAAGGTGCGAAGCTGTTCGAGAAGGAGATTCCCTATATCACAGAATTGGAGGGGGATGTAGAGGGTATGAAGTTTATCATCAAAGGCGAAGGGACAGGGGATGCAACAACTGGAACAATTAAGGCTAAGTACATTTGCACGACCGGCGACGTCCCGGTGCCCTGGTCCACGCTCGTCACCACGCTCACGTACGGAGCCCAGTGCTTTGCCAAATATGGCCCTGAACTTAAAGACTTCTACAAGTCGTGTATGCCGGAGGGATACGTGCAAGAGAGGACGATCACCTTTGAAGGTGACGGAGTATTCAAAACAAGAGCGGAGGTGACGTTCGAGAATGGATCGGTCTATAACCGGGTCAAGCTCAACGGACAGGGCTTTAAGAAAGATGGACACGTCCTTGGGAAGAATTTGGAGTTCAATTTCACCCCGCATTGTCTTTACATCTGGGGTGATCAGGCGAATCACGGGTTGAAATCAGCGTTCAAGATCATGCACGAGATTACGGGGAGCAAAGAGGACTTTATCGTGGCAGACCACACTCAGATGAACACTCCAATCGGAGGGGGTCCCGTACACGTACCCGAGTATCATCACCTGACCGTCTGGACATCGTTTGGAAAAGACCCTGACGACGATGAAACTGATCATCTCAACATTGTGGAAGTGATCAAGGCGGTGGACTTGGAAACATACCGGTGA**

**34. LepB Cellular Reporter (N_out_, LALA TMD, L=55)**

ATGGCGAATATGTTTGCCCTGATTCTGGTGATTGCCACACTGGTGACGGGCATTTTATGGTGCGTGGATAAATTCTTTTTCGCACCTAAACGGCGGGAACGTCAGGCAGCGGCGCAGGCGGCTGCCGGGGACTCACTGGATAAAGCAACGTTGAAAAAGGTTGCGCCGAAGCCTGGCTGGCTGGAAACCGGTGCTTCTGTTATGCCGGTACTGATGATCGTATTGATTGTGCGTTCGTTTATTTATGAACCGTTCCAGATCCCGTCAGGTTCGATGATGCCGACTCTGAACTCTACTGATTTTATTCTGGTAGAGAAGTTTGCTTATGGCATGAAAGATCCTATGTACCAGAAAACGATGATCGAAACCGGTCATCCGAAACGCGGCGATATCATGATGTTTAAATATCCGGAAGATCCAAAGCTTGATTACATCAAGCGCGCGGTGGGTTTACCGGGCGATAAAGTCACTTACGATCCGGTCTCAAAAGAGATGACGATGCAACCGGGATGCAGTTCCGGCCAGGCGTGTGAAAACGCGCTGCCGGTCACCTACTCAAACGTGGAACCGAGCGATTTCGTTCAGACCTTCTCACGCCGTAATGGTGGGGAAGCGACCAGCGGATTCTTTGAAGTGCCGAAACAGGAAACCAAAGAAAATGGAATTCGTCTTTCCGAGACTAGTGGTGGTCCAGGT**GCTCTGGCAGCTCTGGCACTGGCTGCATTAGCCGCTCTGGCCCTGGCCGCCTTAGCC**GGTCCTGGCGGTACGGTACCAGGTCAACAAAATGCAGTTTGGATTGTTCCTCCTGGACAATACTTCATGATGGGCGACAACCGCGACAACAGCGCGGACAGCCGTTACTGGGGCTTTGTGCCGGAAGCGAATCTGGTCGGTCGGGCAACGGCTATCTGGATGAG**TTTTTTT**ACGCAAGAAGGCGAATGGCCGACTGGTCTGCGCTTAAGTCGCATTGGCGGCATCCATCAGGAGGAGGAGGATCT**GCTACTAATTTTTCACTTCTTAAGCAAGCCGGGGATGTCGAAGAAAATCCGGGACCAATGGCCACAACCATGACGGCCCTGACAGAAGGTGCGAAGCTGTTCGAGAAGGAGATTCCCTATATCACAGAATTGGAGGGGGATGTAGAGGGTATGAAGTTTATCATCAAAGGCGAAGGGACAGGGGATGCAACAACTGGAACAATTAAGGCTAAGTACATTTGCACGACCGGCGACGTCCCGGTGCCCTGGTCCACGCTCGTCACCACGCTCACGTACGGAGCCCAGTGCTTTGCCAAATATGGCCCTGAACTTAAAGACTTCTACAAGTCGTGTATGCCGGAGGGATACGTGCAAGAGAGGACGATCACCTTTGAAGGTGACGGAGTATTCAAAACAAGAGCGGAGGTGACGTTCGAGAATGGATCGGTCTATAACCGGGTCAAGCTCAACGGACAGGGCTTTAAGAAAGATGGACACGTCCTTGGGAAGAATTTGGAGTTCAATTTCACCCCGCATTGTCTTTACATCTGGGGTGATCAGGCGAATCACGGGTTGAAATCAGCGTTCAAGATCATGCACGAGATTACGGGGAGCAAAGAGGACTTTATCGTGGCAGACCACACTCAGATGAACACTCCAATCGGAGGGGGTCCCGTACACGTACCCGAGTATCATCACCTGACCGTCTGGACATCGTTTGGAAAAGACCCTGACGACGATGAAACTGATCATCTCAACATTGTGGAAGTGATCAAGGCGGTGGACTTGGAAACATACCGGTGA**

**35. SS_mut_ Control, LepB Cellular Reporter (N_out_, LALA TMD, L=55)**

ATGGCGAATATGTTTGCCCTGATTCTGGTGATTGCCACACTGGTGACGGGCATTTTATGGTGCGTGGATAAATTCTTTTTCGCACCTAAACGGCGGGAACGTCAGGCAGCGGCGCAGGCGGCTGCCGGGGACTCACTGGATAAAGCAACGTTGAAAAAGGTTGCGCCGAAGCCTGGCTGGCTGGAAACCGGTGCTTCTGTTATGCCGGTACTGATGATCGTATTGATTGTGCGTTCGTTTATTTATGAACCGTTCCAGATCCCGTCAGGTTCGATGATGCCGACTCTGAACTCTACTGATTTTATTCTGGTAGAGAAGTTTGCTTATGGCATGAAAGATCCTATGTACCAGAAAACGATGATCGAAACCGGTCATCCGAAACGCGGCGATATCATGATGTTTAAATATCCGGAAGATCCAAAGCTTGATTACATCAAGCGCGCGGTGGGTTTACCGGGCGATAAAGTCACTTACGATCCGGTCTCAAAAGAGATGACGATGCAACCGGGATGCAGTTCCGGCCAGGCGTGTGAAAACGCGCTGCCGGTCACCTACTCAAACGTGGAACCGAGCGATTTCGTTCAGACCTTCTCACGCCGTAATGGTGGGGAAGCGACCAGCGGATTCTTTGAAGTGCCGAAACAGGAAACCAAAGAAAATGGAATTCGTCTTTCCGAGACTAGTGGTGGTCCAGGT**GCTCTGGCAGCTCTGGCACTGGCTGCATTAGCCGCTCTGGCCCTGGCCGCCTTAGCC**GGTCCTGGCGGTACGGTACCAGGTCAACAAAATGCAGTTTGGATTGTTCCTCCTGGACAATACTTCATGATGGGCGACAACCGCGACAACAGCGCGGACAGCCGTTACTGGGGCTTTGTGCCGGAAGCGAATCTGGTCGGTCGGGCAACGGCTATCTGGATGTC**GTTCTTC**ACGCAAGAAGGCGAATGGCCGACTGGTCTGCGCTTAAGTCGCATTGGCGGCATCCATCAGGAGGAGGAGGATCT**GCTACTAATTTTTCACTTCTTAAGCAAGCCGGGGATGTCGAAGAAAATCCGGGACCAATGGCCACAACCATGACGGCCCTGACAGAAGGTGCGAAGCTGTTCGAGAAGGAGATTCCCTATATCACAGAATTGGAGGGGGATGTAGAGGGTATGAAGTTTATCATCAAAGGCGAAGGGACAGGGGATGCAACAACTGGAACAATTAAGGCTAAGTACATTTGCACGACCGGCGACGTCCCGGTGCCCTGGTCCACGCTCGTCACCACGCTCACGTACGGAGCCCAGTGCTTTGCCAAATATGGCCCTGAACTTAAAGACTTCTACAAGTCGTGTATGCCGGAGGGATACGTGCAAGAGAGGACGATCACCTTTGAAGGTGACGGAGTATTCAAAACAAGAGCGGAGGTGACGTTCGAGAATGGATCGGTCTATAACCGGGTCAAGCTCAACGGACAGGGCTTTAAGAAAGATGGACACGTCCTTGGGAAGAATTTGGAGTTCAATTTCACCCCGCATTGTCTTTACATCTGGGGTGATCAGGCGAATCACGGGTTGAAATCAGCGTTCAAGATCATGCACGAGATTACGGGGAGCAAAGAGGACTTTATCGTGGCAGACCACACTCAGATGAACACTCCAATCGGAGGGGGTCCCGTACACGTACCCGAGTATCATCACCTGACCGTCTGGACATCGTTTGGAAAAGACCCTGACGACGATGAAACTGATCATCTCAACATTGTGGAAGTGATCAAGGCGGTGGACTTGGAAACATACCGGTGA**

**36. LepB Cellular Reporter (N_in_, LALA TMD, L=45)**

ATGGCGAATATGTTTGCCCTGATTCTGGTGATTGCCACACTGGTGACGGGCATTTTATGGTGCGTGGATAAATTCTTTTTCGCACCTAAACGGCGGGAACGTCAGGCAGCGGCGCAGGCGGCTGCCGGGGACTCACTGGATAAAGCAACGTTGAAAAAGGTTGCGCCGAAGCCTGGCTGGCTGGAAACCGGTGCTTCTGTTATGCCGGTACTGATGATCGTATTGATTGTGCGTTCGTTTATTTATGAACCGTTCCAGATCCCGTCAGGTTCGATGATGCCGACTCTGAACTCTACTGATTTTATTCTGGTAGAGAAGTTTGCTTATGGCATGAAAGATCCTATGTACCAGAAAACGATGATCGAAACCGGTCATCCGAAACGCGGCGATATCATGATGTTTAAATATCCGGAAGATCCAAAGCTTGATTACATCAAGCGCGCGGTGGGTTTACCGGGCGATAAAGTCACTTACGATCCGGTCTCAAAAGAGATGACGATGCAACCGGGATGCAGTTCCGGCCAGGCGTGTGAAAACGCGCTGCCG**GGTGGTCCAGGCGCATTAGCAGCACTGGCACTGGCGGCGCTGGCTGCACTGGCATTAGCGGCACTGGCGGGTCCAGGCGGC**AGCGATTTCGTTCAGACCTTCTCACGCCGTAATGGTGGGGAAGCGACCAGCGGATTCTTTGAAGTGCCGAAACAGGAAACCAAAGAAAATGGAATTCGTCTTTCCGAGACTAGTGGTGGTCCAGGT**GCTCTGGCAGCTCTGGCACTGGCTGCATTAGCCGCTCTGGCCCTGGCCGCCTTAGCC**GGTCCTGGCGGTACGGTACCAGGTCAACAAAATGCAGTTTGGATTGTTCCTCCTGGACAATACTTCATGATGGGCGACAACCGCGACAACAGCGCGGACAGCCGTTACTGGGGCTTTGTGCCGGAAGCGAATCT**TTTTTTT**CGGGCAACGGCTATCTGGATGAGCTTCGAAAAGCAAGAAGGCGAATGGCCGACTGGTCTGCGCTTAAGTCGCATTGGCGGCATCCATCAGGAGGAGGAGGATCT**GCTACTAATTTTTCACTTCTTAAGCAAGCCGGGGATGTCGAAGAAAATCCGGGACCAATGGCCACAACCATGACGGCCCTGACAGAAGGTGCGAAGCTGTTCGAGAAGGAGATTCCCTATATCACAGAATTGGAGGGGGATGTAGAGGGTATGAAGTTTATCATCAAAGGCGAAGGGACAGGGGATGCAACAACTGGAACAATTAAGGCTAAGTACATTTGCACGACCGGCGACGTCCCGGTGCCCTGGTCCACGCTCGTCACCACGCTCACGTACGGAGCCCAGTGCTTTGCCAAATATGGCCCTGAACTTAAAGACTTCTACAAGTCGTGTATGCCGGAGGGATACGTGCAAGAGAGGACGATCACCTTTGAAGGTGACGGAGTATTCAAAACAAGAGCGGAGGTGACGTTCGAGAATGGATCGGTCTATAACCGGGTCAAGCTCAACGGACAGGGCTTTAAGAAAGATGGACACGTCCTTGGGAAGAATTTGGAGTTCAATTTCACCCCGCATTGTCTTTACATCTGGGGTGATCAGGCGAATCACGGGTTGAAATCAGCGTTCAAGATCATGCACGAGATTACGGGGAGCAAAGAGGACTTTATCGTGGCAGACCACACTCAGATGAACACTCCAATCGGAGGGGGTCCCGTACACGTACCCGAGTATCATCACCTGACCGTCTGGACATCGTTTGGAAAAGACCCTGACGACGATGAAACTGATCATCTCAACATTGTGGAAGTGATCAAGGCGGTGGACTTGGAAACATACCGGTGA**

**37. SS_mut_ Control, LepB Cellular Reporter (N_in_, LALA TMD, L=45)**

ATGGCGAATATGTTTGCCCTGATTCTGGTGATTGCCACACTGGTGACGGGCATTTTATGGTGCGTGGATAAATTCTTTTTCGCACCTAAACGGCGGGAACGTCAGGCAGCGGCGCAGGCGGCTGCCGGGGACTCACTGGATAAAGCAACGTTGAAAAAGGTTGCGCCGAAGCCTGGCTGGCTGGAAACCGGTGCTTCTGTTATGCCGGTACTGATGATCGTATTGATTGTGCGTTCGTTTATTTATGAACCGTTCCAGATCCCGTCAGGTTCGATGATGCCGACTCTGAACTCTACTGATTTTATTCTGGTAGAGAAGTTTGCTTATGGCATGAAAGATCCTATGTACCAGAAAACGATGATCGAAACCGGTCATCCGAAACGCGGCGATATCATGATGTTTAAATATCCGGAAGATCCAAAGCTTGATTACATCAAGCGCGCGGTGGGTTTACCGGGCGATAAAGTCACTTACGATCCGGTCTCAAAAGAGATGACGATGCAACCGGGATGCAGTTCCGGCCAGGCGTGTGAAAACGCGCTGCCG**GGTGGTCCAGGCGCATTAGCAGCACTGGCACTGGCGGCGCTGGCTGCACTGGCATTAGCGGCACTGGCGGGTCCAGGCGGC**AGCGATTTCGTTCAGACCTTCTCACGCCGTAATGGTGGGGAAGCGACCAGCGGATTCTTTGAAGTGCCGAAACAGGAAACCAAAGAAAATGGAATTCGTCTTTCCGAGACTAGTGGTGGTCCAGGT**GCTCTGGCAGCTCTGGCACTGGCTGCATTAGCCGCTCTGGCCCTGGCCGCCTTAGCC**GGTCCTGGCGGTACGGTACCAGGTCAACAAAATGCAGTTTGGATTGTTCCTCCTGGACAATACTTCATGATGGGCGACAACCGCGACAACAGCGCGGACAGCCGTTACTGGGGCTTTGTGCCGGAAGCGAATCT**GTTCTTC**CGGGCAACGGCTATCTGGATGAGCTTCGAAAAGCAAGAAGGCGAATGGCCGACTGGTCTGCGCTTAAGTCGCATTGGCGGCATCCATCAGGAGGAGGAGGATCT**GCTACTAATTTTTCACTTCTTAAGCAAGCCGGGGATGTCGAAGAAAATCCGGGACCAATGGCCACAACCATGACGGCCCTGACAGAAGGTGCGAAGCTGTTCGAGAAGGAGATTCCCTATATCACAGAATTGGAGGGGGATGTAGAGGGTATGAAGTTTATCATCAAAGGCGAAGGGACAGGGGATGCAACAACTGGAACAATTAAGGCTAAGTACATTTGCACGACCGGCGACGTCCCGGTGCCCTGGTCCACGCTCGTCACCACGCTCACGTACGGAGCCCAGTGCTTTGCCAAATATGGCCCTGAACTTAAAGACTTCTACAAGTCGTGTATGCCGGAGGGATACGTGCAAGAGAGGACGATCACCTTTGAAGGTGACGGAGTATTCAAAACAAGAGCGGAGGTGACGTTCGAGAATGGATCGGTCTATAACCGGGTCAAGCTCAACGGACAGGGCTTTAAGAAAGATGGACACGTCCTTGGGAAGAATTTGGAGTTCAATTTCACCCCGCATTGTCTTTACATCTGGGGTGATCAGGCGAATCACGGGTTGAAATCAGCGTTCAAGATCATGCACGAGATTACGGGGAGCAAAGAGGACTTTATCGTGGCAGACCACACTCAGATGAACACTCCAATCGGAGGGGGTCCCGTACACGTACCCGAGTATCATCACCTGACCGTCTGGACATCGTTTGGAAAAGACCCTGACGACGATGAAACTGATCATCTCAACATTGTGGAAGTGATCAAGGCGGTGGACTTGGAAACATACCGGTGA**

**38. SINV Cellular Reporter (L=45)**

ATGTCCGCAGCACCACTGGTCACGGCAATGTGTTTGCTCGGAAATGTGAGCTTCCCATGCGACCGCCCGCCCACATGCTATACCCGCGAACCTTCCAGAGCCCTCGACATCCTTGAAGAGAACGTGAACCATGAGGCCTACGATACCCTGCTCAATGCCATATTGCGGTGCGGATCGTCTGGCAGAAGCAAAAGAAGCGTCACTGACGACTTTACCCTGACCAGCCCCTACTTGGGCACATGCTCGTACTGCCACCATACTGAACCGTGCTTCAGCCCTGTTAAGATCGAGCAGGTCTGGGACGAAGCGGACGATAACACCATACGCATACAGACTTCCGCCCAGTTTGGATACGACCATAGCGGAGCAGCAAGCGCAAACAAGTACCGCTACATGTCGCTTAAGCAGGATCACACCGTTAAAGAAGGCACCATGGATGACATCAAGATTAGCACCTCAGGACCGTGTAGAAGGCTTAGCTACAAAGGATACTTTCTCCTCGCAAAATGCCCTCCAGGGGACAGCGTAACGGTTAGCATAGTGAGTAGCAACTCAGCAACGTCATGTACACTGGCCCGCAAGATAAAACCAAAATTCGTGGGACGGGAAAAATATGATCTACCTCCCGTTCACGGTAAAAAAATTCCTTGCACAGTGTACGACCGTCTGAAAGAAACAACTGCAGGCTACATCACTATGCACAGGCCGGGACCGCACGCTTATACATCCTACCTGGAAGAATCATCAGGGAAAGTTTACGCAAAGCCGCCATCTGGGAAGAACATTACGTATGAGTGCAAGTGCGGCGACTACAAGACCGGAACCGTTTCGACCCGCACCGAAATCACTGGTTGCACCGCCATCAAGCAGTGCGTCGCCTATAAGAGCGACCAAACGAAGTGGGTCTTCAACTCACCGGACTTGATCAGACATGACGACCACACGGCTCAAGGGAAATTGCATTTGCCTTTCAAGTTGATCCCGAGTACCTGCATGGTCCCTGTTGCCCACGCGCCGAATGTAATACATGGCTTTAAACACATCAGCCTCCAATTAGATACAGACCACTTGACATTGCTCACCACCAGGAGACTAGGGGCAAACCCGGAACCAACCACTGAATGGATCGTCGGAAAGACGGTCAGAAACTTCACCGTCGACCGAGATGGCCTGGAATACATATGGGGAAATCATGAGCCAGTGAGGGTCTATGCCCAAGAGTCAGCACCAGGAGACCCTCACGGATGGCCACACGAAATAGTACAGCATTACTACCATCGCCATCCTGTGTACACCATCTTAGCCGTCGCATCAGCTACCGTGGCGATGATGATTGGCGTAACTGTTGCAGTGTTATGTGCCTGTAAAGCGCGCCGTGAGTGC**CTGACGCCATACGCCCTGGCCCCAAACGCCGTAATCCCAACTTCGCTGGCACTCTTGTGCTGCGTTAGG**TCGGCCAATGCTGAAACGTTCACCGAGACCATGAGTTACTTGTGGTCGAACAGTCAGCCGTTCTTCTGGGTCCAGTTGTGCATACCTTTGGCCGCTTTCATCGTTCTAATGCGCTGCTGCTCCTGCTGCCTGCC**TTTTTTA**GTGGTTGCCGGCGCCTACCTGGCGAAGGTAGACGCCTACGAACATGCGACCACTGTTCCAAATGTGCCACAGATACCGCAGGAGGAGGAGGATCT**GCTACTAATTTTTCACTTCTTAAGCAAGCCGGGGATGTCGAAGAAAATCCGGGACCAATGGCCACAACCATGACGGCCCTGACAGAAGGTGCGAAGCTGTTCGAGAAGGAGATTCCCTATATCACAGAATTGGAGGGGGATGTAGAGGGTATGAAGTTTATCATCAAAGGCGAAGGGACAGGGGATGCAACAACTGGAACAATTAAGGCTAAGTACATTTGCACGACCGGCGACGTCCCGGTGCCCTGGTCCACGCTCGTCACCACGCTCACGTACGGAGCCCAGTGCTTTGCCAAATATGGCCCTGAACTTAAAGACTTCTACAAGTCGTGTATGCCGGAGGGATACGTGCAAGAGAGGACGATCACCTTTGAAGGTGACGGAGTATTCAAAACAAGAGCGGAGGTGACGTTCGAGAATGGATCGGTCTATAACCGGGTCAAGCTCAACGGACAGGGCTTTAAGAAAGATGGACACGTCCTTGGGAAGAATTTGGAGTTCAATTTCACCCCGCATTGTCTTTACATCTGGGGTGATCAGGCGAATCACGGGTTGAAATCAGCGTTCAAGATCATGCACGAGATTACGGGGAGCAAAGAGGACTTTATCGTGGCAGACCACACTCAGATGAACACTCCAATCGGAGGGGGTCCCGTACACGTACCCGAGTATCATCACCTGACCGTCTGGACATCGTTTGGAAAAGACCCTGACGACGATGAAACTGATCATCTCAACATTGTGGAAGTGATCAAGGCGGTGGACTTGGAAACATACCGGTGA**

**39. 3ʹ Ter Control, SINV Cellular Reporter (L=45)**

ATGTCCGCAGCACCACTGGTCACGGCAATGTGTTTGCTCGGAAATGTGAGCTTCCCATGCGACCGCCCGCCCACATGCTATACCCGCGAACCTTCCAGAGCCCTCGACATCCTTGAAGAGAACGTGAACCATGAGGCCTACGATACCCTGCTCAATGCCATATTGCGGTGCGGATCGTCTGGCAGAAGCAAAAGAAGCGTCACTGACGACTTTACCCTGACCAGCCCCTACTTGGGCACATGCTCGTACTGCCACCATACTGAACCGTGCTTCAGCCCTGTTAAGATCGAGCAGGTCTGGGACGAAGCGGACGATAACACCATACGCATACAGACTTCCGCCCAGTTTGGATACGACCATAGCGGAGCAGCAAGCGCAAACAAGTACCGCTACATGTCGCTTAAGCAGGATCACACCGTTAAAGAAGGCACCATGGATGACATCAAGATTAGCACCTCAGGACCGTGTAGAAGGCTTAGCTACAAAGGATACTTTCTCCTCGCAAAATGCCCTCCAGGGGACAGCGTAACGGTTAGCATAGTGAGTAGCAACTCAGCAACGTCATGTACACTGGCCCGCAAGATAAAACCAAAATTCGTGGGACGGGAAAAATATGATCTACCTCCCGTTCACGGTAAAAAAATTCCTTGCACAGTGTACGACCGTCTGAAAGAAACAACTGCAGGCTACATCACTATGCACAGGCCGGGACCGCACGCTTATACATCCTACCTGGAAGAATCATCAGGGAAAGTTTACGCAAAGCCGCCATCTGGGAAGAACATTACGTATGAGTGCAAGTGCGGCGACTACAAGACCGGAACCGTTTCGACCCGCACCGAAATCACTGGTTGCACCGCCATCAAGCAGTGCGTCGCCTATAAGAGCGACCAAACGAAGTGGGTCTTCAACTCACCGGACTTGATCAGACATGACGACCACACGGCTCAAGGGAAATTGCATTTGCCTTTCAAGTTGATCCCGAGTACCTGCATGGTCCCTGTTGCCCACGCGCCGAATGTAATACATGGCTTTAAACACATCAGCCTCCAATTAGATACAGACCACTTGACATTGCTCACCACCAGGAGACTAGGGGCAAACCCGGAACCAACCACTGAATGGATCGTCGGAAAGACGGTCAGAAACTTCACCGTCGACCGAGATGGCCTGGAATACATATGGGGAAATCATGAGCCAGTGAGGGTCTATGCCCAAGAGTCAGCACCAGGAGACCCTCACGGATGGCCACACGAAATAGTACAGCATTACTACCATCGCCATCCTGTGTACACCATCTTAGCCGTCGCATCAGCTACCGTGGCGATGATGATTGGCGTAACTGTTGCAGTGTTATGTGCCTGTAAAGCGCGCCGTGAGTGC**CTGACGCCATACGCCCTGGCCCCAAACGCCGTAATCCCAACTTCGCTGGCACTCTTGTGCTGCGTTAGG**TCGGCCAATGCTGAAACGTTCACCGAGACCATGAGTTACTTGTGGTCGAACAGTCAGCCGTTCTTCTGGGTCCAGTTGTGCATACCTTTGGCCGCTTTCATCGTTCTAATGCGCTGCTGCTCCTGCTGCCTGCCTTTTTTAGTGGTTGCCGGCGCCTACCTGGCGAAGGTAGACGCCTACGAACATGCGACCACTGTTCCAAATgtgccacagatacc**taa**gcaggaggaggaggATCT**GCTACTAATTTTTCACTTCTTAAGCAAGCCGGGGATGTCGAAGAAAATCCGGGACCAATGGCCACAACCATGACGGCCCTGACAGAAGGTGCGAAGCTGTTCGAGAAGGAGATTCCCTATATCACAGAATTGGAGGGGGATGTAGAGGGTATGAAGTTTATCATCAAAGGCGAAGGGACAGGGGATGCAACAACTGGAACAATTAAGGCTAAGTACATTTGCACGACCGGCGACGTCCCGGTGCCCTGGTCCACGCTCGTCACCACGCTCACGTACGGAGCCCAGTGCTTTGCCAAATATGGCCCTGAACTTAAAGACTTCTACAAGTCGTGTATGCCGGAGGGATACGTGCAAGAGAGGACGATCACCTTTGAAGGTGACGGAGTATTCAAAACAAGAGCGGAGGTGACGTTCGAGAATGGATCGGTCTATAACCGGGTCAAGCTCAACGGACAGGGCTTTAAGAAAGATGGACACGTCCTTGGGAAGAATTTGGAGTTCAATTTCACCCCGCATTGTCTTTACATCTGGGGTGATCAGGCGAATCACGGGTTGAAATCAGCGTTCAAGATCATGCACGAGATTACGGGGAGCAAAGAGGACTTTATCGTGGCAGACCACACTCAGATGAACACTCCAATCGGAGGGGGTCCCGTACACGTACCCGAGTATCATCACCTGACCGTCTGGACATCGTTTGGAAAAGACCCTGACGACGATGAAACTGATCATCTCAACATTGTGGAAGTGATCAAGGCGGTGGACTTGGAAACATACCGGTGA**

**40. 5ʹTer Control, SINV Cellular Reporter (L=45)**

ATGTCCGCAGCACCACTGGTCACGGCAATGTGTTTGCTCGGAAATGTGAGCTTCCCATGCGACCGCCCGCCCACATGCTATACCCGCGAACCTTCCAGAGCCCTCGACATCCTTGAAGAGAACGTGAACCATGAGGCCTACGATACCCTGCTCAATGCCATATTGCGGTGCGGATCGTCTGGCAGAAGCAAAAGAAGCGTCACTGACGACTTTACCCTGACCAGCCCCTACTTGGGCACATGCTCGTACTGCCACCATACTGAACCGTGCTTCAGCCCTGTTAAGATCGAGCAGGTCTGGGACGAAGCGGACGATAACACCATACGCATACAGACTTCCGCCCAGTTTGGATACGACCATAGCGGAGCAGCAAGCGCAAACAAGTACCGCTACATGTCGCTTAAGCAGGATCACACCGTTAAAGAAGGCACCATGGATGACATCAAGATTAGCACCTCAGGACCGTGTAGAAGGCTTAGCTACAAAGGATACTTTCTCCTCGCAAAATGCCCTCCAGGGGACAGCGTAACGGTTAGCATAGTGAGTAGCAACTCAGCAACGTCATGTACACTGGCCCGCAAGATAAAACCAAAATTCGTGGGACGGGAAAAATATGATCTACCTCCCGTTCACGGTAAAAAAATTCCTTGCACAGTGTACGACCGTCTGAAAGAAACAACTGCAGGCTACATCACTATGCACAGGCCGGGACCGCACGCTTATACATCCTACCTGGAAGAATCATCAGGGAAAGTTTACGCAAAGCCGCCATCTGGGAAGAACATTACGTATGAGTGCAAGTGCGGCGACTACAAGACCGGAACCGTTTCGACCCGCACCGAAATCACTGGTTGCACCGCCATCAAGCAGTGCGTCGCCTATAAGAGCGACCAAACGAAGTGGGTCTTCAACTCACCGGACTTGATCAGACATGACGACCACACGGCTCAAGGGAAATTGCATTTGCCTTTCAAGTTGATCCCGAGTACCTGCATGGTCCCTGTTGCCCACGCGCCGAATGTAATACATGGCTTTAAACACATCAGCCTCCAATTAGATACAGACCACTTGACATTGCTCACCACCAGGAGACTAGGGGCAAACCCGGAACCAACCACTGAATGGATCGTCGGAAAGACGGTCAGAAACTTCACCGTCGACCGAGATGGCCTGGAATACATATGGGGAAATCATGAGCCAGTGAGGGTCTATGCCCAAGAGTCAGCACCAGGAGACCCTCACGGATGGCCACACGAAATAGTACAGCATTACTACCATCGCCATCCTGTGTACACCATCTTAGCCGTCGCATCAGCTACCGTGGCGATGATGATTGGCGTAACTGTTGCAGTGTTATGTGCCTGTAAAGCGCGCCGTGAGTGC**CTGACGCCATACGCCCTGGCCCCAAACGCCGTAATCCCAACTTCGCTGGCACTCTTGTGCTGCGTTAGG**TCGGCCAATGCTGAAACGTTCACCGAGACCATGAGTTACTTGTGGtcgaacagtcagccg**taa**ttcttctgggtccagTTGTGCATACCTTTGGCCGCTTTCATCGTTCTAATGCGCTGCTGCTCCTGCTGCCTGCC**TTTTTTA**GTGGTTGCCGGCGCCTACCTGGCGAAGGTAGACGCCTACGAACATGCGACCACTGTTCCAAATGTGCCACAGATACCGCAGGAGGAGGAGGATCT**GCTACTAATTTTTCACTTCTTAAGCAAGCCGGGGATGTCGAAGAAAATCCGGGACCAATGGCCACAACCATGACGGCCCTGACAGAAGGTGCGAAGCTGTTCGAGAAGGAGATTCCCTATATCACAGAATTGGAGGGGGATGTAGAGGGTATGAAGTTTATCATCAAAGGCGAAGGGACAGGGGATGCAACAACTGGAACAATTAAGGCTAAGTACATTTGCACGACCGGCGACGTCCCGGTGCCCTGGTCCACGCTCGTCACCACGCTCACGTACGGAGCCCAGTGCTTTGCCAAATATGGCCCTGAACTTAAAGACTTCTACAAGTCGTGTATGCCGGAGGGATACGTGCAAGAGAGGACGATCACCTTTGAAGGTGACGGAGTATTCAAAACAAGAGCGGAGGTGACGTTCGAGAATGGATCGGTCTATAACCGGGTCAAGCTCAACGGACAGGGCTTTAAGAAAGATGGACACGTCCTTGGGAAGAATTTGGAGTTCAATTTCACCCCGCATTGTCTTTACATCTGGGGTGATCAGGCGAATCACGGGTTGAAATCAGCGTTCAAGATCATGCACGAGATTACGGGGAGCAAAGAGGACTTTATCGTGGCAGACCACACTCAGATGAACACTCCAATCGGAGGGGGTCCCGTACACGTACCCGAGTATCATCACCTGACCGTCTGGACATCGTTTGGAAAAGACCCTGACGACGATGAAACTGATCATCTCAACATTGTGGAAGTGATCAAGGCGGTGGACTTGGAAACATACCGGTGA**

**41. SS_mut_ Control, SINV Cellular Reporter (L=45)**

ATGTCCGCAGCACCACTGGTCACGGCAATGTGTTTGCTCGGAAATGTGAGCTTCCCATGCGACCGCCCGCCCACATGCTATACCCGCGAACCTTCCAGAGCCCTCGACATCCTTGAAGAGAACGTGAACCATGAGGCCTACGATACCCTGCTCAATGCCATATTGCGGTGCGGATCGTCTGGCAGAAGCAAAAGAAGCGTCACTGACGACTTTACCCTGACCAGCCCCTACTTGGGCACATGCTCGTACTGCCACCATACTGAACCGTGCTTCAGCCCTGTTAAGATCGAGCAGGTCTGGGACGAAGCGGACGATAACACCATACGCATACAGACTTCCGCCCAGTTTGGATACGACCATAGCGGAGCAGCAAGCGCAAACAAGTACCGCTACATGTCGCTTAAGCAGGATCACACCGTTAAAGAAGGCACCATGGATGACATCAAGATTAGCACCTCAGGACCGTGTAGAAGGCTTAGCTACAAAGGATACTTTCTCCTCGCAAAATGCCCTCCAGGGGACAGCGTAACGGTTAGCATAGTGAGTAGCAACTCAGCAACGTCATGTACACTGGCCCGCAAGATAAAACCAAAATTCGTGGGACGGGAAAAATATGATCTACCTCCCGTTCACGGTAAAAAAATTCCTTGCACAGTGTACGACCGTCTGAAAGAAACAACTGCAGGCTACATCACTATGCACAGGCCGGGACCGCACGCTTATACATCCTACCTGGAAGAATCATCAGGGAAAGTTTACGCAAAGCCGCCATCTGGGAAGAACATTACGTATGAGTGCAAGTGCGGCGACTACAAGACCGGAACCGTTTCGACCCGCACCGAAATCACTGGTTGCACCGCCATCAAGCAGTGCGTCGCCTATAAGAGCGACCAAACGAAGTGGGTCTTCAACTCACCGGACTTGATCAGACATGACGACCACACGGCTCAAGGGAAATTGCATTTGCCTTTCAAGTTGATCCCGAGTACCTGCATGGTCCCTGTTGCCCACGCGCCGAATGTAATACATGGCTTTAAACACATCAGCCTCCAATTAGATACAGACCACTTGACATTGCTCACCACCAGGAGACTAGGGGCAAACCCGGAACCAACCACTGAATGGATCGTCGGAAAGACGGTCAGAAACTTCACCGTCGACCGAGATGGCCTGGAATACATATGGGGAAATCATGAGCCAGTGAGGGTCTATGCCCAAGAGTCAGCACCAGGAGACCCTCACGGATGGCCACACGAAATAGTACAGCATTACTACCATCGCCATCCTGTGTACACCATCTTAGCCGTCGCATCAGCTACCGTGGCGATGATGATTGGCGTAACTGTTGCAGTGTTATGTGCCTGTAAAGCGCGCCGTGAGTGC**CTGACGCCATACGCCCTGGCCCCAAACGCCGTAATCCCAACTTCGCTGGCACTCTTGTGCTGCGTTAGG**TCGGCCAATGCTGAAACGTTCACCGAGACCATGAGTTACTTGTGGTCGAACAGTCAGCCGTTCTTCTGGGTCCAGTTGTGCATACCTTTGGCCGCTTTCATCGTTCTAATGCGCTGCTGCTCCTgctgcctgcc**gttccta**gtggttgccgGCGCCTACCTGGCGAAGGTAGACGCCTACGAACATGCGACCACTGTTCCAAATGTGCCACAGATACCGCAGGAGGAGGAGGATCT**GCTACTAATTTTTCACTTCTTAAGCAAGCCGGGGATGTCGAAGAAAATCCGGGACCAATGGCCACAACCATGACGGCCCTGACAGAAGGTGCGAAGCTGTTCGAGAAGGAGATTCCCTATATCACAGAATTGGAGGGGGATGTAGAGGGTATGAAGTTTATCATCAAAGGCGAAGGGACAGGGGATGCAACAACTGGAACAATTAAGGCTAAGTACATTTGCACGACCGGCGACGTCCCGGTGCCCTGGTCCACGCTCGTCACCACGCTCACGTACGGAGCCCAGTGCTTTGCCAAATATGGCCCTGAACTTAAAGACTTCTACAAGTCGTGTATGCCGGAGGGATACGTGCAAGAGAGGACGATCACCTTTGAAGGTGACGGAGTATTCAAAACAAGAGCGGAGGTGACGTTCGAGAATGGATCGGTCTATAACCGGGTCAAGCTCAACGGACAGGGCTTTAAGAAAGATGGACACGTCCTTGGGAAGAATTTGGAGTTCAATTTCACCCCGCATTGTCTTTACATCTGGGGTGATCAGGCGAATCACGGGTTGAAATCAGCGTTCAAGATCATGCACGAGATTACGGGGAGCAAAGAGGACTTTATCGTGGCAGACCACACTCAGATGAACACTCCAATCGGAGGGGGTCCCGTACACGTACCCGAGTATCATCACCTGACCGTCTGGACATCGTTTGGAAAAGACCCTGACGACGATGAAACTGATCATCTCAACATTGTGGAAGTGATCAAGGCGGTGGACTTGGAAACATACCGGTGA**

**42. 0-Frame Control, SINV Cellular Reporter (L=45)**

ATGTCCGCAGCACCACTGGTCACGGCAATGTGTTTGCTCGGAAATGTGAGCTTCCCATGCGACCGCCCGCCCACATGCTATACCCGCGAACCTTCCAGAGCCCTCGACATCCTTGAAGAGAACGTGAACCATGAGGCCTACGATACCCTGCTCAATGCCATATTGCGGTGCGGATCGTCTGGCAGAAGCAAAAGAAGCGTCACTGACGACTTTACCCTGACCAGCCCCTACTTGGGCACATGCTCGTACTGCCACCATACTGAACCGTGCTTCAGCCCTGTTAAGATCGAGCAGGTCTGGGACGAAGCGGACGATAACACCATACGCATACAGACTTCCGCCCAGTTTGGATACGACCATAGCGGAGCAGCAAGCGCAAACAAGTACCGCTACATGTCGCTTAAGCAGGATCACACCGTTAAAGAAGGCACCATGGATGACATCAAGATTAGCACCTCAGGACCGTGTAGAAGGCTTAGCTACAAAGGATACTTTCTCCTCGCAAAATGCCCTCCAGGGGACAGCGTAACGGTTAGCATAGTGAGTAGCAACTCAGCAACGTCATGTACACTGGCCCGCAAGATAAAACCAAAATTCGTGGGACGGGAAAAATATGATCTACCTCCCGTTCACGGTAAAAAAATTCCTTGCACAGTGTACGACCGTCTGAAAGAAACAACTGCAGGCTACATCACTATGCACAGGCCGGGACCGCACGCTTATACATCCTACCTGGAAGAATCATCAGGGAAAGTTTACGCAAAGCCGCCATCTGGGAAGAACATTACGTATGAGTGCAAGTGCGGCGACTACAAGACCGGAACCGTTTCGACCCGCACCGAAATCACTGGTTGCACCGCCATCAAGCAGTGCGTCGCCTATAAGAGCGACCAAACGAAGTGGGTCTTCAACTCACCGGACTTGATCAGACATGACGACCACACGGCTCAAGGGAAATTGCATTTGCCTTTCAAGTTGATCCCGAGTACCTGCATGGTCCCTGTTGCCCACGCGCCGAATGTAATACATGGCTTTAAACACATCAGCCTCCAATTAGATACAGACCACTTGACATTGCTCACCACCAGGAGACTAGGGGCAAACCCGGAACCAACCACTGAATGGATCGTCGGAAAGACGGTCAGAAACTTCACCGTCGACCGAGATGGCCTGGAATACATATGGGGAAATCATGAGCCAGTGAGGGTCTATGCCCAAGAGTCAGCACCAGGAGACCCTCACGGATGGCCACACGAAATAGTACAGCATTACTACCATCGCCATCCTGTGTACACCATCTTAGCCGTCGCATCAGCTACCGTGGCGATGATGATTGGCGTAACTGTTGCAGTGTTATGTGCCTGTAAAGCGCGCCGTGAGTGC**CTGACGCCATACGCCCTGGCCCCAAACGCCGTAATCCCAACTTCGCTGGCACTCTTGTGCTGCGTTAGG**TCGGCCAATGCTGAAACGTTCACCGAGACCATGAGTTACTTGTGGTCGAACAGTCAGCCGTTCTTCTGGGTCCAGTTGTGCATACCTTTGGCCGCTTTCATCGTTCTAATGCGCTGCTGCTCCTGCTGCCTGCC**TTTTTTA**GTGGTTGCCGGCGCCTACCTGGCGAAGGTAGACGCCTACGAACATGCGACCACTGTTCCAAATGTGccacagataccgca**g**ggaggaggaggatct**gCTACTAATTTTTCACTTCTTAAGCAAGCCGGGGATGTCGAAGAAAATCCGGGACCAATGGCCACAACCATGACGGCCCTGACAGAAGGTGCGAAGCTGTTCGAGAAGGAGATTCCCTATATCACAGAATTGGAGGGGGATGTAGAGGGTATGAAGTTTATCATCAAAGGCGAAGGGACAGGGGATGCAACAACTGGAACAATTAAGGCTAAGTACATTTGCACGACCGGCGACGTCCCGGTGCCCTGGTCCACGCTCGTCACCACGCTCACGTACGGAGCCCAGTGCTTTGCCAAATATGGCCCTGAACTTAAAGACTTCTACAAGTCGTGTATGCCGGAGGGATACGTGCAAGAGAGGACGATCACCTTTGAAGGTGACGGAGTATTCAAAACAAGAGCGGAGGTGACGTTCGAGAATGGATCGGTCTATAACCGGGTCAAGCTCAACGGACAGGGCTTTAAGAAAGATGGACACGTCCTTGGGAAGAATTTGGAGTTCAATTTCACCCCGCATTGTCTTTACATCTGGGGTGATCAGGCGAATCACGGGTTGAAATCAGCGTTCAAGATCATGCACGAGATTACGGGGAGCAAAGAGGACTTTATCGTGGCAGACCACACTCAGATGAACACTCCAATCGGAGGGGGTCCCGTACACGTACCCGAGTATCATCACCTGACCGTCTGGACATCGTTTGGAAAAGACCCTGACGACGATGAAACTGATCATCTCAACATTGTGGAAGTGATCAAGGCGGTGGACTTGGAAACATACCGGTGA**

**43. KCNQ1 Cellular Reporter (L=67)**

atggccgcggcctcctccccgcccagggccgagaggaagcgctggggttggggccgcctgccaggcgcccggcggggcagcgcgggcctggccaagaagtgccccttctcgctggagctggcggagggcggcccggcgggcggcgcgctctacgcgcccatcgcgcccggcgccccaggtcccgcgccccctgcgtccccggccgcgcccgccgcgcccccagttgcctccgaccttggcccgcggccgccggtgagcctagacccgcgcgtctccatctacagcacgcgccgcccggtgttggcgcgcacccacgtccagggccgcgtctacaacttcctcgagcgtcccaccggctggaaatgcttcgtttaccacttcgccgtcttcctcatcgtcctggtctgcctcatcttcagcgtgctgtccaccatcgagcagtatgccgccctggccacggggactctcttctggatggagatcgtgctggtggtgttcttcgggacggagtacgtggtccgcctctggtccgccggctgccgcagcaagtacgtgggcctctgggggcggctgcgctttgcccggaagcccatttccatcatcgacctcatcgtggtcgtggcctccatggtggtcctctgcgtgggctccaaggggcaggtgtttgccacgtcggccatcaggggcatccgcttcctgcagatcctgaggatgctacacgtcgaccgccagggaggcacctggaggctcctgggctccgtggtcttcatccaccgccaggagctgataaccaccctgtacatcggcttcctgggcctcatcttctcctcgtactttgtgtacctggctgagaaggacgcggtgaacgagtcaggccgcgtggagttcggcagctacgcagatgcgctgtggtggggggtggtcacagtcaccaccatcggctatggggacaaggtgccccagacgtgggtcgggaagaccatcgcctcc**tgcttctctgtctttgccatctccttctttgcgctcccagcggggattcttggctcggggtttgccctg**aaggtgcagcagaagcagaggcagaagcacttcaaccggcagatcccggcggcagcctcactcattcagaccgcatggaggtgctatgctgccgagaaccccgactcctccacctggaagatctacatccggaaggccccccggagccacactctgctgtcacccagccccaaacccaagaagtctgtggtggtaaagaa**aaaaaag**ttcaagctggacaaagacaatggggtgactcctggagagaagatgctcacagtcccccatatcacgtgcgaccccccagaagagcggcggctggaccacccGGAGGAGGAGGATCT**GCTACTAATTTTTCACTTCTTAAGCAAGCCGGGGATGTCGAAGAAAATCCGGGACCAATGGCCACAACCATGACGGCCCTGACAGAAGGTGCGAAGCTGTTCGAGAAGGAGATTCCCTATATCACAGAATTGGAGGGGGATGTAGAGGGTATGAAGTTTATCATCAAAGGCGAAGGGACAGGGGATGCAACAACTGGAACAATTAAGGCTAAGTACATTTGCACGACCGGCGACGTCCCGGTGCCCTGGTCCACGCTCGTCACCACGCTCACGTACGGAGCCCAGTGCTTTGCCAAATATGGCCCTGAACTTAAAGACTTCTACAAGTCGTGTATGCCGGAGGGATACGTGCAAGAGAGGACGATCACCTTTGAAGGTGACGGAGTATTCAAAACAAGAGCGGAGGTGACGTTCGAGAATGGATCGGTCTATAACCGGGTCAAGCTCAACGGACAGGGCTTTAAGAAAGATGGACACGTCCTTGGGAAGAATTTGGAGTTCAATTTCACCCCGCATTGTCTTTACATCTGGGGTGATCAGGCGAATCACGGGTTGAAATCAGCGTTCAAGATCATGCACGAGATTACGGGGAGCAAAGAGGACTTTATCGTGGCAGACCACACTCAGATGAACACTCCAATCGGAGGGGGTCCCGTACACGTACCCGAGTATCATCACCTGACCGTCTGGACATCGTTTGGAAAAGACCCTGACGACGATGAAACTGATCATCTCAACATTGTGGAAGTGATCAAGGCGGTGGACTTGGAAACATACCGGTGA**

**44. 3ʹ Ter Control, KCNQ1 Cellular Reporter (L=67)**

atggccgcggcctcctccccgcccagggccgagaggaagcgctggggttggggccgcctgccaggcgcccggcggggcagcgcgggcctggccaagaagtgccccttctcgctggagctggcggagggcggcccggcgggcggcgcgctctacgcgcccatcgcgcccggcgccccaggtcccgcgccccctgcgtccccggccgcgcccgccgcgcccccagttgcctccgaccttggcccgcggccgccggtgagcctagacccgcgcgtctccatctacagcacgcgccgcccggtgttggcgcgcacccacgtccagggccgcgtctacaacttcctcgagcgtcccaccggctggaaatgcttcgtttaccacttcgccgtcttcctcatcgtcctggtctgcctcatcttcagcgtgctgtccaccatcgagcagtatgccgccctggccacggggactctcttctggatggagatcgtgctggtggtgttcttcgggacggagtacgtggtccgcctctggtccgccggctgccgcagcaagtacgtgggcctctgggggcggctgcgctttgcccggaagcccatttccatcatcgacctcatcgtggtcgtggcctccatggtggtcctctgcgtgggctccaaggggcaggtgtttgccacgtcggccatcaggggcatccgcttcctgcagatcctgaggatgctacacgtcgaccgccagggaggcacctggaggctcctgggctccgtggtcttcatccaccgccaggagctgataaccaccctgtacatcggcttcctgggcctcatcttctcctcgtactttgtgtacctggctgagaaggacgcggtgaacgagtcaggccgcgtggagttcggcagctacgcagatgcgctgtggtggggggtggtcacagtcaccaccatcggctatggggacaaggtgccccagacgtgggtcgggaagaccatcgcctcc**tgcttctctgtctttgccatctccttctttgcgctcccagcggggattcttggctcggggtttgccctg**aaggtgcagcagaagcagaggcagaagcacttcaaccggcagatcccggcggcagcctcactcattcagaccgcatggaggtgctatgctgccgagaaccccgactcctccacctggaagatctacatccggaaggccccccggagccacactctgctgtcacccagccccaaacccaagaagtctgtggtggtaaagaa**aaaaaag**ttcaagctggacaaagacaatggggtgactcctggagagaagatgctcacagtcccccatatcacgtgcgaccccccagaagagcggcggctggaccaccc**taa**ggaggaggaggATCT**GCTACTAATTTTTCACTTCTTAAGCAAGCCGGGGATGTCGAAGAAAATCCGGGACCAATGGCCACAACCATGACGGCCCTGACAGAAGGTGCGAAGCTGTTCGAGAAGGAGATTCCCTATATCACAGAATTGGAGGGGGATGTAGAGGGTATGAAGTTTATCATCAAAGGCGAAGGGACAGGGGATGCAACAACTGGAACAATTAAGGCTAAGTACATTTGCACGACCGGCGACGTCCCGGTGCCCTGGTCCACGCTCGTCACCACGCTCACGTACGGAGCCCAGTGCTTTGCCAAATATGGCCCTGAACTTAAAGACTTCTACAAGTCGTGTATGCCGGAGGGATACGTGCAAGAGAGGACGATCACCTTTGAAGGTGACGGAGTATTCAAAACAAGAGCGGAGGTGACGTTCGAGAATGGATCGGTCTATAACCGGGTCAAGCTCAACGGACAGGGCTTTAAGAAAGATGGACACGTCCTTGGGAAGAATTTGGAGTTCAATTTCACCCCGCATTGTCTTTACATCTGGGGTGATCAGGCGAATCACGGGTTGAAATCAGCGTTCAAGATCATGCACGAGATTACGGGGAGCAAAGAGGACTTTATCGTGGCAGACCACACTCAGATGAACACTCCAATCGGAGGGGGTCCCGTACACGTACCCGAGTATCATCACCTGACCGTCTGGACATCGTTTGGAAAAGACCCTGACGACGATGAAACTGATCATCTCAACATTGTGGAAGTGATCAAGGCGGTGGACTTGGAAACATACCGGTGA**

**45. 5ʹTer Control, KCNQ1 Cellular Reporter (L=67)**

atggccgcggcctcctccccgcccagggccgagaggaagcgctggggttggggccgcctgccaggcgcccggcggggcagcgcgggcctggccaagaagtgccccttctcgctggagctggcggagggcggcccggcgggcggcgcgctctacgcgcccatcgcgcccggcgccccaggtcccgcgccccctgcgtccccggccgcgcccgccgcgcccccagttgcctccgaccttggcccgcggccgccggtgagcctagacccgcgcgtctccatctacagcacgcgccgcccggtgttggcgcgcacccacgtccagggccgcgtctacaacttcctcgagcgtcccaccggctggaaatgcttcgtttaccacttcgccgtcttcctcatcgtcctggtctgcctcatcttcagcgtgctgtccaccatcgagcagtatgccgccctggccacggggactctcttctggatggagatcgtgctggtggtgttcttcgggacggagtacgtggtccgcctctggtccgccggctgccgcagcaagtacgtgggcctctgggggcggctgcgctttgcccggaagcccatttccatcatcgacctcatcgtggtcgtggcctccatggtggtcctctgcgtgggctccaaggggcaggtgtttgccacgtcggccatcaggggcatccgcttcctgcagatcctgaggatgctacacgtcgaccgccagggaggcacctggaggctcctgggctccgtggtcttcatccaccgccaggagctgataaccaccctgtacatcggcttcctgggcctcatcttctcctcgtactttgtgtacctggctgagaaggacgcggtgaacgagtcaggccgcgtggagttcggcagctacgcagatgcgctgtggtggggggtggtcacagtcaccaccatcggctatggggacaaggtgccccagacgtgggtcgggaagaccatcgcctcc**tgcttctctgtctttgccatctccttctttgcgctcccagcggggattcttggctcggggtttgccctg**aaggtgcagcagaagcagaggcagaagcacttcaaccggcagatcccggcggcagcctcactcattcagaccgcatggaggtgctatgctgccgagaaccccgactcctccacctggaagatctacatccggaaggccccccggagccacactctgctgtcacccagcccc**taa**aaacccaagaagtctgtggtggtaaagaa**aaaaaag**ttcaagctggacaaagacaatggggtgactcctggagagaagatgctcacagtcccccatatcacgtgcgaccccccagaagagcggcggctggaccacccGGAGGAGGAGGATCT**GCTACTAATTTTTCACTTCTTAAGCAAGCCGGGGATGTCGAAGAAAATCCGGGACCAATGGCCACAACCATGACGGCCCTGACAGAAGGTGCGAAGCTGTTCGAGAAGGAGATTCCCTATATCACAGAATTGGAGGGGGATGTAGAGGGTATGAAGTTTATCATCAAAGGCGAAGGGACAGGGGATGCAACAACTGGAACAATTAAGGCTAAGTACATTTGCACGACCGGCGACGTCCCGGTGCCCTGGTCCACGCTCGTCACCACGCTCACGTACGGAGCCCAGTGCTTTGCCAAATATGGCCCTGAACTTAAAGACTTCTACAAGTCGTGTATGCCGGAGGGATACGTGCAAGAGAGGACGATCACCTTTGAAGGTGACGGAGTATTCAAAACAAGAGCGGAGGTGACGTTCGAGAATGGATCGGTCTATAACCGGGTCAAGCTCAACGGACAGGGCTTTAAGAAAGATGGACACGTCCTTGGGAAGAATTTGGAGTTCAATTTCACCCCGCATTGTCTTTACATCTGGGGTGATCAGGCGAATCACGGGTTGAAATCAGCGTTCAAGATCATGCACGAGATTACGGGGAGCAAAGAGGACTTTATCGTGGCAGACCACACTCAGATGAACACTCCAATCGGAGGGGGTCCCGTACACGTACCCGAGTATCATCACCTGACCGTCTGGACATCGTTTGGAAAAGACCCTGACGACGATGAAACTGATCATCTCAACATTGTGGAAGTGATCAAGGCGGTGGACTTGGAAACATACCGGTGA**

**46. SS_mut_ Control, KCNQ1 Cellular Reporter (L=67)**

atggccgcggcctcctccccgcccagggccgagaggaagcgctggggttggggccgcctgccaggcgcccggcggggcagcgcgggcctggccaagaagtgccccttctcgctggagctggcggagggcggcccggcgggcggcgcgctctacgcgcccatcgcgcccggcgccccaggtcccgcgccccctgcgtccccggccgcgcccgccgcgcccccagttgcctccgaccttggcccgcggccgccggtgagcctagacccgcgcgtctccatctacagcacgcgccgcccggtgttggcgcgcacccacgtccagggccgcgtctacaacttcctcgagcgtcccaccggctggaaatgcttcgtttaccacttcgccgtcttcctcatcgtcctggtctgcctcatcttcagcgtgctgtccaccatcgagcagtatgccgccctggccacggggactctcttctggatggagatcgtgctggtggtgttcttcgggacggagtacgtggtccgcctctggtccgccggctgccgcagcaagtacgtgggcctctgggggcggctgcgctttgcccggaagcccatttccatcatcgacctcatcgtggtcgtggcctccatggtggtcctctgcgtgggctccaaggggcaggtgtttgccacgtcggccatcaggggcatccgcttcctgcagatcctgaggatgctacacgtcgaccgccagggaggcacctggaggctcctgggctccgtggtcttcatccaccgccaggagctgataaccaccctgtacatcggcttcctgggcctcatcttctcctcgtactttgtgtacctggctgagaaggacgcggtgaacgagtcaggccgcgtggagttcggcagctacgcagatgcgctgtggtggggggtggtcacagtcaccaccatcggctatggggacaaggtgccccagacgtgggtcgggaagaccatcgcctcc**tgcttctctgtctttgccatctccttctttgcgctcccagcggggattcttggctcggggtttgccctg**aaggtgcagcagaagcagaggcagaagcacttcaaccggcagatcccggcggcagcctcactcattcagaccgcatggaggtgctatgctgccgagaaccccgactcctccacctggaagatctacatccggaaggccccccggagccacactctgctgtcacccagccccaaacccaagaagtctgtggtggta**aaggcgcgcgag**ttcaagctggacaaagacaatggggtgactcctggagagaagatgctcacagtcccccatatcacgtgcgaccccccagaagagcggcggctggaccacccGGAGGAGGAGGATCT**GCTACTAATTTTTCACTTCTTAAGCAAGCCGGGGATGTCGAAGAAAATCCGGGACCAATGGCCACAACCATGACGGCCCTGACAGAAGGTGCGAAGCTGTTCGAGAAGGAGATTCCCTATATCACAGAATTGGAGGGGGATGTAGAGGGTATGAAGTTTATCATCAAAGGCGAAGGGACAGGGGATGCAACAACTGGAACAATTAAGGCTAAGTACATTTGCACGACCGGCGACGTCCCGGTGCCCTGGTCCACGCTCGTCACCACGCTCACGTACGGAGCCCAGTGCTTTGCCAAATATGGCCCTGAACTTAAAGACTTCTACAAGTCGTGTATGCCGGAGGGATACGTGCAAGAGAGGACGATCACCTTTGAAGGTGACGGAGTATTCAAAACAAGAGCGGAGGTGACGTTCGAGAATGGATCGGTCTATAACCGGGTCAAGCTCAACGGACAGGGCTTTAAGAAAGATGGACACGTCCTTGGGAAGAATTTGGAGTTCAATTTCACCCCGCATTGTCTTTACATCTGGGGTGATCAGGCGAATCACGGGTTGAAATCAGCGTTCAAGATCATGCACGAGATTACGGGGAGCAAAGAGGACTTTATCGTGGCAGACCACACTCAGATGAACACTCCAATCGGAGGGGGTCCCGTACACGTACCCGAGTATCATCACCTGACCGTCTGGACATCGTTTGGAAAAGACCCTGACGACGATGAAACTGATCATCTCAACATTGTGGAAGTGATCAAGGCGGTGGACTTGGAAACATACCGGTGA**

**47. 0-Frame Control, KCNQ1 Cellular Reporter (L=67)**

atggccgcggcctcctccccgcccagggccgagaggaagcgctggggttggggccgcctgccaggcgcccggcggggcagcgcgggcctggccaagaagtgccccttctcgctggagctggcggagggcggcccggcgggcggcgcgctctacgcgcccatcgcgcccggcgccccaggtcccgcgccccctgcgtccccggccgcgcccgccgcgcccccagttgcctccgaccttggcccgcggccgccggtgagcctagacccgcgcgtctccatctacagcacgcgccgcccggtgttggcgcgcacccacgtccagggccgcgtctacaacttcctcgagcgtcccaccggctggaaatgcttcgtttaccacttcgccgtcttcctcatcgtcctggtctgcctcatcttcagcgtgctgtccaccatcgagcagtatgccgccctggccacggggactctcttctggatggagatcgtgctggtggtgttcttcgggacggagtacgtggtccgcctctggtccgccggctgccgcagcaagtacgtgggcctctgggggcggctgcgctttgcccggaagcccatttccatcatcgacctcatcgtggtcgtggcctccatggtggtcctctgcgtgggctccaaggggcaggtgtttgccacgtcggccatcaggggcatccgcttcctgcagatcctgaggatgctacacgtcgaccgccagggaggcacctggaggctcctgggctccgtggtcttcatccaccgccaggagctgataaccaccctgtacatcggcttcctgggcctcatcttctcctcgtactttgtgtacctggctgagaaggacgcggtgaacgagtcaggccgcgtggagttcggcagctacgcagatgcgctgtggtggggggtggtcacagtcaccaccatcggctatggggacaaggtgccccagacgtgggtcgggaagaccatcgcctcc**tgcttctctgtctttgccatctccttctttgcgctcccagcggggattcttggctcggggtttgccctg**aaggtgcagcagaagcagaggcagaagcacttcaaccggcagatcccggcggcagcctcactcattcagaccgcatggaggtgctatgctgccgagaaccccgactcctccacctggaagatctacatccggaaggccccccggagccacactctgctgtcacccagccccaaacccaagaagtctgtggtggtaaagaa**aaaaaag**ttcaagctggacaaagacaatggggtgactcctggagagaagatgctcacagtcccccatatcacgtgcgaccccccagaagagcggcggctggaccaccc**c**ggaggaggaggaTCT**GCTACTAATTTTTCACTTCTTAAGCAAGCCGGGGATGTCGAAGAAAATCCGGGACCAATGGCCACAACCATGACGGCCCTGACAGAAGGTGCGAAGCTGTTCGAGAAGGAGATTCCCTATATCACAGAATTGGAGGGGGATGTAGAGGGTATGAAGTTTATCATCAAAGGCGAAGGGACAGGGGATGCAACAACTGGAACAATTAAGGCTAAGTACATTTGCACGACCGGCGACGTCCCGGTGCCCTGGTCCACGCTCGTCACCACGCTCACGTACGGAGCCCAGTGCTTTGCCAAATATGGCCCTGAACTTAAAGACTTCTACAAGTCGTGTATGCCGGAGGGATACGTGCAAGAGAGGACGATCACCTTTGAAGGTGACGGAGTATTCAAAACAAGAGCGGAGGTGACGTTCGAGAATGGATCGGTCTATAACCGGGTCAAGCTCAACGGACAGGGCTTTAAGAAAGATGGACACGTCCTTGGGAAGAATTTGGAGTTCAATTTCACCCCGCATTGTCTTTACATCTGGGGTGATCAGGCGAATCACGGGTTGAAATCAGCGTTCAAGATCATGCACGAGATTACGGGGAGCAAAGAGGACTTTATCGTGGCAGACCACACTCAGATGAACACTCCAATCGGAGGGGGTCCCGTACACGTACCCGAGTATCATCACCTGACCGTCTGGACATCGTTTGGAAAAGACCCTGACGACGATGAAACTGATCATCTCAACATTGTGGAAGTGATCAAGGCGGTGGACTTGGAAACATACCGGTGA**

**48. KCNQ1 Alt-Splice Cellular Reporter (L=39)**

ATGGCCGCGGCCTCCTCCCCGCCCAGGGCCGAGAGGAAGCGCTGGGGTTGGGGCCGCCTGCCAGGCGCCCGGCGGGGCAGCGCGGGCCTGGCCAAGAAGTGCCCCTTCTCGCTGGAGCTGGCGGAGGGCGGCCCGGCGGGCGGCGCGCTCTACGCGCCCATCGCGCCCGGCGCCCCAGGTCCCGCGCCCCCTGCGTCCCCGGCCGCGCCCGCCGCGCCCCCAGTTGCCTCCGACCTTGGCCCGCGGCCGCCGGTGAGCCTAGACCCGCGCGTCTCCATCTACAGCACGCGCCGCCCGGTGTTGGCGCGCACCCACGTCCAGGGCCGCGTCTACAACTTCCTCGAGCGTCCCACCGGCTGGAAATGCTTCGTTTACCACTTCGCCGTCTTCCTCATCGTCCTGGTCTGCCTCATCTTCAGCGTGCTGTCCACCATCGAGCAGTATGCCGCCCTGGCCACGGGGACTCTCTTCTGGATGGTCACAGTCACCACCATCGGCTATGGGGACAAGGTGCCCCAGACGTGGGTCGGGAAG**ACCATCGCCTCCTGCTTCTCTGTCTTTGCCATCTCCTTCTTTGCGCTCCCAGCGACCGCATGGAGGTGC**TATGCTGCCGAGAACCCCGACTCCTCCACCTGGAAGATCTACATCCGGAAGGCCCCCCGGAGCCACACTCTGCTGTCACCCAGCCCCAAACCCAAGAAGTCTGTGGTGGTAAAGAA**AAAAAAG**TTCAAGCTGGACAAAGACAATGGGGTGACTCCTGGAGAGAAGATGCTCACAGTCCCCCATATCACGTGCGACCCCCCAGAAGAGCGGCGGCTGGACCACccGGAGGAGGAGGATCT**GCTACTAATTTTTCACTTCTTAAGCAAGCCGGGGATGTCGAAGAAAATCCGGGACCAATGGCCACAACCATGACGGCCCTGACAGAAGGTGCGAAGCTGTTCGAGAAGGAGATTCCCTATATCACAGAATTGGAGGGGGATGTAGAGGGTATGAAGTTTATCATCAAAGGCGAAGGGACAGGGGATGCAACAACTGGAACAATTAAGGCTAAGTACATTTGCACGACCGGCGACGTCCCGGTGCCCTGGTCCACGCTCGTCACCACGCTCACGTACGGAGCCCAGTGCTTTGCCAAATATGGCCCTGAACTTAAAGACTTCTACAAGTCGTGTATGCCGGAGGGATACGTGCAAGAGAGGACGATCACCTTTGAAGGTGACGGAGTATTCAAAACAAGAGCGGAGGTGACGTTCGAGAATGGATCGGTCTATAACCGGGTCAAGCTCAACGGACAGGGCTTTAAGAAAGATGGACACGTCCTTGGGAAGAATTTGGAGTTCAATTTCACCCCGCATTGTCTTTACATCTGGGGTGATCAGGCGAATCACGGGTTGAAATCAGCGTTCAAGATCATGCACGAGATTACGGGGAGCAAAGAGGACTTTATCGTGGCAGACCACACTCAGATGAACACTCCAATCGGAGGGGGTCCCGTACACGTACCCGAGTATCATCACCTGACCGTCTGGACATCGTTTGGAAAAGACCCTGACGACGATGAAACTGATCATCTCAACATTGTGGAAGTGATCAAGGCGGTGGACTTGGAAACATACCGGTGA**

**49. 3ʹ Ter Control, KCNQ1 Alt-Splice Cellular Reporter (L=39)**

ATGGCCGCGGCCTCCTCCCCGCCCAGGGCCGAGAGGAAGCGCTGGGGTTGGGGCCGCCTGCCAGGCGCCCGGCGGGGCAGCGCGGGCCTGGCCAAGAAGTGCCCCTTCTCGCTGGAGCTGGCGGAGGGCGGCCCGGCGGGCGGCGCGCTCTACGCGCCCATCGCGCCCGGCGCCCCAGGTCCCGCGCCCCCTGCGTCCCCGGCCGCGCCCGCCGCGCCCCCAGTTGCCTCCGACCTTGGCCCGCGGCCGCCGGTGAGCCTAGACCCGCGCGTCTCCATCTACAGCACGCGCCGCCCGGTGTTGGCGCGCACCCACGTCCAGGGCCGCGTCTACAACTTCCTCGAGCGTCCCACCGGCTGGAAATGCTTCGTTTACCACTTCGCCGTCTTCCTCATCGTCCTGGTCTGCCTCATCTTCAGCGTGCTGTCCACCATCGAGCAGTATGCCGCCCTGGCCACGGGGACTCTCTTCTGGATGGTCACAGTCACCACCATCGGCTATGGGGACAAGGTGCCCCAGACGTGGGTCGGGAAG**ACCATCGCCTCCTGCTTCTCTGTCTTTGCCATCTCCTTCTTTGCGCTCCCAGCGACCGCATGGAGGTGC**TATGCTGCCGAGAACCCCGACTCCTCCACCTGGAAGATCTACATCCGGAAGGCCCCCCGGAGCCACACTCTGCTGTCACCCAGCCCCAAACCCAAGAAGTCTGTGGTGGTAAAGAA**AAAAAAG**TTCAAGCTGGACAAAGACAATGGGGTGACTCCTGGAGAGAAGATGCTCACAGTCCCCCATATCACGTGCGACCCCCCAGAAGAgcggcggctgga**taa**ccacccggagGAGGAGGATCT**GCTACTAATTTTTCACTTCTTAAGCAAGCCGGGGATGTCGAAGAAAATCCGGGACCAATGGCCACAACCATGACGGCCCTGACAGAAGGTGCGAAGCTGTTCGAGAAGGAGATTCCCTATATCACAGAATTGGAGGGGGATGTAGAGGGTATGAAGTTTATCATCAAAGGCGAAGGGACAGGGGATGCAACAACTGGAACAATTAAGGCTAAGTACATTTGCACGACCGGCGACGTCCCGGTGCCCTGGTCCACGCTCGTCACCACGCTCACGTACGGAGCCCAGTGCTTTGCCAAATATGGCCCTGAACTTAAAGACTTCTACAAGTCGTGTATGCCGGAGGGATACGTGCAAGAGAGGACGATCACCTTTGAAGGTGACGGAGTATTCAAAACAAGAGCGGAGGTGACGTTCGAGAATGGATCGGTCTATAACCGGGTCAAGCTCAACGGACAGGGCTTTAAGAAAGATGGACACGTCCTTGGGAAGAATTTGGAGTTCAATTTCACCCCGCATTGTCTTTACATCTGGGGTGATCAGGCGAATCACGGGTTGAAATCAGCGTTCAAGATCATGCACGAGATTACGGGGAGCAAAGAGGACTTTATCGTGGCAGACCACACTCAGATGAACACTCCAATCGGAGGGGGTCCCGTACACGTACCCGAGTATCATCACCTGACCGTCTGGACATCGTTTGGAAAAGACCCTGACGACGATGAAACTGATCATCTCAACATTGTGGAAGTGATCAAGGCGGTGGACTTGGAAACATACCGGTGA**

**50. 5ʹTer Control, KCNQ1 Alt-Splice Cellular Reporter (L=39)**

ATGGCCGCGGCCTCCTCCCCGCCCAGGGCCGAGAGGAAGCGCTGGGGTTGGGGCCGCCTGCCAGGCGCCCGGCGGGGCAGCGCGGGCCTGGCCAAGAAGTGCCCCTTCTCGCTGGAGCTGGCGGAGGGCGGCCCGGCGGGCGGCGCGCTCTACGCGCCCATCGCGCCCGGCGCCCCAGGTCCCGCGCCCCCTGCGTCCCCGGCCGCGCCCGCCGCGCCCCCAGTTGCCTCCGACCTTGGCCCGCGGCCGCCGGTGAGCCTAGACCCGCGCGTCTCCATCTACAGCACGCGCCGCCCGGTGTTGGCGCGCACCCACGTCCAGGGCCGCGTCTACAACTTCCTCGAGCGTCCCACCGGCTGGAAATGCTTCGTTTACCACTTCGCCGTCTTCCTCATCGTCCTGGTCTGCCTCATCTTCAGCGTGCTGTCCACCATCGAGCAGTATGCCGCCCTGGCCACGGGGACTCTCTTCTGGATGGTCACAGTCACCACCATCGGCTATGGGGACAAGGTGCCCCAGACGTGGGTCgggaag**accatcgcctaatcctgcttctctgtcTTTGCCATCTCCTTCTTTGCGCTCCCAGCGACCGCATGGAGGTGC**TATGCTGCCGAGAACCCCGACTCCTCCACCTGGAAGATCTACATCCGGAAGGCCCCCCGGAGCCACACTCTGCTGTCACCCAGCCCCAAACCCAAGAAGTCTGTGGTGGTAAAGAA**AAAAAAG**TTCAAGCTGGACAAAGACAATGGGGTGACTCCTGGAGAGAAGATGCTCACAGTCCCCCATATCACGTGCGACCCCCCAGAAGAGCGGCGGCTGGACCACccGGAGGAGGAGGATCT**GCTACTAATTTTTCACTTCTTAAGCAAGCCGGGGATGTCGAAGAAAATCCGGGACCAATGGCCACAACCATGACGGCCCTGACAGAAGGTGCGAAGCTGTTCGAGAAGGAGATTCCCTATATCACAGAATTGGAGGGGGATGTAGAGGGTATGAAGTTTATCATCAAAGGCGAAGGGACAGGGGATGCAACAACTGGAACAATTAAGGCTAAGTACATTTGCACGACCGGCGACGTCCCGGTGCCCTGGTCCACGCTCGTCACCACGCTCACGTACGGAGCCCAGTGCTTTGCCAAATATGGCCCTGAACTTAAAGACTTCTACAAGTCGTGTATGCCGGAGGGATACGTGCAAGAGAGGACGATCACCTTTGAAGGTGACGGAGTATTCAAAACAAGAGCGGAGGTGACGTTCGAGAATGGATCGGTCTATAACCGGGTCAAGCTCAACGGACAGGGCTTTAAGAAAGATGGACACGTCCTTGGGAAGAATTTGGAGTTCAATTTCACCCCGCATTGTCTTTACATCTGGGGTGATCAGGCGAATCACGGGTTGAAATCAGCGTTCAAGATCATGCACGAGATTACGGGGAGCAAAGAGGACTTTATCGTGGCAGACCACACTCAGATGAACACTCCAATCGGAGGGGGTCCCGTACACGTACCCGAGTATCATCACCTGACCGTCTGGACATCGTTTGGAAAAGACCCTGACGACGATGAAACTGATCATCTCAACATTGTGGAAGTGATCAAGGCGGTGGACTTGGAAACATACCGGTGA**

**51. SS_mut_ Control, KCNQ1 Alt-Splice Cellular Reporter (L=39)**

ATGGCCGCGGCCTCCTCCCCGCCCAGGGCCGAGAGGAAGCGCTGGGGTTGGGGCCGCCTGCCAGGCGCCCGGCGGGGCAGCGCGGGCCTGGCCAAGAAGTGCCCCTTCTCGCTGGAGCTGGCGGAGGGCGGCCCGGCGGGCGGCGCGCTCTACGCGCCCATCGCGCCCGGCGCCCCAGGTCCCGCGCCCCCTGCGTCCCCGGCCGCGCCCGCCGCGCCCCCAGTTGCCTCCGACCTTGGCCCGCGGCCGCCGGTGAGCCTAGACCCGCGCGTCTCCATCTACAGCACGCGCCGCCCGGTGTTGGCGCGCACCCACGTCCAGGGCCGCGTCTACAACTTCCTCGAGCGTCCCACCGGCTGGAAATGCTTCGTTTACCACTTCGCCGTCTTCCTCATCGTCCTGGTCTGCCTCATCTTCAGCGTGCTGTCCACCATCGAGCAGTATGCCGCCCTGGCCACGGGGACTCTCTTCTGGATGGTCACAGTCACCACCATCGGCTATGGGGACAAGGTGCCCCAGACGTGGGTCGGGAAG**ACCATCGCCTCCTGCTTCTCTGTCTTTGCCATCTCCTTCTTTGCGCTCCCAGCGACCGCATGGAGGTGC**TATGCTGCCGAGAACCCCGACTCCTCCACCTGGAAGATCTACATCCGGAAGGCCCCCCGGAGCCACACTCTGCTGTCACCCAGCCCCAAACCCAAGAAGTCTGTGGTGGTAAAGGC**GCGCGAG**TTCAAGCTGGACAAAGACAATGGGGTGACTCCTGGAGAGAAGATGCTCACAGTCCCCCATATCACGTGCGACCCCCCAGAAGAGCGGCGGCTGGACCACccGGAGGAGGAGGATCT**GCTACTAATTTTTCACTTCTTAAGCAAGCCGGGGATGTCGAAGAAAATCCGGGACCAATGGCCACAACCATGACGGCCCTGACAGAAGGTGCGAAGCTGTTCGAGAAGGAGATTCCCTATATCACAGAATTGGAGGGGGATGTAGAGGGTATGAAGTTTATCATCAAAGGCGAAGGGACAGGGGATGCAACAACTGGAACAATTAAGGCTAAGTACATTTGCACGACCGGCGACGTCCCGGTGCCCTGGTCCACGCTCGTCACCACGCTCACGTACGGAGCCCAGTGCTTTGCCAAATATGGCCCTGAACTTAAAGACTTCTACAAGTCGTGTATGCCGGAGGGATACGTGCAAGAGAGGACGATCACCTTTGAAGGTGACGGAGTATTCAAAACAAGAGCGGAGGTGACGTTCGAGAATGGATCGGTCTATAACCGGGTCAAGCTCAACGGACAGGGCTTTAAGAAAGATGGACACGTCCTTGGGAAGAATTTGGAGTTCAATTTCACCCCGCATTGTCTTTACATCTGGGGTGATCAGGCGAATCACGGGTTGAAATCAGCGTTCAAGATCATGCACGAGATTACGGGGAGCAAAGAGGACTTTATCGTGGCAGACCACACTCAGATGAACACTCCAATCGGAGGGGGTCCCGTACACGTACCCGAGTATCATCACCTGACCGTCTGGACATCGTTTGGAAAAGACCCTGACGACGATGAAACTGATCATCTCAACATTGTGGAAGTGATCAAGGCGGTGGACTTGGAAACATACCGGTGA**

**52. 0-Frame Control, KCNQ1 Alt-Splice Cellular Reporter (L=39)**

ATGGCCGCGGCCTCCTCCCCGCCCAGGGCCGAGAGGAAGCGCTGGGGTTGGGGCCGCCTGCCAGGCGCCCGGCGGGGCAGCGCGGGCCTGGCCAAGAAGTGCCCCTTCTCGCTGGAGCTGGCGGAGGGCGGCCCGGCGGGCGGCGCGCTCTACGCGCCCATCGCGCCCGGCGCCCCAGGTCCCGCGCCCCCTGCGTCCCCGGCCGCGCCCGCCGCGCCCCCAGTTGCCTCCGACCTTGGCCCGCGGCCGCCGGTGAGCCTAGACCCGCGCGTCTCCATCTACAGCACGCGCCGCCCGGTGTTGGCGCGCACCCACGTCCAGGGCCGCGTCTACAACTTCCTCGAGCGTCCCACCGGCTGGAAATGCTTCGTTTACCACTTCGCCGTCTTCCTCATCGTCCTGGTCTGCCTCATCTTCAGCGTGCTGTCCACCATCGAGCAGTATGCCGCCCTGGCCACGGGGACTCTCTTCTGGATGGTCACAGTCACCACCATCGGCTATGGGGACAAGGTGCCCCAGACGTGGGTCGGGAAG**ACCATCGCCTCCTGCTTCTCTGTCTTTGCCATCTCCTTCTTTGCGCTCCCAGCGACCGCATGGAGGTGC**TATGCTGCCGAGAACCCCGACTCCTCCACCTGGAAGATCTACATCCGGAAGGCCCCCCGGAGCCACACTCTGCTGTCACCCAGCCCCAAACCCAAGAAGTCTGTGGTGGTAAAGAA**AAAAAAG**TTCAAGCTGGACAAAGACAATGGGGTGACTCCTGGAGAGAAGATGCTCACAGTCCCCCATATCACGTGCGACCCCCCAGAAGAGCGGCGgctggaccaccc**c**ggaggaggaggaTCT**GCTACTAATTTTTCACTTCTTAAGCAAGCCGGGGATGTCGAAGAAAATCCGGGACCAATGGCCACAACCATGACGGCCCTGACAGAAGGTGCGAAGCTGTTCGAGAAGGAGATTCCCTATATCACAGAATTGGAGGGGGATGTAGAGGGTATGAAGTTTATCATCAAAGGCGAAGGGACAGGGGATGCAACAACTGGAACAATTAAGGCTAAGTACATTTGCACGACCGGCGACGTCCCGGTGCCCTGGTCCACGCTCGTCACCACGCTCACGTACGGAGCCCAGTGCTTTGCCAAATATGGCCCTGAACTTAAAGACTTCTACAAGTCGTGTATGCCGGAGGGATACGTGCAAGAGAGGACGATCACCTTTGAAGGTGACGGAGTATTCAAAACAAGAGCGGAGGTGACGTTCGAGAATGGATCGGTCTATAACCGGGTCAAGCTCAACGGACAGGGCTTTAAGAAAGATGGACACGTCCTTGGGAAGAATTTGGAGTTCAATTTCACCCCGCATTGTCTTTACATCTGGGGTGATCAGGCGAATCACGGGTTGAAATCAGCGTTCAAGATCATGCACGAGATTACGGGGAGCAAAGAGGACTTTATCGTGGCAGACCACACTCAGATGAACACTCCAATCGGAGGGGGTCCCGTACACGTACCCGAGTATCATCACCTGACCGTCTGGACATCGTTTGGAAAAGACCCTGACGACGATGAAACTGATCATCTCAACATTGTGGAAGTGATCAAGGCGGTGGACTTGGAAACATACCGGTGA**
